# Convergent loss of an EDS1/PAD4 signalling pathway in several plant lineages predicts new components of plant immunity and drought response

**DOI:** 10.1101/572560

**Authors:** EL Baggs, AS Thanki, R O’Grady, C Schudoma, W Haerty, KV Krasileva

## Abstract

Plant innate immunity relies on NLR receptors that recognize pathogen derived molecules and activate downstream signalling pathways. We analyzed the variation in copy number of NLR genes across flowering plants, and identified a number of species with a low number of NLRs relative to sister species. Two distinct lineages, one monocot (Lentibulariaceae) and one dicot (Alismatales) encapsulate four species with particularly few NLR genes. In these lineages, loss of NLRs coincided with loss of the well-known downstream immune signalling complex (EDS1-PAD4). When we expanded our analysis across the whole proteomes, we were able to identify other characterized immune genes absent only in Lentibulariaceae and Alismatales. Additionally, we identified a small subset of genes with unknown function convergently lost in all four species. We predicted that some of these genes may have a role in plant immunity. Gene expression analyses confirmed that a group of these genes was differentially expressed under pathogen infection. Another subset of these genes was differentially expressed upon drought providing further evidence of a link between the drought and plant immunity.

## Introduction

Plants evolved from a common ancestor with charophyte green algae upon a major change in lifestyle, the transition from water to land, over 450 million years ago (MYA) (Sanderson et al. 2004; Zhong, Sun, and Penny 2015). Extant plant lineages, such as bryophytes (mosses, liverworts and hornworts), terrestrial vascular plants (ferns), gymnosperms (pine, spruce) and angiosperms (monocots, dicots) diverged from the ancestor of terrestrial plants over 300 MYA (Zeng et al. 2014). Crops in modern agriculture typically belong to the monocot or dicot angiosperm classes. They are continuously exposed to both biotic and abiotic stresses, which can result in major yield losses. These losses will be exacerbated as a consequence of climate change (Ramegowda and Senthil-Kumar 2015). There is an urgent need to further our understanding of the mechanisms of stress responses in wild relatives of crop plants as such knowledge can facilitate the breeding of more stress resilient crops.

Land plant evolution has likely always been accompanied by the presence of microbes, including some whose interaction would be antagonistic to plant fitness and therefore deemed pathogenic. The initial plant disease resistance response to a pathogen relies on recognition of extracellular microbe associated molecular patterns (MAMPs) and is termed MAMP-triggered immunity (MTI). Most pathogens deploy effectors that can suppress MTI to facilitate virulence. Hence, a second intracellular monitoring system of effector triggered immunity (ETI) is essential for resistance to many pathogens. Plant nucleotide binding–leucine-rich repeat (NLR) immune receptors mediate ETI upon detection of intracellular pathogen molecules. The NLR immune recognition system predates land plant emergence as proteins with a similar architecture are present in Charophyta and red algae (Gao, 2018). The NLR proteins are typically composed of three or more domains (Baggs, Dagdas, and Krasileva 2017; Jones, Vance, and Dangl 2016). The Nucleotide Binding (NB-ARC) domain is the common central component of NLRs and is involved in receptor activation, similar to the NACHT domain found in animal NLR immune receptors (Jones, Vance, and Dangl 2016). The NB-ARC domain is commonly followed by a series of Leucine Rich Repeats (LRRs), previously shown to mediate intramolecular interactions within NLRs and intermolecular binding of effector proteins (Dodds, Lawrence, and Ellis 2001; Catanzariti et al. 2010; Krasileva, Dahlbeck, and Staskawicz 2010). A Toll-like, Interleukin-1 (TIR-1) or a coiled-coil (CC) domain, are typically found at the N-terminus, where they function in the initiation of the signalling cascade (Bernoux et al. 2011). NLRs containing a TIR-1 domain are referred to as TNLs, while CNL refers to NLRs with a CC domain. A third clade of NLRs called RNLs is characterized by the presence of an RPW8 domain. Members of all three clades, CNLs, TNLs and RNLs are present in basal angiosperms, such as Amborella, and gymnosperms (Van Ghelder et al. 2019; Gao et al. 2018).

A subset of NLRs are not involved in the perception of effector induced changes but instead perpetuate signalling downstream of effector sensing NLRs (Le Roux et al. 2015; Ortiz et al. 2017; Sarris et al. 2016; Saucet et al. 2015). All TNLs characterized to date depend on downstream activity of members in the RNL clade, such as *N REQUIREMENT GENE 1* (NRG1) and *ACTIVATED DISEASE RESISTANCE-LIKE1* (ADR1) (Lapin et al. 2019; Castel et al. 2018; Qi et al. 2018; Z. Wu et al. 2019). Within TNLs and CNLs, there are additional helper NLR sub-clades, which help to activate effector sensing NLRs and sometimes form sensor-helper networks in which multiple sensors interact with the same helper (Narusaka et al. 2014; C.-H. Wu et al. 2017; C.-H. Wu, Derevnina, and Kamoun 2018).

Improvement of crop varieties to resist new and evolving pathogen threats often utilises introgression ofNLR genes to convey new pathogen recognition specificity by taking advantage of conserved signalling pathways. Despite 120-180 MYA of independent monocot and dicot evolution, some NLRs such as barley *MILDEW LOCUS A (MLA)* are functional when transformed into Arabidopsis *(Maekawa et al. 2012)*. This indicates overall conservation of downstream signalling pathways across monocots and dicots. However, NLR subclasses present in monocots and dicots differ. Among dicots, typically over half the NLRs contain a TIR-1 domain (Sarris et al. 2016; Meyers et al. 2003). In contrast, no TIR-1 domain containing NLR receptors have been identified in monocots to date.

The two key signalling components that are downstream of TIR-1 NLRs, ENHANCED DISEASE SUSCEPTIBILITY 1 (EDS1) and PHYTOALEXIN DEFICIENT 4 (PAD4), are present in almost all angiosperms despite the absence of TIR-1 NLRs in monocots (Lapin et al. 2019; Bhandari et al. 2019; Wagner et al. 2013). Although wheat has no TIR-1 NLRs, EDS1 has retained functional importance in immunity in wheat, since overexpression of TaEDS1 in a susceptible wheat cultivar results in reduced *Blumeria graminis f.sp. tritici* haustorial growth (Chen et al. 2018). Moreover, wheat *EDS1* was able to complement the *eds1* mutant in Arabidopsis (Chen et al. 2018), indicating a highly conserved role in immune signalling. EDS1 in dicots forms a complex with PAD4 and also SENESCENCE ASSOCIATED GENE 101 (SAG101) (Wagner et al. 2013). *In planta* work in Arabidopsis showed that EDS1 binds directly to PAD4 and SAG101 to form mutually exclusive heterodimeric complexes (Wagner et al. 2013), with the subcellular localization of EDS1 depending on the interacting partner (Zhu et al. 2011). Recent work has also demonstrated molecular and genetic evidence that Arabidopsis EDS1, SAG101 and NRG1 interact in a complex upon TNL activation, resulting in cell death (Lapin et al. 2019). Among angiosperms, *EDS1* and *PAD4* are typically conserved, whilst the *SAG101* gene is absent from available genomes of grasses (Wagner et al. 2013). TNL signalling involves enzymatic catalysis of NAD+ which activates EDS1 through a yet unknown mechanism (Wan et al. 2019; Horsefield et al. 2019). RNLs act downstream of EDS1/PAD4/SAG101 and are involved in the induction of cell death (Lapin et al. 2019; Castel et al. 2018; Qi et al. 2018; Z. Wu et al. 2019).

Another signalling component that is often vital in plant immunity, but whose function remains elusive, is NDR1. First discovered in Arabidopsis, NDR1 has since been shown to be conserved across dicots and required for signalling of several CNLs (Century, Holub, and Staskawicz 1995; Coppinger et al. 2004). NDR1 is thought to mediate resistance by controlling fluid loss in the cell (Knepper, Savory, and Day 2011). Both NDR1 and EDS1/PAD4/SAG101 mediated signalling converge in triggering increase in SA (H. Cui et al. 2018; Venugopal et al. 2009). Conserved signalling components of plant immunity have been utilized to engineer broad spectrum resistance across monocots and dicots (Cao, Li, and Dong 1998; Xu et al. 2017), demonstrating the translational impacts on crop production from understanding evolution of immune signalling.

Within flowering plants, there is a huge amount of genetic, genomic and phenotypic diversity that can be mined to understand plant molecular pathways. Previous studies looking at convergent evolution have successfully predicted gains and losses of genetic pathways and identified new pathway components (Ibarra-Laclette et al. 2013; Michael et al. 2017; Olsen et al. 2016; W. Wang et al. 2014). In the study of plant symbiotic interactions this approach has been particularly fruitful, identifying many proteins which have been functionally validated in facilitating arbuscular mycorrhization of plants (Griesmann et al. 2018; Bravo et al. 2016; Radhakrishnan et al. 2019) and in uncovering the origin of nitrogen-fixing rhizobium symbioses (Griesmann et al. 2018; van Velzen et al. 2018). The growing number of available plant genomes from species with divergent environments increases the opportunity for elucidation of pathways that facilitate acclimation to their new environments.

In this study, we used comparative genomics to look at the NLR copy number variation across angiosperms. We identified independent contractions in NLR number and diversity among monocots and dicots and discovered that EDS1/PAD4 pathways have been convergently lost in at least five species. We further analysed components of disease resistance pathways and used gene family clustering methods to identify new genes following the same evolutionary pattern. Our analyses predicted new candidates involved in the EDS1/PAD4 signalling pathway and provide links between plant immunity and drought response.

## Results

### Several lineages of flowering plants have lost most of the NLR plant immune receptor clades present in the common ancestor of flowering plants

The NLR gene family is complex, with copy numbers variable by 10 fold between species of the same family (Supplemental Figure 1, Supplemental Table 1) (Baggs, Dagdas, and Krasileva 2017). The variation in intraspecific and interspecific copy number of NLR*s* is becoming more apparent with the increasing number of available genomes. The historical bias in genome sequencing in favour of economically important or model species means most studies focus on the *Brassicaceae*, *Solanaceae* and *Poaceae* families (Van de Weyer et al. 2019; Stam, Silva-Arias, and Tellier 2019; Sarris et al. 2016). We surveyed NLR copy number variation in 95 publicly available angiosperm genomes spanning 24 orders (Figure 1, Supplemental Figure 1, Supplemental Table 1). While most of the sequenced plant genomes contain between 200-500 NLRs (Figure 1A), there have been extreme losses and expansions in the NLR gene family among both monocots and dicots. NLR copy numbers in the Poaceae range from 33 in *Oropetium thomaeum* to 1,157 in wheat (*Triticum aestivum*) (Supplemental Figure 1, Supplemental Table 1). We decided to further investigate plant genomes that have convergently lost the majority of NLRs.

**Fig 1.**
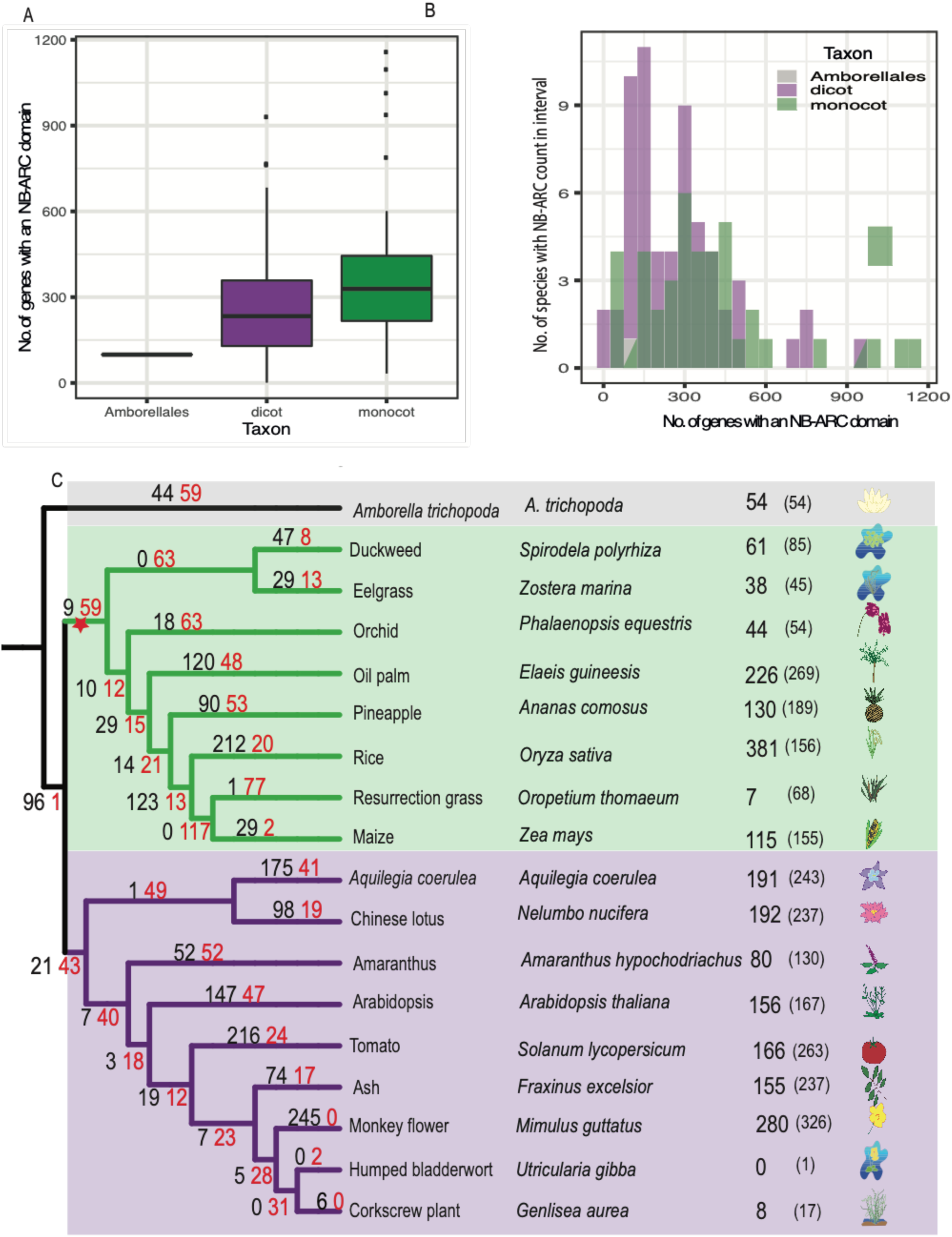
Phylogenetic relationship and NLR repertoires of the plant species used in this study. Phylogenetic relationship and NLR repertoires of the plant species used in this study (A) Boxplot showing the variation in number of NB-ARC domains across monocot and dicot genomes available on Phytozome, EnsemblePlants or CoGe. (B) Histogram of no. of NB-ARC domains identified in a vailable genomes. (C) A species tree of monocot and dicot genomes of interest. Number of NLRs with all 6 characteristic NLR amino-acid motifs annotated in each species displayed in line with leaf tips with number of NLRs identified by PfamScan and the plant_rgenes pipeline in brackets. Black numbers on branches indicate number of NLRs gained and red numbers refer to NLRs lost. A red star indicates loss of SAG101 and TIR1-NLRs.

For further analyses, we selected 18 species to represent a broad range of families, sister species with highly divergent NLR copy number and high-quality genome assembly (Figure 1B). We considered an NLR number to be low if it was below the 1st quartile: 217 NLRs for monocots (eelgrass (*Zostera marina),* duckweed (*Spirodela polyrhiza),* orchid (*Phalaenopsis equestris),* pineapple (*Ananas comosus),* resurrection grass (*Oropetium thomaeum),* maize *(Zea mays)*) and 129 NLRs for dicot species (humped bladderwort (*Utricularia gibba)* and corkscrew plant (*Genlisea aurea)*). To test if the reduction of NLR numbers is a consequence of incomplete annotations (e.g. as a result of low genome assembly contiguity), we mined the genomic sequences of these plants using NLRannotator. This software predicts NLR motifs at the DNA level and is therefore independent of protein annotation (Steuernagel et al. 2015, 2018) (Supplemental Table 2). NLRannotator results confirmed the reduced NLR numbers in eelgrass, duckweed, orchid, resurrection grass, maize, pineapple, humped bladderwort and corkscrew plant. The results show the low NLR number at the proteome level is consistent with genome wide prediction of NLRs.

When investigating plant genomes with a low number of NLR genes, we identified two groups of species, those which retained all major NLR sub-classes (CNL, TNL, RNL) and those which did not. In order to identify the presence and absence variation of TNL, CNL and RNL, we built a maximum likelihood phylogeny (Figure 2) on the NB-ARC domain of all NLRs retaining 6 key functional motifs (Walker A, RNBS-A, WALKER-B, RNBS-C, GLPL and RNBS-D) (Wen et al. 2017; G.-F. Wang et al. 2015).

**Fig 2.**
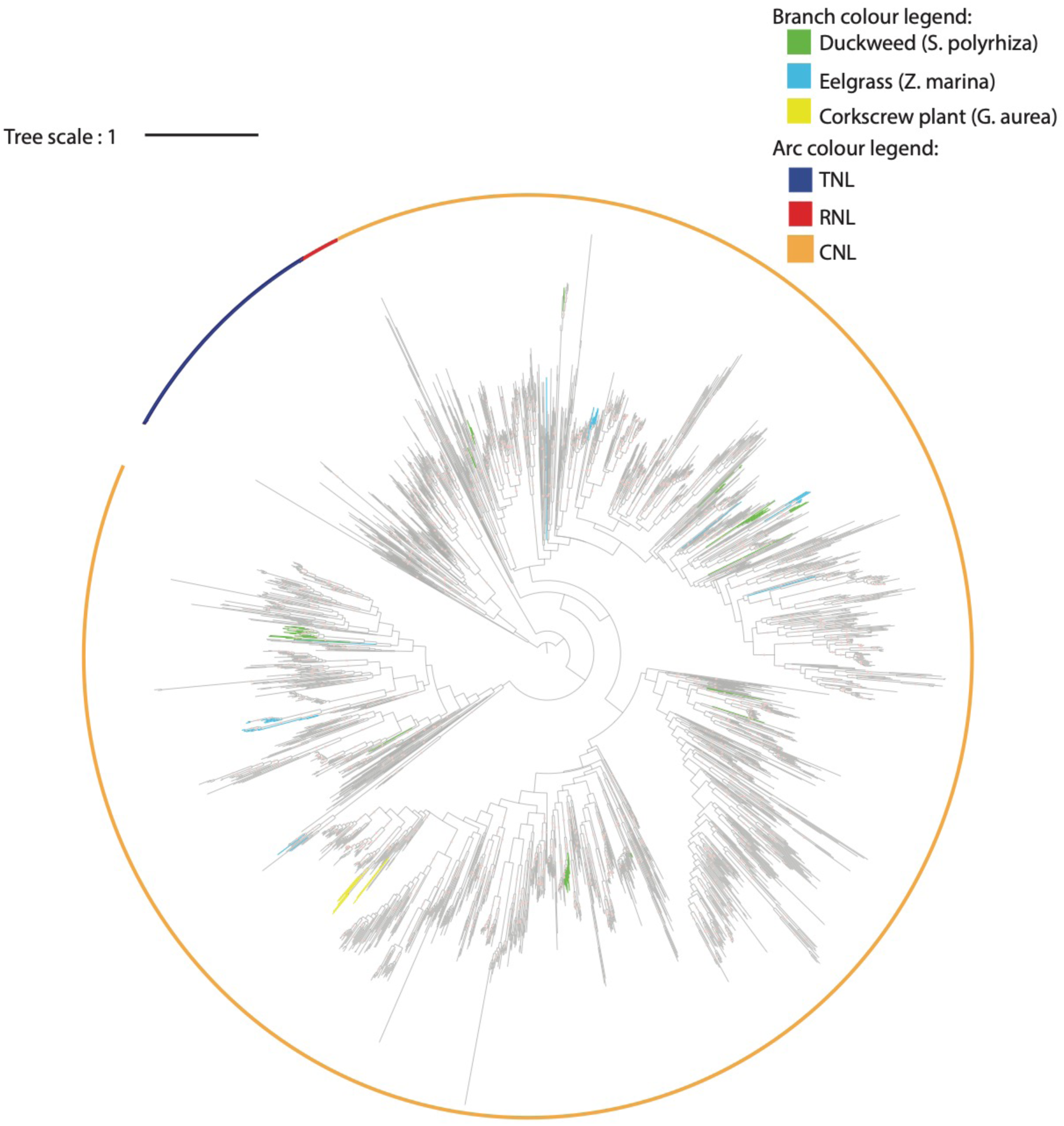
Maximum likelihood phylogeny of NLRs in the 18 representative plant species and selected reference NLRs. The maximum likelihood tree is based on the alignment of NB-ARC domain for the 18 representative species of Amborella, eelgrass, duckweed, oil palm, pineapple orchid, resurrection grass, maize, rice, arabidopsis, *A. coerulea*, lotus, amaranthus, tomato, ash, monkey flower, humped bladderwort and corkscrew plant. Bootstraps >80 are indicated by a red dot, branch colours denote species. Clades as defined by bootstrap >80, orange arc represents CNLs, blue for TNLs and red for RNLs.

We applied tree reconciliation between the species tree and NLR gene tree to quantify NLR gains and losses across the phylogeny (Figure 1C). We observed a high turnover of NLRs across the phylogeny, with large expansions in specific lineages (*O. sativa, E. guineensis, A. coerula, A. thaliana, S. lycopersicum, E. guttata*), as well as extensive losses such as in the ancestral lineage of *Z. mays* and *O. thomaeum*. The NLR gene tree was consistent with the loss of TNLs in monocot species and monkey flower as previously reported (J. Kim et al. 2012; Sarris et al. 2016). We identified the absence of TNLs in two further dicot species (humped bladderwort, corkscrew plant). The RNL subclass NRG1 is genetically required for signalling of many TNLs and among dicots there is co-occurrence of NRG1 and TNLs (Castel et al. 2018; Lapin et al. 2019; Z. Wu et al. 2019; Qi et al. 2018). We established the absence of the NRG1 type RNLs in all monocots in our analysis and in those dicots that have also lost TNLs: monkey flower (consistent with (J. Kim et al. 2012)), humped bladderwort and corkscrew plant (Supplemental Figure 2). In addition, no NRG1 ortholog could be identified through phylogeny or reciprocal BLAST in *A. trichopoda* and *A. hypochondriacus* which have 14 and 2 TNLs, respectively. Of note is that 10 *A. trichopoda* and 1 *A. hypochondriacus* TNLs belong to a single clade (bootstrap = 65), suggesting that they might function independently of NRG1. To verify that RNLs were not removed due to strict alignment criteria or misannotation, we performed a modified reciprocal blast which confirmed that no RNLs could be identified in duckweed, eelgrass, humped bladderwort or corkscrew plant. Conversely, RNLs of the ADR1 subclass were identified in amborella and rice for whom the RNLs were absent in the phylogeny. The presence of the RNL clade in monocots, dicots and amborella suggests convergent clade specific loss of RNLs in duckweed, eelgrass, humped bladderwort and corkscrew plant.

Despite the loss of many NLRs, as shown by our gain and loss analysis (Figure 1C) those retained in duckweed, eelgrass and humped bladderwort have undergone recent lineage specific expansions (Figure 2). In duckweed, eelgrass and humped bladderwort 88%, 84% and 100% of NLRs respectively are present in a monophyletic species-specific clade with bootstrap support >80. Several duckweed and eelgrass NLRs arose from recent tandem duplications as can be seen from their consecutive gene identifiers (Figure 2). This suggests that despite overall gene loss, both species still require the remaining NLR clades.

To test if a general reductin in large protein families was responsible for low NLR numbers in plants that have lost NLR sub-classes (duckweed, eelgrass, humped bladderwort and corkscrew plant), we annotated receptors belonging to the Receptor-Like Kinase family (RLKs), a MTI immune receptor family and a family of actin proteins which is less variable in copy number than immune genes. The reduction of RLKs compared to sister lineages was not as pronounced as with NLRs, with the exception of corkscrew plant (Supplemental Table 2, 3). For the actin gene family, the percentage of actin encoding genes in the proteome was similar between RNL absent species and sister species (Supplemental Table 3). This suggests that the reduction in NLR copy number and the loss of subclasses can not be explained by the general contraction of large protein families.

### Independent loss of immune signalling pathway components *EDS1/PAD4/SAG101* among angiosperm lineages

The co-occurence of TNL and RNL loss prompted us to investigate if the species without these NLR subclasses have also lost any other immune signalling components. This led us to identify the absence of EDS1/PAD4/SAG101 and NDR1 among species that lost RNLs and TNLs. In total, we surveyed for the presence of 15 known immune signalling components using reciprocal blastp followed by tblastn to remove false negatives due to incomplete annotation (Figure 3 and Supplemental Figure 3, Supplemental Table 4, 5). Of the 15 immune genes characterized in Arabidopsis, 5 including *RPM1 INTERACTING PROTEIN 4 (RIN4)*, *MAP KINASE 4*, *ISOCHORISMATE SYNTHASE 1 (ICS1)*, *METACASPASE 4/ 5* and *REQUIRED FOR MLA12 RESISTANCE 1 (RAR1),* were conserved across all 18 monocots and dicots (S4 Table). The presence of salicylic acid (SA) biosynthetic gene *ICS1* and SA receptor *NPR* gene family members in all species, except humped bladderwort, suggests the SA biosynthesis and signaling remains intact among most species. Thus, it appears that the SA pathway is retained independently of TNLs and RNLs that utilise SA for signaling.

**Fig 3.**
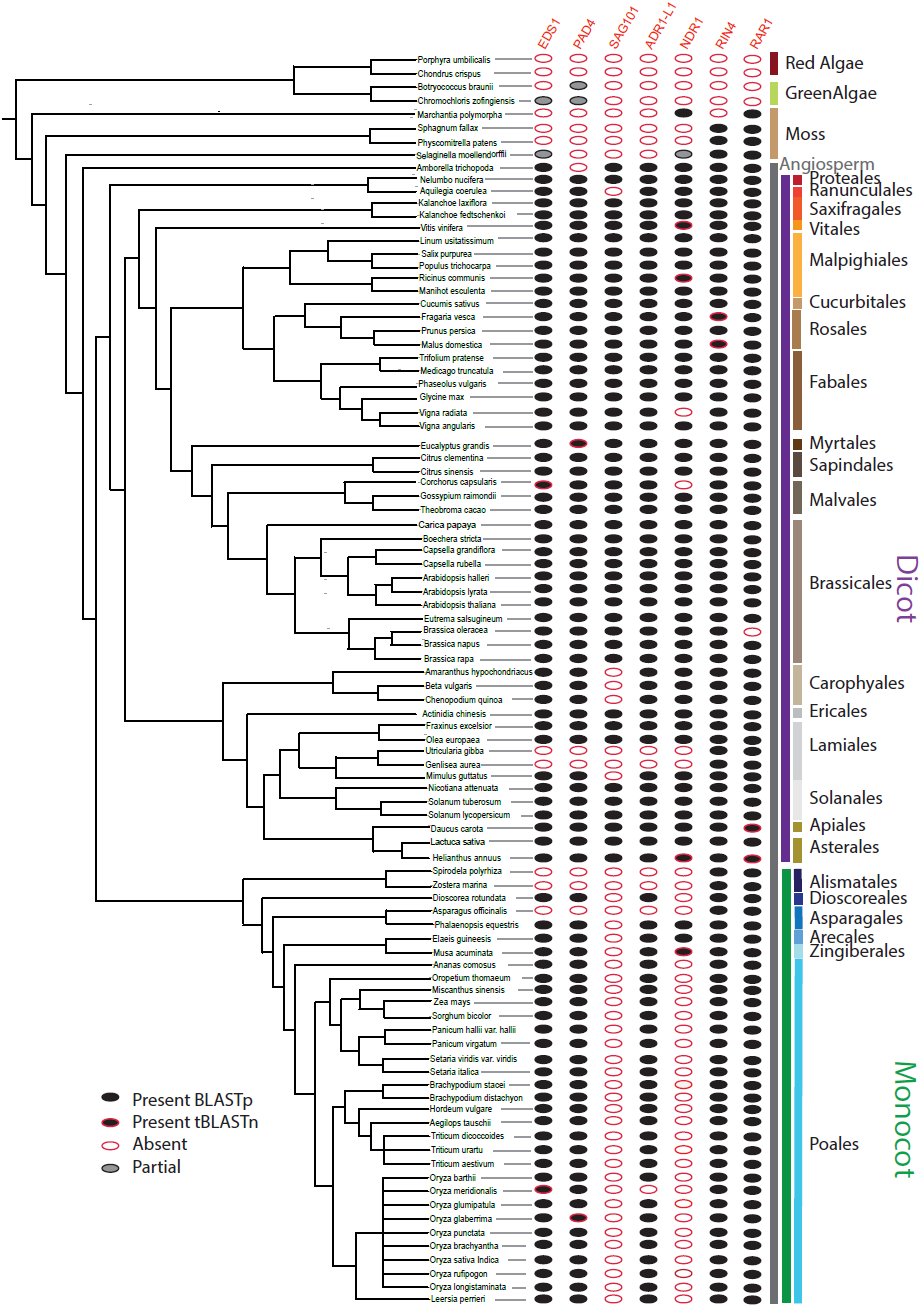
Presence/absence analysis of known plant immunity components. Rows denote species, which are arranged as per phylogenetic relationship, with the green bar indicating monocots and purple bar for dicots. Gene names are listed at the top. Circles in columns denote the presence or absence of known components of the NLR signalling pathway. Black filled circles - ortholog identified by reciprocal blastp and supported by tblastn indicated by red outline, grey circles-partial ortholog and white circles with grey outline if no ortholog could be identified. Orthology was also manually curated using EnsemblePlant gene trees or Phytozome synteny where available.

The loss TNL and RNL receptors coincides with the loss of SAG101 (Wagner et al. 2013). All monocots in our analyses as well as *A. coerulea* and *E. guttata* appear to have lost *SAG101 (Wagner et al. 2013)*. In addition to *SAG101*, *EDS1, PAD4* and *NDR1* were absent in the same species that lacked RNLs (Figure 3). NDR1 similar to SAG101 is also absent in all Poales. To confirm that the inferred absence of *EDS1, PAD4*, *SAG101* and *NDR1* is not an annotation artifact in duckweed, eelgrass, humped bladderwort and corkscrew plant, we scanned the respective genomes using tblastn and HMMER searches for the indicative lipase 3 motif. Both analyses supported the absence of EDS1, PAD4 and SAG101 (Supplemental Figure 4, https://github.com/krasileva-group/Aquatic_NLR/tree/master/). RNLs, EDS1 and PAD4 are known to interact in a protein complex and our data suggests that they also form an evolutionary unit.

To identify if the loss of *EDS1*, *PAD4*, *SAG101* and RNLs was common among plants we repeated the reciprocal blastp followed by tblastn on 95 available genomes used in our initial survey of NLRs (Figure 3, Supplemental Table 5). Orthology was then manually curated using EnsemblPlant gene trees or Phytozome synteny, where available. This identified *Asparagus officinalis* as the only other angiosperm with a sequenced genome that is missing EDS1, PAD4, SAG101, NDR1 and RNLs. The loss of *EDS1*, *PAD4*, *SAG101* and RNLs appears to be limited to species with low NLR number and in the absence of NLR subclasses, RNL and TNL.

To investigate if the EDS1, PAD4, RNLs and NDR1 were present in basal monocots, we downloaded an orthogonal dataset of the 1,000 plant transcriptomes (Wickett et al. 2014; Matasci et al. 2014; One Thousand Plant Transcriptomes Initiative 2019). We then queried transcriptomes of species within the earliest branching angiosperm orders of Magnoliales, Piperales and Laurales (Supplemental Figure 5, Supplemental Table 6). With the exception of *Peperomia fraseri,* species within these orders had transcripts that were orthologous to at least three of the signalling genes expected to be present in monocots EDS1, PAD4, RNLs and NDR1 (Supplemental Figure 6, Supplemental Table 7). Generally, EDS1, PAD4, RNLs and NDR1 transcripts were present in the majority of Pandales, Dioscoreales, Acorales species with available transcriptomes. This supports the notion that the EDS1, PAD4, RNLs and NDR1 proteins were present in the most recent common ancestor of monocots.

Recent research supports that gene loss events are less likely to be annotation artifacts if they are clade-specific loss events rather than being absent from a single species (Deutekom et al. 2019). In the absence of more available genomes, we therefore looked at the orthogonal dataset of available transcriptomes. Among the Lamiales (humped bladderwort, corkscrew plant), only 4/48 transcriptomes did not have transcripts corresponding to EDS1, PAD4, SAG101, RNLs and NDR1. Three of these species were members of the Lentibulariaceae family (humped bladderwort, corkscrew plant). On the other hand, transcripts for EDS1, PAD4, RNLs and NDR1 were found in the transcriptomes of species from the sister family Bignoniaceae (Supplemental Fig 5, Supplemental Table 6). EDS1, PAD4, RNLs and NDR1 transcripts were absent among multiple species in the orders of Piperales, Alismatales, Asparagales and Liliales, supporting our finding of loss of these components in Alismatales (duckweed and eelgrass) and Asparagales (asparagus). Although EDS1, PAD4, RNLs and NDR1 transcripts were absent in these orders, without complete genomes we cannot exclude the possibility that these genes were not expressed at the tissue and time point sampled for the transcriptomics analysis. Duckweed and eelgrass belong to the Araceae and Zosteraceae families, respectively. Since EDS1, PAD4, RNL and NDR1 are present in the transcriptome of at least one other member of the Araceae (*Pistia stratiotes*), we concluded that these genes were lost independently in eelgrass and duckweed.

**Fig 5.**
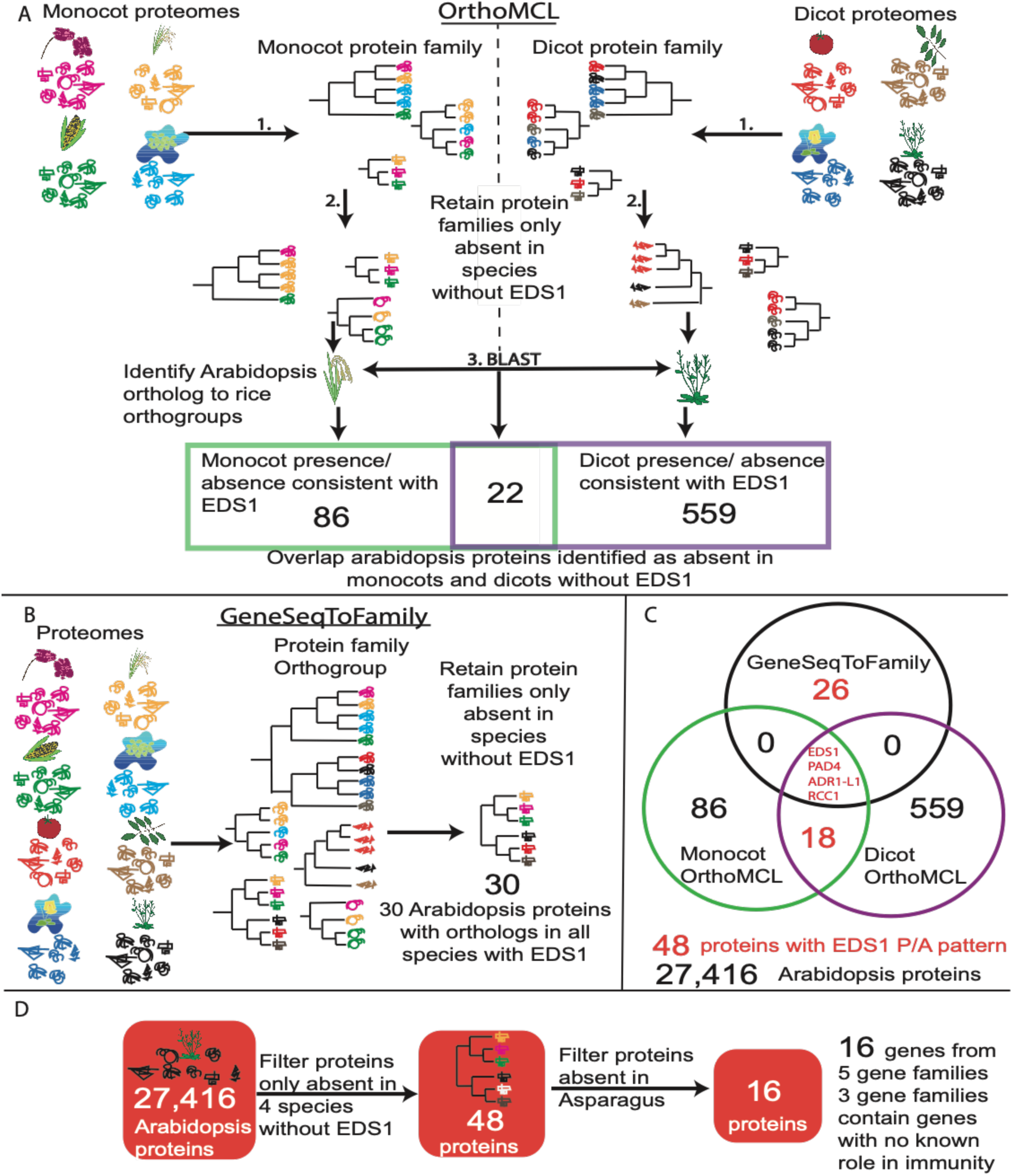
Protein family analysis to identify *AngioSperm Typically-Retained, EDS1-Lost (ASTREL)* genes. (A) Schematic of the OrthoMCL approach to cluster protein families separately among monocot and dicot species and then filtering for protein families present in terrestrial and absent in aquatic species. Proteins are denoted by different line drawings, colours of proteins represent species origin and phylogenetic trees represent gene trees for each protein. The Venn diagram at bottom provides the number of Arabidopsis proteins, which are absent in just monocot, dicot or all angiosperms without EDS1 and interacting proteins. (B) Illustration of the GeneSeqToFamily method which uses the monocot and dicot proteomes together to establish gene trees across the angiosperms. (C) A Venn diagram summarizing the results of two methods. Genes and numbers marked in red are those subsequently referred to as *ASTREL* genes. (D) Schematic of gene clustering followed by blastp and tblastn to filter ASTREL genes for presence or absence in the asparagus genome.

In order to further our understanding of the effects of the gene loss in the context of the genomic locus, we reconstructed the homologous regions surrounding EDS1, PAD4 and ADR1 between *Arabidopsis thaliana* and duckweed, eelgrass, humped bladderwort or corkscrew plant (Supplemental Figure 7). Owing to the large divergence time and the low contiguity of some of the assemblies we found little synteny conservation. Therefore, we explored the conservation of the genomic regions containing EDS1, PAD4, and ADR1 in the recent duckweed chromosome level genome assembly. We used protein orthology between pineapple and duckweed predicted with GeneSeqtoFamily (Thanki et al. 2018) and homologous gene regions identified with SynMap2 (Haug-Baltzell et al. 2017). While we were unable to find a contiguous genomic region with synteny across EDS1 (Figure 4), we could observe a relative conservation of synteny for the homologous regions of PAD4 (Figure 4) and ADR1 subclass NLRs (Figure 4) as genes neighbouring the target genes in *A. comosus* are found on the same chromosome and in close vicinity. The lack of conservation of strand orientation between orthologs suggests recombination after speciation. The reconstruction of homologous regions revealed that the region around the lost genes was assembled and the gene loss event in duckweed was due to deletions affecting single or just a few genes rather than a consequence of the loss of large regions.

**Fig 4.**
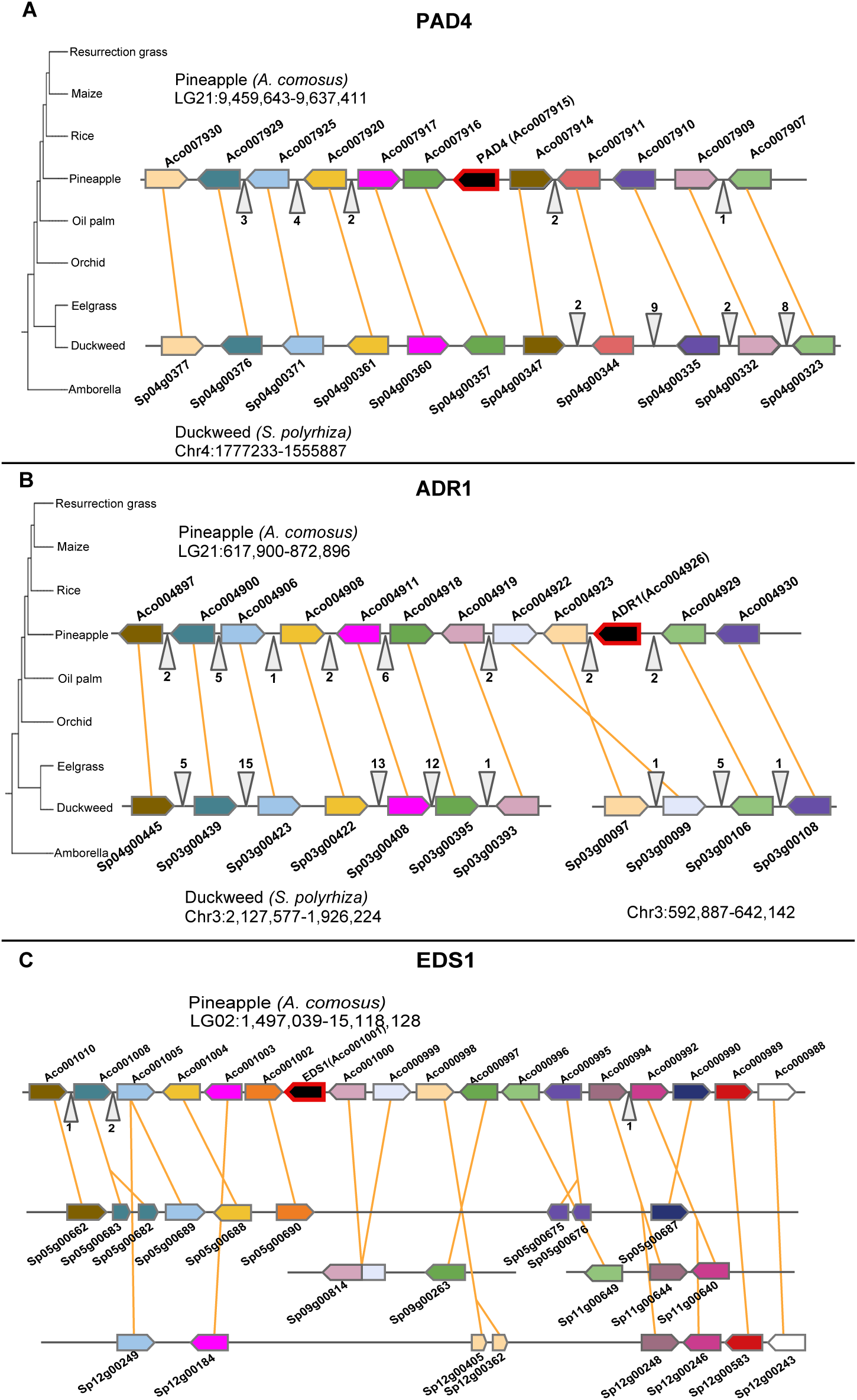
Syntenic block of ASTREL genes PAD4, EDS1 and ADR1 genomic locus between pineapple and duckweed. Boxes indicate genes direction of point on box indicates gene orientation, with orange lines connecting orthologs between pineapple and duckweed. Grey triangles indicate genes not displayed. Focal gene is highlighted with a red outline (A) Synteny plot of genes upstream and downstream of PAD4 in pineapple relative to duckweed. (B) Synteny plot of genes upstream and downstream of ADR1 in pineapple relative to duckweed. (C) Synteny plot of genes upstream and downstream of EDS1 in pineapple relative to duckweed.

### Orthogroup analysis of protein families provides a global view of the genes lost together with EDS1/PAD4

The convergent loss of *EDS1/PAD4* and *EDS1-*dependent NLRs led us to hypothesize that other, as yet unknown, components of the *EDS1*-dependent signalling cascade would also have been convergently lost in these species. To uncover novel proteins that can potentially function in conjunction with the EDS1-mediated NLR signalling cascade, we performed two analyses: OrthoMCL (L. Li, Stoeckert, and Roos 2003) and GeneSeqToFamily (Thanki et al. 2018) to cluster proteomes into orthogroups and examine gene loss patterns. We analysed the 18 plant proteomes, which we surveyed earlier for NLR sub-classes: monocots (*Z. marina, S. polyrhiza, P. equestris, D. rotundata*, *E. guineesis, O. thomaeum, O. sativa* and *Z. mays*) and dicots (*A. coerulea, N. nucifera, A. thaliana, A. hypochondriachus, S. lycopersicum, F. excelsior, E. guttata, G. aurea* and *U. gibba*) with outgroups *Selaginella moellendorffii* and *Amborella trichopoda* (Figure 5).

Through the combined OrthoMCL analyses, we identified 17 Arabidopsis genes that lacked orthologous groups only in four species that lost EDS1/PAD4/RNLs: duckweed, eelgrass and humped bladderwort but had orthologs in other 13 species. The GeneSeqToFamily analysis identified 31 Arabidopsis genes from 10 orthogroups that followed the same evolutionary pattern. (Figure 5). Due to the implementation of different algorithms, OrthoMCL and GeneSeqToFamily split protein families into different sizes (L. Li, Stoeckert, and Roos 2003; Thanki et al. 2018). OrthoMCL usually subdivides a family from GeneSeqToFamily into smaller orthogroups. By using both methods we are able to identify losses of a single protein within large complex families as well as losses of all members of a large protein family. We combined all outputs by identifying orthologs to the Arabidopsis proteome either directly from the results or by reciprocal blastp between rice and Arabidopsis. Four genes were identified by both pipelines including *EDS1* and *PAD4,* an RNL from the *ADR1* clade and a *REGULATOR OF CHROMOSOME CONDENSATION 1-LIKE (RCC1-like)*. The former three are known to be involved in plant defence, whilst *RCC1-like* has not previously been implicated in defence responses. We further focused on the 44 genes identified by either OrthoMCL or GeneSeqToFamily methods (Figure 5, Supplemental Table 8, 9). We designated genes absent only among species without EDS1 as *AngioSperm Typically-Retained, EDS1-Lost (ASTREL)* and hypothesised some may also play a role alongside EDS1/PAD4 and RNLs in immunity.

To further narrow down our list of ASTREL candidates to those with a potential function in the immune pathway, we looked for presence or absence of the ASTREL genes in the asparagus genome. Asparagus is also missing EDS1, PAD4, RNLs and NDR1 and therefore provides an additional filter. This resulted in a list of 5 orthogroups which contained 16 Arabidopsis genes, 12 of which belonged to EDS1, PAD4 or RNL protein families and 4 genes which have not previously been implicated in effector triggered immunity (Table 1, Supplemental Table 11).

**Table 1:**
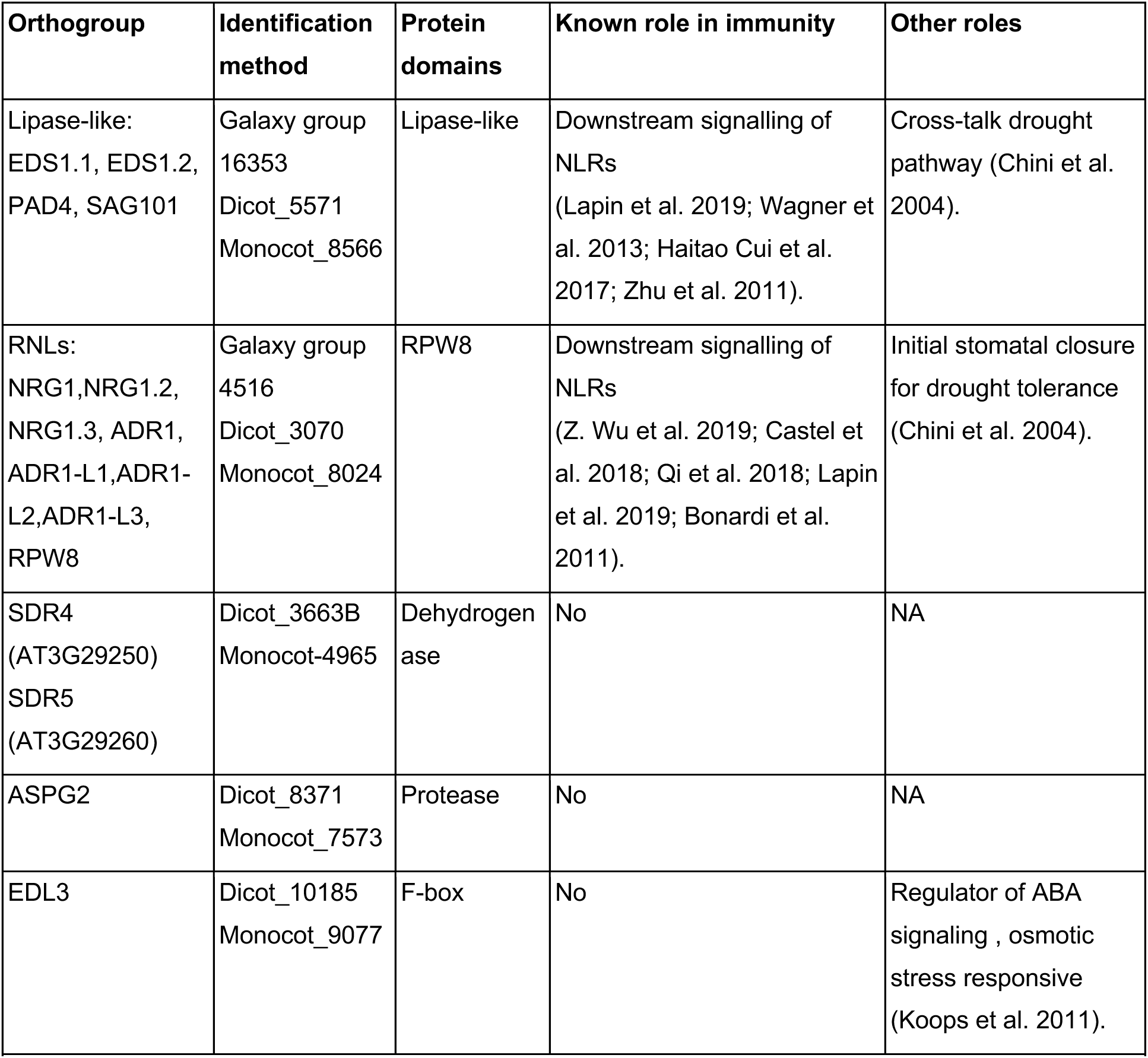
ASTREL high confidence orthogroups absent in all 5 species (duckweed, eelgrass, humped bladderwort, corkscrew plant and asparagus) which are missing EDS1.

### Arabidopsis and rice homologs of ASTREL genes are differentially regulated upon drought response and disease resistance

To understand whether ASTREL genes are differentially expressed during biotic stress or involved in any other specific plant response pathways, we looked for co-expression patterns. We analysed orthologs in Arabidopsis and rice using Arabidopsis mRNAseq and Arabidopsis Affymetrix ATH1 array perturbation experiments (53, 489) and rice RNAseq and microarray datasets (25, resp. 142) (Supplemental Figure 8). Utilising this dataset allowed us to identify two distinct stresses, biotrophic pathogen and ABA/drought stress, under which ASTREL genes are differentially expressed.

We hypothesised that due to the role of EDS1 and RNLs in effector triggered immunity the ASTREL genes may be differentially expressed upon pathogen infection. We first analysed the whole 44 ASTREL genes in full (Supplemental Data 3, Supplemental Figure 8), and then focused on the stringent set of ASTREL genes that were additionally absent in the genome of Asparagus (Supplemental Figure 9 Supplemental Data 1,2,4). As expected, in *Arabidopsis,* EDS1, PAD4 and RNLs were typically upregulated to a much greater extent by biotrophic pathogen infection than necrotrophic pathogen infection (Figure 6). Due to the known role of EDS1 and the RNL NRG1 downstream of TNLs, we investigated if ASTREL gene expression was regulated by nicotinamide, a recently described enzymatic product of TNLs (Wan et al. 2019; Horsefield et al. 2019). The gene expression changes resulting from addition of nicotinamide clustered with those of biotrophic pathogen treatment: PAD4 and all RNLs except ADR1-L3 and NRG1.3 (p-value <0.05, Fold change > 1.5) were found to be upregulated in at least one time point. A subset of the ASTREL genes, made up of EDS1 and SDR4, show opposing expression patterns upon addition of biotrophic pathogens and nicotinamide in Arabidopsis.

**Fig 6.**
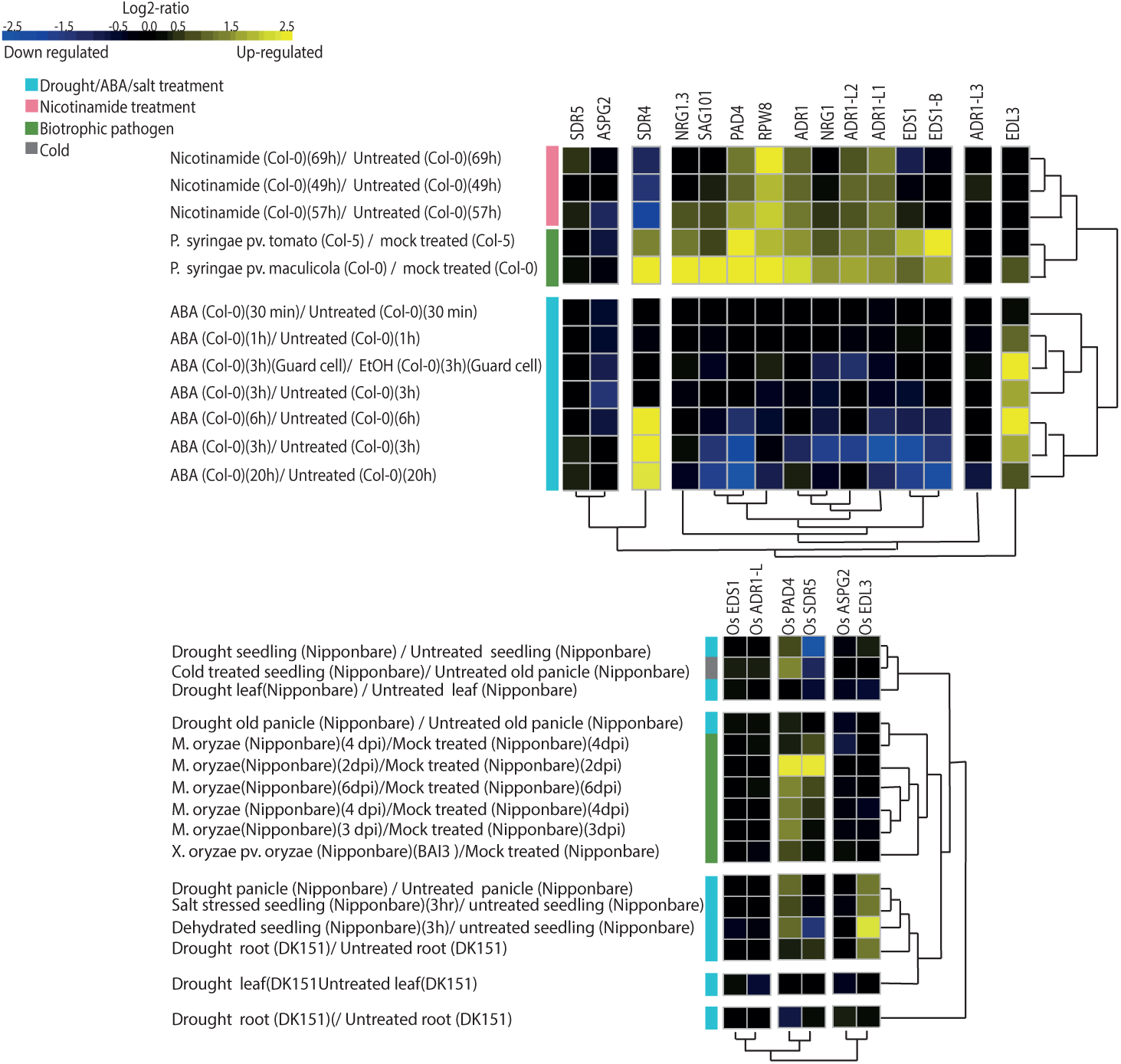
Differential gene expression analysis of arabidopsis and rice high confidence *ASTREL* genes upon biotic and abiotic stress. (A) Pearson hierarchical clustering of differential gene expression of ASTREL genes from *A. thaliana* upon pathogen, ABA and nicotinamide treatments. (B) Pearson hierarchical clustering of differential gene expression of ASTREL genes from rice upon pathogen, drought, cold and salt treatment.

To investigate if any other perturbations regulated ASTREL gene expression, we used hierarchical clustering to identify treatments that caused similar widespread changes in ASTREL gene expression (Supplemental Figure 9 Supplemental data 4). We noticed that drought and ABA treatment also resulted in the differential expression of ASTREL genes in Arabidopsis. Pearson clustering identified three clades of differential expressed ASTREL genes upon drought and ABA response: down-regulated (EDS1, PAD4 and RNLs), upregulated (SDR4, and EDL3) and no effect on expression (SDR5, ASPG-2 and ADR1-L3). Therefore, in addition to pathogen stress we identified ABA/drought to be a regulator of ASTREL gene expression.

To investigate whether the effects of these stresses were specific to Arabidopsis, we mined available gene expression data for ASTREL orthologs in rice. Upon Pearson clustering we observed the same pattern of separation of gene expression response between pathogens and drought (Figure 6B, Supplemental Figure 10, 11). However, the differential expression of genes upon pathogen treatment grouped into different clades, the OsSDR and OsPAD4, were highly upregulated upon *Magnaporthe oryzae* treatment with less drastic expression changes upon drought. We also noted a similar differential expression pattern in rice of OsSDR and OsPAD4 when subjected to either cold stress or drought. Conversely, the OsEDL3.1 clade was consistently upregulated during drought stress. In contrast to Arabidopsis OsEDS1 and OsADR1-L were not upregulated upon biotrophic pathogen treatment of rice. Altogether, the expression data supports the role of ASTREL genes in biotic interactions and provides a link between the EDS1/PAD4 pathway and ABA response.

## Discussion

In this study, we investigated the convergent loss of NLR genes across multiple plant lineages, and linked this phenomenon to gene loss in the ETI immune signalling pathway. Using comparative genomics, we identified a set of genes that were convergently lost among lineages together with the immune signalling genes EDS1/PAD4. The identified genes were implicated in defence and drought response pathways by differential co-expression analyses.

It has been previously shown that there is high variability in the number of NLRs between even closely related species due to lineage specific expansions and contractions (Sarris et al. 2016; Kroj et al. 2016; Van de Weyer et al. 2019; Gao et al. 2018). Phylogenetic analysis showed that most of them retained representatives of major NLR classes including members of RNL and CNL clades, and TNLs among dicots. We identified five species, duckweed, eelgrass, asparagus, corkscrew plant and humped bladderwort, which lack RNL type genes. RNL genes are often subdivided into two main sub-clades: NRG1-like that are required for TNL signalling and are genetically downstream of EDS1, and ADR1-like that are required for the signalling of many CNLs and some TNLs (Castel et al. 2018; Lapin et al. 2019; Z. Wu et al. 2019; Qi et al. 2018). In addition, ADR1-like RNLs also play a role in basal defence, which can limit growth of virulent pathogens (Gantner et al. 2019; Bonardi et al. 2011). Monocots do not have NRG1-like RNLs or TNLs, though they do retain an ADR1 like RNL. We identified that the two dicots, corkscrew plant and humped bladderwort, also lacked TNLs. Previous work has established that absence of TNLs, in both monocots and a few dicot lineages, coincides with the absence of the RNL *NRG1 (Lapin et al. 2019; J. Kim et al. 2012)* this was also the case for corkscrew plant and humped bladderwort. This suggests that RNLs despite their overall conservation across plant lineages are not required for a minimal plant immune system.

Both TNL receptors and RNL signalling components function together with EDS1/PAD4/SAG101 proteins to induce signalling (Castel et al. 2018; Lapin et al. 2019; Qi et al. 2018; Z. Wu et al. 2019), while some CNLs require the presence of NDR1 (Coppinger et al. 2004; Century, Holub, and Staskawicz 1995; Day, Dahlbeck, and Staskawicz 2006). Previous studies have also shown that despite the absence of TNLs, most monocots retain *EDS1* (Chen et al. 2018) suggesting that it might be required for CNL signalling as well as for basal immunity in monocots. In addition to NLR loss, we were able to identify the convergent loss of the EDS1/PAD4 immune complex in at least five species, which also lacked *SAG101 (Lapin et al. 2019; Wagner et al. 2013; Zhu et al. 2011), NRG1 (Castel et al. 2018; Lapin et al. 2019; Z. Wu et al. 2019; Qi et al. 2018), ADR1 (Bonardi et al. 2011; Z. Wu et al. 2019)* and *NDR1 (Day, Dahlbeck, and Staskawicz 2006; Knepper, Savory, and Day 2011)*. The findings of the recurrent loss of *EDS1/PAD4* and *SAG101* together in duckweed, eelgrass, humped bladderwort, asparagus and corkscrew plant is consistent with genetic, biochemical and structural studies of EDS1/PAD4/SAG101, which show the three proteins function in heterodimeric complexes (Lapin et al. 2019; Wagner et al. 2013).

NDR1 follows a very similar pattern of variation in presence and absence as NRG1 and SAG101. These genes are never found without EDS1 and appear to be lost earlier than PAD4. NDR1 was absent in multiple species despite the retention of CNLs. Interestingly, NDR1 is thought to mediate resistance by controlling fluid loss in the cell (Knepper, Savory, and Day 2011), suggesting that the loss of NDR1 can be linked to traits outside of plant immunity. Both NDR1 and RNL mediated signalling converge in triggering increase in SA and changes in gene expression (Haitao Cui et al. 2017). The SA pathway is conserved in all five species in our study that have lost EDS1, PAD4, SAG101, NDR1 and RNLs. Since four of the species in our analyses have retained and expanded the few remaining CNL clades, it suggests that they are utilizing a yet undiscovered signalling pathway. This is consistent with previous reports of some CNLs, such as RPP13 being independent of all known signalling components (Bittner-Eddy and Beynon 2001). Whether remaining CNLs can induce cell death remains to be tested, although another duckweed *Lemna minor* a close relative of *S. polyrhiza* has been shown to induce reactive oxygen burst and cell death response upon abiotic stress (Wenguo Wang et al. 2016).

The repeated absence of TNLs, SAG101 and NDR1 across many lineages suggests that EDS1/PAD4 and RNLs can function on their own. It has been shown that EDS1/PAD4 and EDS1/SAG101 can function differently in divergent lineages. In Solanaceous species, the EDS1/SAG101 heterodimer is required for NRG1-dependent cell death, although in Arabidopsis it is the EDS1/PAD4 heterodimer that is dominant for cell death (Lapin et al. 2019; Gantner et al. 2019). Based on the functional and evolutionary data (loss of EDS1/PAD4 occurs only after SAG101 and NDR1), we can conclude that EDS1/PAD4 represent a functional unit on their own, and their role can be sub-functionalized in presence of other components. Exactly what constitutes a EDS1/SAG101 pathway appears to be species specific (Lapin et al. 2019; Gantner et al. 2019) and how it is differentiated from EDS1/PAD4 remains unclear. Recently, it has also been shown that PAD4 has domain specific functions in herbivore resistance which are independent of EDS1 (Dongus et al. 2019).

Gaps in the pathways can be filled through comparative phylogenomic analysis. Work on plant symbiosis associations has shown that the presence and absence of a plant’s ability to form symbiosis can be used as a phenotype to correlate with patterns of presence and absence of genes which are part of the molecular pathway of symbiosis formation (Griesmann et al. 2018; Bravo et al. 2016; van Velzen et al. 2018). We took a similar approach testing what other genes have been lost in the species without EDS1, PAD4 and RNLs. We identified orthogroups of genes convergently lost in five monocot and dicot lineages that lost NLRs and immune components discussed above, naming newly identified candidates ASTREL. Our prediction is that some of the ASTREL genes will function together with EDS1/PAD4 in one of their roles that is conserved across monocots and dicots.

Genes which function together often show patterns of conserved co-expression (Hansen et al. 2014). Subsets of ASTREL genes showed similar patterns of expression upon perturbation providing further evidence for functional links. Interestingly, differentially expression of ASTREL genes was only observed in a few stress conditions, including pathogen response, nicotinamide, drought, salinity and ABA response. We found that differential expression of ASTREL genes upon addition of nicotinamide was strikingly similar to response to *P. syringae,* consistent with recent literature implicating nicotinamide in plant immune signalling *(Horsefield et al. 2019; Wan et al. 2019)*. Upon pathogen stimulus there are some *ASTREL* genes which are not differentially expressed, however, we cannot rule out the possibility that those genes may be involved in disease resistance. We have only looked at differential expression in a small subset of the possible combinations of conditions, tissues and pathogens that can result in an immune response. The clear patterns of differential expression offers a starting point to characterize the pathway involving ASTREL genes.

Surprisingly, another perturbation which induced changes in gene expression was drought stress. Expression analysis in Arabidopsis appeared to show this was likely due to drought induced increase in ABA. Previous work has demonstrated antagonism between ABA and SA hormone response with stress prioritisation seemingly playing out at the level of hormone regulation of gene expression as well as modulating the turnover of NPR1, a positive regulator of SA and negative regulator of JA (Ryals et al. 1997; Cao, Li, and Dong 1998; Shah, Tsui, and Klessig 1997). Consistent with the antagonism data from Arabidopsis (Moeder et al. 2010; Lievens et al. 2017; de Torres Zabala et al. 2009), a cluster of the ASTREL genes containing EDS1 and RNLs are down regulated upon ABA and upregulated upon pathogen stimulus which triggers SA production. Although this pattern holds in dicot species it is not evident in rice. AtEDL3 is an F-box transcription factor which is a positive regulator of an ABA-dependent signalling cascade (Koops et al. 2011). The loss of EDL3 together with immune signalling components suggests that the transcriptional changes might be regulated through theEDS1/PAD4 pathway. While EDS1 and PAD4 have been reported to be required for *P. syringae* interference with ABA regulation (T.-H. Kim et al. 2011), the mechanism of this inference is largely unknown. Based on our data, EDL3 might play a key role in EDS1/PAD4 dependent ABA cross-talk.

Based on known interactions, functional data (Bonardi et al. 2011; Castel et al. 2018; Qi et al. 2018; H. Cui et al. 2018; Z. Wu et al. 2019; Horsefield et al. 2019; Wan et al. 2019; Lapin et al. 2019) and co-expression analyses, we assembled a working model of how ASTREL genes might fit in the known plant immunity and drought pathways (Fig. 7, S12, S13 Table). Due to the homology of SDR5 to dehydrogenase reductase enzymes and ASPG2 to aspartic proteases, we hypothesise they would be part of the initial cascade which we know to be initiated by TNLs through their NADase activity (Wan et al. 2019; Horsefield et al. 2019). This is consistent with their unchanged gene expression pattern upon infection, suggesting the proteins are already present at the time of infection. Consistent with the requirement of NRG1 for TNL signalling, we propose that all members of the RNL clade play a role in EDS1 dependent NLR signalling as we find the RNL clade is absent in the independent groups of species missing EDS1. Furthermore, we propose the role of transcription factor ELD3 would be downstream of this signalling unit, regulating the cross-section between SA and ABA pathways (Figure 7).

**Fig 7.**
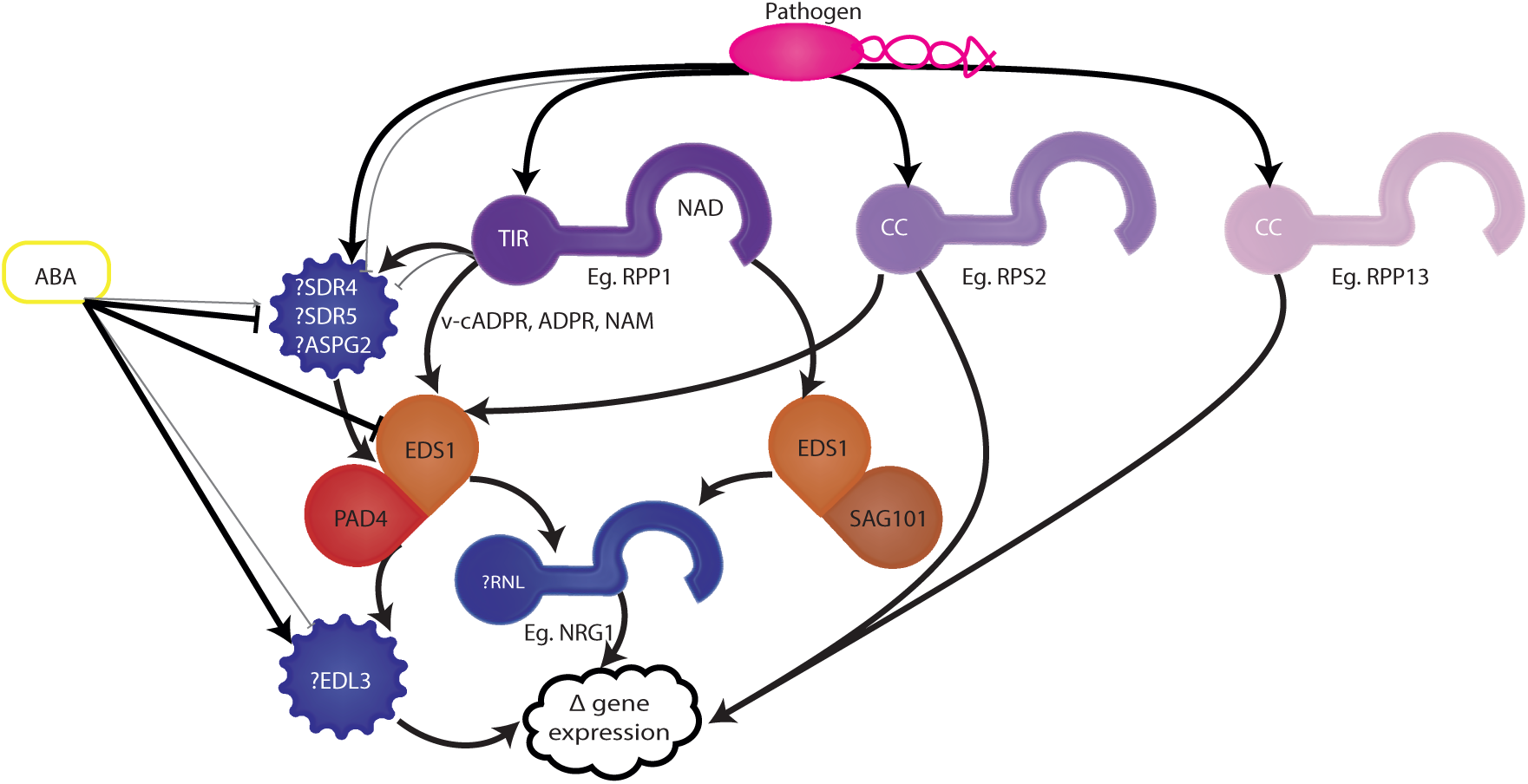
Schematic model of relationship between *ASTREL* genes and known biotic and abiotic stress pathways in *A. thaliana.* Model based on literature review and available gene expression of potential position of ASTREL genes within known *A. thaliana* biotrophic pathogen disease resistance genetic pathway.

Interestingly, four of the five species we identified to be missing ASTREL genes were from water saturated environments ranging from marine to bog habitats. Previous studies have investigated the genome content of one or two aquatic plant species and highlighted gene loss linked to embryogenesis and root development in humped bladderwort (Leushkin et al. 2013), cell wall processes and ABA in duckweed (Michael et al. 2017; W. Wang et al. 2014), and defence response, stomata, terpenoid and hormone pathways in eelgrass (Olsen et al. 2016). This opens up the question of whether the adaptation to a water saturated environment may place selective pressure on signalling pathways including the immune system. There are, however, several species such as lotus and rice which according to our analysis have retained EDS1/PAD4 and other signalling pathway members whilst inhabiting water saturated environments. Furthermore, controversy surrounds the question of whether the most recent common ancestor of monocots could have had an aquatic lifestyle (Du, Wang, and Consortium 2016; Givnish et al. 2018). It is nevertheless tempting to speculate that loss of NLR and EDS1 pathway can be a byproduct of the physiological changes and differing environmental selection pressures associated with derived lifestyles. The transcriptome data suggests that loss of EDS1/PAD4 might be more frequent in monocots than dicots and availability of more genomes could help to determine which environmental and/or physiological traits that influence loss of this pathway.

Until now, discoveries of crucial components of the plant immune system have relied on mutant screens, proteomic analyses and differential expression analysis. Here we have shown a complementary approach to identify potential actors in the plant immune system, which can circumvent issues of genetic redundancy by harnessing conservation and independent transitions in distantly related plant lineages. In breeding for durable disease resistance, NLRs are often stacked to slow the rate of breakdown of resistance by fast evolving pathogens. The downstream signalling components required for NLRs within a stack are rarely considered but if they converge on a single helper or signaller, this creates a strong selection for effectors that would compromise the downstream component and subsequently break the defence conferred by several NLRs at once. In addition, it is crucial to understand the conservation of downstream signalling components to facilitate the successful interspecies transfer of *NLR*s. Our approach has highlighted the conservation of EDS1/PAD4 as a functional unit and identified candidate genes involved in EDS1-mediated immunity.

Our analysis also provides fundamental understanding towards a minimum plant immune system that does not depend on EDS1/PAD4 signalling. We propose duckweed as a new model system, a rapid growing small plant whose reduced ETI immune system could provide a reduced-complexity background for investigating plant immunity. Unexpectedly, the *ASTREL* genes have begun to further elucidate the complex cross-talk between the plant immune system and drought tolerance and we provide evidence for specific transcription factors that can mediate this crosstalk through EDS1/PAD4. Future studies could use the ASTREL genes to test for their functional roles in immune signalling and to further query the interconnection between abiotic and biotic stress pathways.

## Materials and methods

### Genomic datasets used in this study

Genomic assemblies and annotations were obtained from: Phytozome V12 (https://phytozome.jgi.doe.gov/pz/portal.html) for *A. coerulea*, *A. comosus* (v3), *A. hypochondriacus* (325_v1.0), *A. thaliana* (167_TAIR9), *A. trichopoda* (291_v1.0), E. guttata (256_v2.0), *O. sativa* (v7), *O. thomaeum*, *S. lycopersicum* (390_v2.5), *S. moellendorffii* (91_v1), *S. polyrhiza* (v2) and *Z.marina* (v2.2); from COGE (https://genomevolution.org/coge/) for *U. gibba (29027)*; from KEGG for *N. nucifera* (4432), *E. guineesis* (TO3921); from NCBI *P. equestrius* (PRJNA382149), from Ash Tree Genomes (http://www.ashgenome.org/data) for *F. excelsior* (BATG-0.5), from Ensembl for *Dioscorea rotundata* (TDr96_F1_Pseudo_Chromosome_v1.0) and Maize genome database (https://www.maizegdb.org/) for *Z. mays* (AGPv4). The BUSCO scores were calculated using BUSCO (v1.22) to compare proteomes to the embryophyta_odb9 BUSCO lineage (S12, S13 Table) (Simão et al. 2015).

### Annotation, alignment and phylogenetic analysis of NLRs

To annotate NLRs in plant proteomes, the MEME suite (T. L. Bailey et al. 2006) based tool NLR-parser (Steuernagel et al. 2015) was used in addition to the updated version of the NLR-ID pipeline (Sarris et al. 2016) available at https://github.com/krasileva-group/plant_rgenes/releases/tag/ASTREL_v1_2019_10.1101.572560. Annotations were combined into a non-redundant list of putative NLRs. Where multiple transcripts were present, these were filtered to retain the longest transcript. In addition a series of characterised NLRs were added. These are available at https://itol.embl.de/sharedProject.cgi user ID: erin_baggs. The HMMALIGN programme from the HMMER3.0 suite (Wheeler and Eddy 2013) was used to align proteins to the NB-ARC1_ARC2_prank_aln_domain_ONLY HMM of the NB-ARC domain (P. C. Bailey et al. 2018). The alignment was trimmed to the NB-ARC domain region using Belvu (Barson and Griffiths 2016) and columns and sequences with over 80% gaps where removed. The NB-ARC domain of the remaining NLRs was then manually curated in Jalview (Waterhouse et al. 2009) allowing no more than 2 consecutive characteristic NB-ARC domain motifs (Walker A, RNBS-A, WALKER-B, RNBS-C, GLPL, RNBS-D) to be absent. A maximum likelihood phylogenetic tree was constructed using RAXML-MPI (v.8.2.9) (Stamatakis 2014) with parameters set as: -f a -x 1123 -p 2341 -# 100 -m PROTCATJTT. Tree visualisation and annotation using iTOL and are available at: http://itol.embl.de/shared/erin_baggs.

NLRs gains and losses across the phylogeny, were computed using tree reconciliation (Notung 2.9) (Stolzer et al. 2012) based on a maximum likelihood species tree built on the concatenated amino acid sequences of 1:1 orthologs across all 18 species identified by GeneSeqToFamily and built using RAXML (8.2.11, JTT model) and a gene tree comprising 2237 NLRs.

### Ortholog identification

To identify orthologs of specific genes of interest, we used reciprocal BLAST searches using blastp with parameters -max_target_seqs 1 -evalue 1e-10 (BLAST+ 2.2.28.mt) (Camacho et al. 2009). If upon reciprocal BLAST, a homologous gene was not identified, we used ensemble gene trees to check for recent duplication events in the Arabidopsis lineage and to confirm our results. Results were manually inspected and filtered using a script available at project github. The presence of *EDS1* in *O. thomaeum* was validated using RNAseq data (BioProject SRS957807) mapped onto the Oropetium V1 Bio_nano genome assembly (http://www.oropetium.org/resources) (VanBuren et al. 2018, 2015) using HISAT2 (D. Kim, Langmead, and Salzberg 2015). Bam files were processed using SAMtools-1.7 (H. Li et al. 2009) and results visualised using IGB (Nicol et al. 2009). The absence of EDS1 was validated by running HMMSEARCH was run on the proteomes of *A. thaliana*, *S. polyrhiza* and *Z. marina* using the Lipase 3 domain which is characteristic of EDS1. Proteins containing the domain were then aligned against the domain using HMMALIGN and the Pfam Lipase 3 HMM. The alignment was manually curated in Belvu before submission to RAXML as above.

To identify orthologous gene groups, OrthoMCL (v2.0.9) (L. Li, Stoeckert, and Roos 2003) was used as described previously (Johnson et al., 2018). Due to large computational requirements, we ran OrthoMCL separately for monocots (*Z. marina, S. polyrhiza, P. equestris, Dioscorea rotundata*, *Eleais guineesis, O. thomaeum, Oryza sativa* and *Zea mays*) and dicots (*Aquilegia coerulea, Nelumbo nucifera, Arabidopsis thaliana, Amaranthus hypochondriachus, Solanum lycopersicum, Fraxinus excelsior, E. guttata, G. aurea* and *U. gibba*) both with the outgroups *Selaginella moellendorffii* and *Amborella trichopoda.* We overlaid monocot analyses with dicot orthogroups by mapping the monocot gene IDs to the Arabidopsis homologs using reciprocal blastp. Additionally, we applied the GeneSeqToFamily pipeline (Thanki et al. 2018) on all 19 genomes. We then identified orthogroups lost in aquatic species (*S. polyrhiza*, *Z. marina, U. gibba*, *G. aurea)* but retained in all terrestrial species of the same monocot/dicot clade. We cannot preclude the possibility that some gene families that have been convergently lost in aquatic species have not also been lost independently in some of the terrestrial lineages. After grouping and manual curation (see Material and Methods), gene families for which pan-species evolutionary history had been previously established were compared to gene families in our orthogroups. This curation led to the decision to mask *S. moellendorffii*, *A. trichopoda*, *A. coerulea* and *D. rotundata* from later analysis with the former two species rarely having homologs due to large phylogenetic distance and the latter two species having many erroneous gene fusion annotations. The analyses were filtered using scripts available at https://github.com/krasileva-group/Aquatic_NLR. For comparison of OrthoMCL gene families retained between monocots and dicots, we used Arabidopsis and rice proteins as representative genome members and cross-referenced them using blastp reciprocal search (e-value cutoff 1e-10). The validity of this approach was checked on random protein families using Plant Ensembl trees (http://plants.ensembl.org/index.html). The rice gene ids were converted between Phytozome and Plant Ensembl using Plant Ensembl conversion tool (http://rapdb.dna.affrc.go.jp/tools/converter/run).

### Pairwise synteny analysis for EDS1, PAD4, ADR1, between *A. comosus* and *S. polyrhiza* as performed using the output of GeneSeqToFamily and from SynMap2

To identify the syntenic region in *S. polyrhiza* that corresponds to the region where EDS1, PAD4 and ADR1 are present in *A. comosus*, GeneSeqToFamily was used to identify orthologs between *A. comosus* and *S. polyrhiza (Thanki et al. 2018)*. The location of the orthologous genes immediately upstream and downstream of the ASTREL genes in *A. comosus* was identified in *S. polyrhiza* and its genomic location recorded. Synteny between genes in the neighbourhood of EDS1, PAD4 and ADR1 was further confirmed by submission of genomic sequence upstream and downstream of ASTREL genes of interest in *A. comosus* to SynMap2 (Haug-Baltzell et al. 2017), which confirmed the presence of microsynteny at the DNA level on scaffolds of *S. polyrhiza* oxford v3 genome equivalent to those identified in chromosomal assembly of *S. polyrhiza*.

### Expression profiling

The expression analysis for this study was performed and visualised using the 706 rice mRNA samples and 2,836 rice Affymetrix rice genome array samples available on Genevestigator v7.0.3 (https://genevestigator.com) (Hruz et al. 2008) a list of NCBI GEO dataset identifiers are available(https://github.com/krasileva-group/Aquatic_NLR/tree/master/Supplemental-Data/Supplement_Differential_Expression_All). Arabidopsis RNAseq and Affymetrix ATH1 array (53, 489) and rice RNAseq and microarray datasets (25, resp. 142) available on Genevestigator v7.0.3 (https://genevestigator.com) were used to visualise conditions resulting in differential patterns of gene expression. A list of NCBI GEO dataset identifiers for the datasets used are available at: (https://github.com/krasileva-group/Aquatic_NLR). The datasets used to investigate further drought and pathogen infection induced changes in gene expression across the 52 *A. thaliana ASTREL*, monocot and dicot overlapping genes were as follows: Microarray - Pathogen - AT-00202 (Craigon et al. 2004), AT-00406 (Craigon et al. 2004), OS-00057 (Yu et al. 2011), OS-00045 (Marcel et al. 2010), OS-00011(Haiyan et al. 2012), microarray - nicotinamide AT-00398 (Dalchau et al. 2010), microarray - drought - AT-00110 (Kilian et al. 2007), AT-00433 (Pandey et al. 2010), AT-00468 (Böhmer and Schroeder 2011), AT-00541 (T.-H. Kim et al. 2011), OS-00008 (Jain et al. 2007), OS-00041 (D. Wang et al. 2011), OS-00224 (Krishnan et al. 2010) microarray - salt - OS-00008 (Jain et al. 2007). Hierarchical clustering was generated considering both genes and conditions with parameters of Pearson’s correlation coefficient and optimal leaf-ordering.

## Acknowledgements

The authors are grateful to all members of the Krasileva group and their many colleagues, for thoughtful discussion on the presented material. We thank Dr. Daniil Prigozhin and Dr. Janina Tamborski for suggestions on the manuscript and Dr. Burkhard Steuernagel for his help with NLRannotator software. The high-performance computing resources and services used in this work were supported by the EI Scientific Computing group alongside the NBIP Computing infrastructure for Science (CiS) group.

## Data Availability Statement

All genomic and gene expression datasets used for this study were already available from public repositories (see Materials and Methods). All scripts used to analyze data are available from https://github.com/krasileva-group/Aquatic_NLR. High resolution phylogenetic tree of NLRs across 18 species in study is available on iToL: http://itol.embl.de/shared/erin_baggs.

## Author contributions

ELB, WH and KVK designed the study. ELB performed NLR, phylogenetic, analysis. AST ran analysis with the GeneSeqToFamily pipeline. ELB assisted by CS analysed orthogroups from OrthoMCL and GeneSeqToFamily to identify ASTRAL candidates. RNAseq analysis of ASTRAL genes performed by ELB and statistical tests by ELB and KVK. Model of gene interaction by ELB. Figure preparation by RO and ELB. All authors contributed to final manuscript. All authors read and approved final manuscript.

## Supplementary material

Supplementary table 1 - Table of the counts of Pfam NB-ARC domain for 95 species with available genomes.

Supplementary table 2 - Table of genome assembly size, NLRannotator NLR number prediction and number of encoded high and low confidence NLR genes defined through alignment.

Supplementary table 3 - a) Table of number of BUSCOs identified. b) Table of actin and kinase HMM domain encoding genes present in the genomes of species analysed with OrthoMCL and GeneSeqToFamily.

Supplementary table 4 - Table of genes and gene identifiers of characterised immune genes.

Supplementary table 5 - Table of NLR number and presence absence of ETI components across 95 plant species genomes.

Supplementary table 6 - Plot of presence and absence of 7 characterised ETI components across transcriptomes of dicots and magnoliids.

Supplementary table 7 - Plot of presence and absence of 7 characterised ETI components across transcriptomes of monocots.

Supplementary table 8 - Table of *Arabidopsis ASTREL* gene identifiers.

Supplementary table 9 - Table of ASTREL gene orthogroups and representative rice genes.

Supplementary table 10 - Plot of reciprocal BLAST of ASTREL genes to the asparagus genome.

Supplementary table 11 - List of high confidence ASTREL gene Identifiers.

Supplementary table 12 - Literature linking ASTREL genes to abscisic acid.

Supplementary table 13 - Literature linking ASTREL genes to effector triggered immunity.

Supplementary fig 1 - Histogram of the number of genes with NB-ARC domains for 109 available plant genomes.

Supplementary fig 2 - Maximum likelihood phylogeny of the NB-ARC domain of RNL genes across the 18 representative species.

Supplementary fig 3 -Extended species plot of presence and absence of known immunity genes from reciprocal BLAST searches.

Supplementary fig 4 - Maximum-likelihood gene tree from proteins containing lipase-3 HMM domain.

Supplementary fig 5 - Plot of presence and absence of 7 characterised ETI components across transcriptomes of dicots and magnoliids.

Supplementary fig 6 - Plot of presence and absence of 7 characterised ETI components across transcriptomes of monocots.

Supplementary fig 7 - Gene tree for EDS1, PAD4 and SAG101 gene family from orthologs identified from GeneSeqToFamily.

Supplementary fig 8 - Heatmap of differential expression of ASTREL genes across publicly available RNAseq experiments using Geneinvestigator.

Supplementary fig 9 - Heatmap of differential expression of high confidence ASTREL genes across publicly available RNAseq experiments using Geneinvestigator.

Supplementary fig 10 - Heatmap of differential expression of high confidence ASTREL genes across publicly available RNAseq experiments using Geneinvestigator.

Supplementary fig 11 - Heatmap of differential expression of high confidence ASTREL genes across publicly available microarray experiments using Geneinvestigator.

Supplementary data 1 - Experiment identifiers and differential expression values for all the mRNAseq datasets queried to identify patterns of differential expression among ASTREL high confidence genes.

Supplementary data 2 - Experiment identifiers and differential expression values for all the mycroarray datasets queried to identify patterns of differential expression among ASTREL high confidence genes.

Supplementary data 3 - Heatmap of differential expression of ASTREL genes across publicly available microarray experiments using Geneinvestigator.

Supplementary dataset 4 - Heatmap of differential expression of high confidence ASTREL genes across publicly available microarray experiments using Geneinvestigator.

## Supplemental figure legends

Sfig 1 - **Graph of the number of genes with an NB-ARC domain across 95 plant species genomes.** On the Y-axis is the name of the genome variant used in the analysis consistent with Phytozome and Ensembl Plant databases. The colour of the bars distinguishes species based on taxonomy. Numbers at the end of bars are the number of NB-ARC domains identified in that species.

Sfig 2 - **Maximum likelihood phylogeny based on the NB-ARC domain of RNL clade.** Bootstrap values are given on branches with leaf labels as the genus then species followed by the gene identifier. The red bar indicates RNLs and the orange and blue bars distinguish the NRG1 from the ADR1 clade.

Sfig 3 - **Table of presence and absence of immunity associated components when queried by reciprocal BLASTp.** Red indicates the absence of an ortholog by reciprocal protein BLAST whilst green indicates and identified ortholog. The first row gives gene name followed by gene name followed by gene identifier from *A. thaliana*.

Sfig 4 - **Maximum likelihood phylogeny of proteins with a lipase 3 HMM Motif.** Tree was constructed using the Amborella, rice, arabidopsis, duckweed and eelgrass proteomes. Blue branches indicate the EDS1/PAD4 clade. Bootstrap values >80 are indicated with a circle. Leafs are the gene identifiers for species proteins.

Sfig 5 - **Identification of immune gene orthologs in Lamiales species with available transcriptomes.** Phylogeny from NCBI taxonomy. Gene names of the arabidopsis query proteins are given along the top of the plot. Black dots represent identified ortholog within transcriptome. Grey circles indicate no orthologous transcript identified in transcriptome.

Sfig 6 - **Identification of immune gene orthologs in magnoliids and early diverging monocot species with available transcriptomes.** Phylogeny from NCBI taxonomy. Gene names of the arabidopsis and rice query proteins are given along the top of the plot. Black dots represent identified ortholog within transcriptome. Grey circles indicate no orthologous transcript identified in transcriptome.

Sfig 7 - **Phylogenetic tree of EDS1 proteins identified by GeneSeqToFamily.** Leaves give gene identifier followed00 by the first letter of genus and then 3 letters of species name.

Sfig 8 - **Heatmap of changes in gene expression of arabidopsis ASTREL genes upon available RNAseq conditions.** Blue indicates downregulation and yellow indicates upregulation.

Sfig 9 - **Heatmap of changes in gene expression of arabidopsis high confidence ASTREL genes upon available RNAseq conditions.** Blue indicates downregulation and yellow indicates upregulation. Pearson correlation was used to group conditions and genes, with optimal leaf ordering.

Sfig 10 - **Heatmap of changes in gene expression of rice high confidence ASTREL genes upon available RNAseq conditions.** Blue indicates downregulation and yellow indicates upregulation. Pearson correlation was used to group conditions and genes, with optimal leaf ordering.

Sfig 11 - **Heatmap of changes in gene expression of rice high confidence ASTREL genes upon available microarray conditions.** Blue indicates downregulation and yellow indicates upregulation. Pearson correlation was used to group conditions and genes, with optimal leaf ordering.

Sdata 1 - **Table of ASTREL gene expression RNAseq perturbations.** Experiment identifiers and differential expression values for all the mRNAseq datasets queried to identify patterns of differential expression among ASTREL high confidence genes.

Sdata 2 - **Table of ASTREL gene expression microarrray perturbations.** Experiment identifiers and differential expression values for all the microarray datasets queried to identify patterns of differential expression among ASTREL high confidence genes.

Sdataset 3 - **Heatmap of changes in gene expression of arabidopsis ASTREL genes upon available microarray conditions.** Blue indicates downregulation and yellow indicates upregulation.

Sdataset 4 - **Heatmap of changes in gene expression of arabidopsis high confidence ASTREL genes upon available microarray conditions.** Blue indicates downregulation and yellow indicates upregulation. Pearson correlation was used to group conditions and genes, with optimal leaf ordering.

## References

Baggs, E., G. Dagdas, and K. V. Krasileva. 2017. “NLR Diversity, Helpers and Integrated Domains: Making Sense of the NLR IDentity.” Current Opinion in Plant Biology 38 (August): 59–67.

Bailey, Paul C., Christian Schudoma, William Jackson, Erin Baggs, Gulay Dagdas, Wilfried Haerty, Matthew Moscou, and Ksenia V. Krasileva. 2018. “Dominant Integration Locus Drives Continuous Diversification of Plant Immune Receptors with Exogenous Domain Fusions.” Genome Biology 19 (1): 23.

Bailey, Timothy L., Nadya Williams, Chris Misleh, and Wilfred W. Li. 2006. “MEME: Discovering and Analyzing DNA and Protein Sequence Motifs.” Nucleic Acids Research 34 (suppl_2): W36–373.

Barson, Gemma, and Ed Griffiths. 2016. “SeqTools: Visual Tools for Manual Analysis of Sequence Alignments.” BMC Research Notes 9 (1): 39.

Bernoux, Maud, Thomas Ve, Simon Williams, Christopher Warren, Danny Hatters, Eugene Valkov, Xiaoxiao Zhang, Jeffrey G. Ellis, Bostjan Kobe, and Peter N. Dodds. 2011. “Structural and Functional Analysis of a Plant Resistance Protein TIR Domain Reveals Interfaces for Self-Association, Signaling, and Autoregulation.” Cell Host & Microbe 9 (3): 200–211.

Bhandari, Deepak D., Dmitry Lapin, Barbara Kracher, Patrick von Born, Jaqueline Bautor, Karsten Niefind, and Jane E. Parker. 2019. “An EDS1 Heterodimer Signalling Surface Enforces Timely Reprogramming of Immunity Genes in Arabidopsis.” Nature Communications 10 (1): 772.

Bittner-Eddy, P. D., and J. L. Beynon. 2001. “The Arabidopsis Downy Mildew Resistance Gene, RPP13-Nd, Functions Independently of NDR1 and EDS1 and Does Not Require the Accumulation of Salicylic Acid.” Molecular Plant-Microbe Interactions: MPMI 14 (3): 416–21.

Böhmer, Maik, and Julian I. Schroeder. 2011. “Quantitative Transcriptomic Analysis of Abscisic Acid-Induced and Reactive Oxygen Species-Dependent Expression Changes and Proteomic Profiling in Arabidopsis Suspension Cells.” The Plant Journal. https://doi.org/10.1111/j.1365-313x.2011.04579.x.

Bonardi, Vera, Saijun Tang, Anna Stallmann, Melinda Roberts, Karen Cherkis, and Jeffery L. Dangl. 2011. “Expanded Functions for a Family of Plant Intracellular Immune Receptors beyond Specific Recognition of Pathogen Effectors.” Proceedings of the National Academy of Sciences of the United States of America 108 (39): 16463–68.

Bravo, Armando, Thomas York, Nathan Pumplin, Lukas A. Mueller, and Maria J. Harrison. 2016. “Genes Conserved for Arbuscular Mycorrhizal Symbiosis Identified through Phylogenomics.” Nature Plants 2: 15208.

Camacho, C., G. Coulouris, V. Avagyan, N. Ma, J. Papadopoulos, K. Bealer, and T. L. Madden. 2009. “BLAST+: Architecture and Applications.” BMC Bioinformatics 10: 42–421.

Cao, H., X. Li, and X. Dong. 1998. “Generation of Broad-Spectrum Disease Resistance by Overexpression of an Essential Regulatory Gene in Systemic Acquired Resistance.” Proceedings of the National Academy of Sciences of the United States of America 95 (11): 6531–36.

Castel, Baptiste, Pok-Man Ngou, Volkan Cevik, Amey Redkar, Dae-Sung Kim, Ying Yang, Pingtao Ding, and Jonathan D. G. Jones. 2018. “Diverse NLR Immune Receptors Activate Defence via the RPW8-NLR NRG1.” The New Phytologist 0 (0). https://doi.org/10.1111/nph.15659.

Catanzariti, Ann-Maree, Peter N. Dodds, Thomas Ve, Bostjan Kobe, Jeffrey G. Ellis, and Brian J. Staskawicz. 2010. “The AvrM Effector from Flax Rust Has a Structured C-Terminal Domain and Interacts Directly with the M Resistance Protein.” Molecular Plant-Microbe Interactions: MPMI 23 (1): 49–57.

Century, K. S., E. B. Holub, and B. J. Staskawicz. 1995. “NDR1, a Locus of Arabidopsis Thaliana That Is Required for Disease Resistance to Both a Bacterial and a Fungal Pathogen.” Proceedings of the National Academy of Sciences of the United States of America 92 (14): 6597–6601.

Chen, Guiping, Bo Wei, Guoliang Li, Caiyan Gong, Renchun Fan, and Xiangqi Zhang. 2018. “TaEDS1 Genes Positively Regulate Resistance to Powdery Mildew in Wheat.” Plant Molecular Biology 96 (6): 607–25.

Chini, A., J. J. Grant, M. Seki, K. Shinozaki, and G. J. Loake. 2004. “Drought Tolerance Established by Enhanced Expression of the CC-NBS-LRR Gene, ADR1, Requires Salicylic Acid, EDS1 and ABI1.” The Plant Journal: For Cell and Molecular Biology 38 (5): 810–22.

Coppinger, Peter, Peter P. Repetti, Brad Day, Douglas Dahlbeck, Angela Mehlert, and Brian J. Staskawicz. 2004. “Overexpression of the Plasma Membrane-Localized NDR1 Protein Results in Enhanced Bacterial Disease Resistance in Arabidopsis Thaliana.” The Plant Journal: For Cell and Molecular Biology 40 (2): 225–37.

Craigon, David J., Nick James, John Okyere, Janet Higgins, Joan Jotham, and Sean May. 2004. “NASCArrays: A Repository for Microarray Data Generated by NASC’s Transcriptomics Service.” Nucleic Acids Research 32 (Database issue): D575–77.

Cui, Haitao, Enrico Gobbato, Barbara Kracher, Jingde Qiu, Jaqueline Bautor, and Jane E. Parker. 2017. “A Core Function of EDS1 with PAD4 Is to Protect the Salicylic Acid Defense Sector in Arabidopsis Immunity.” The New Phytologist 213 (4): 1802–17.

Cui, H., J. Qiu, Y. Zhou, D. D. Bhandari, C. Zhao, J. Bautor, and J. E. Parker. 2018. “Antagonism of Transcription Factor MYC2 by EDS1/PAD4 Complexes Bolsters Salicylic Acid Defense in Arabidopsis Effector-Triggered Immunity.” Molecular Plant 11 (8): 1053–66.

Dalchau, Neil, Katharine E. Hubbard, Fiona C. Robertson, Carlos T. Hotta, Helen M. Briggs, Guy-Bart Stan, Jorge M. Gonçalves, and Alex A. R. Webb. 2010. “Correct Biological Timing in Arabidopsis Requires Multiple Light-Signaling Pathways.” Proceedings of the National Academy of Sciences of the United States of America 107 (29): 13171–76.

Day, Brad, Douglas Dahlbeck, and Brian J. Staskawicz. 2006. “NDR1 Interaction with RIN4 Mediates the Differential Activation of Multiple Disease Resistance Pathways in Arabidopsis.” The Plant Cell 18 (10): 2782–91.

Deutekom, Eva S., Julian Vosseberg, Teunis J. P. van Dam, and Berend Snel. 2019. “Measuring the Impact of Gene Prediction on Gene Loss Estimates in Eukaryotes by Quantifying Falsely Inferred Absences.” PLoS Computational Biology 15 (8): e1007301.

Dodds, P. N., G. J. Lawrence, and J. G. Ellis. 2001. “Six Amino Acid Changes Confined to the Leucine-Rich Repeat Beta-Strand/beta-Turn Motif Determine the Difference between the P and P2 Rust Resistance Specificities in Flax.” The Plant Cell 13 (1): 163–78.

Dongus, Joram A., Deepak D. Bhandari, Monika Patel, Lani Archer, Lucas Dijkgraaf, Laurent Deslandes, Jyoti Shah, and Jane E. Parker. 2019. “Arabidopsis PAD4 Lipase-like Domain Is a Minimal Functional Unit in Resistance to Green Peach Aphid.” bioRxiv. https://doi.org/10.1101/769125.

Du, Zhi-Yuan, Qing-Feng Wang, and China Phylogeny Consortium. 2016. “Phylogenetic Tree of Vascular Plants Reveals the Origins of Aquatic Angiosperms.” Journal of Systematics and Evolution 54 (4): 342–48.

Gantner, Johannes, Jana Ordon, Carola Kretschmer, Raphael Guerois, and Johannes Stuttmann. 2019. “An EDS1-SAG101 Complex Is Essential for TNL-Mediated Immunity in Nicotiana Benthamiana.” The Plant Cell, tpc.00099.2019.

Gao, Y., W. Wang, T. Zhang, Z. Gong, H. Zhao, and G. Z. Han. 2018. “Out of Water: The Origin and Early Diversification of Plant R-Genes.” Plant Physiology 177 (1): 82–89.

Givnish, Thomas J., Alejandro Zuluaga, Daniel Spalink, Marybel Soto Gomez, Vivienne K. Y. Lam, Jeffrey M. Saarela, Chodon Sass, et al. 2018. “Monocot Plastid Phylogenomics, Timeline, Net Rates of Species Diversification, the Power of Multi-Gene Analyses, and a Functional Model for the Origin of Monocots.” American Journal of Botany. https://doi.org/10.1002/ajb2.1178.

Griesmann, Maximilian, Yue Chang, Xin Liu, Yue Song, Georg Haberer, Matthew B. Crook, Benjamin Billault-Penneteau, et al. 2018. “Phylogenomics Reveals Multiple Losses of Nitrogen-Fixing Root Nodule Symbiosis.” Science 361 (6398): eaat1743.

Haiyan, Hu, Zhuang Jieyun, Zheng Kangle, Liu Mingjiu, and Others. 2012. “Characterization of Differentially Expressed Genes Induced by Virulent and Avirulent Magnaporthe Grisea Strains in Rice.” Plant Omics 5 (6): 542.

Hansen, Bjoern O., Neha Vaid, Magdalena Musialak-Lange, Marcin Janowski, and Marek Mutwil. 2014. “Elucidating Gene Function and Function Evolution through Comparison of Co-Expression Networks of Plants.” Frontiers in Plant Science. https://doi.org/10.3389/fpls.2014.00394.

Haug-Baltzell, A., S. A. Stephens, S. Davey, C. E. Scheidegger, and E. Lyons. 2017. “SynMap2 and SynMap3D: Web-Based Whole-Genome Synteny Browsers.” Bioinformatics 33 (14): 2197–98.

Horsefield, Shane, Hayden Burdett, Xiaoxiao Zhang, Mohammad K. Manik, Yun Shi, Jian Chen, Tiancong Qi, et al. 2019. “NAD+ Cleavage Activity by Animal and Plant TIR Domains in Cell Death Pathways.” Science 365 (6455): 793–99.

Hruz, Tomas, Oliver Laule, Gabor Szabo, Frans Wessendorp, Stefan Bleuler, Lukas Oertle, Peter Widmayer, Wilhelm Gruissem, and Philip Zimmermann. 2008. “Genevestigator v3: A Reference Expression Database for the Meta-Analysis of Transcriptomes.” Advances in Bioinformatics 2008 (July): 420747.

Ibarra-Laclette, Enrique, Eric Lyons, Gustavo Hernández-Guzmán, Claudia Anahí Pérez-Torres, Lorenzo Carretero-Paulet, Tien-Hao Chang, Tianying Lan, et al. 2013. “Architecture and Evolution of a Minute Plant Genome.” Nature 498: 94.

Jain, Mukesh, Aashima Nijhawan, Rita Arora, Pinky Agarwal, Swatismita Ray, Pooja Sharma, Sanjay Kapoor, Akhilesh K. Tyagi, and Jitendra P. Khurana. 2007. “F-Box Proteins in Rice. Genome-Wide Analysis, Classification, Temporal and Spatial Gene Expression during Panicle and Seed Development, and Regulation by Light and Abiotic Stress.” Plant Physiology. https://doi.org/10.1104/pp.106.091900.

Jones, Jonathan D. G., Russell E. Vance, and Jeffery L. Dangl. 2016. “Intracellular Innate Immune Surveillance Devices in Plants and Animals.” Science 354 (6316). http://science.sciencemag.org/content/354/6316/aaf6395.abstract.

Kilian, Joachim, Dion Whitehead, Jakub Horak, Dierk Wanke, Stefan Weinl, Oliver Batistic, Cecilia D’Angelo, Erich Bornberg-Bauer, Jörg Kudla, and Klaus Harter. 2007. “The AtGenExpress Global Stress Expression Data Set: Protocols, Evaluation and Model Data Analysis of UV-B Light, Drought and Cold Stress Responses.” The Plant Journal: For Cell and Molecular Biology 50 (2): 347–63.

Kim, Daehwan, Ben Langmead, and Steven L. Salzberg. 2015. “HISAT: A Fast Spliced Aligner with Low Memory Requirements.” Nature Methods 12 (4): 357–60.

Kim, Jungeun, Chan Ju Lim, Bong-Woo Lee, Jae-Pil Choi, Sang-Keun Oh, Raza Ahmad, Suk-Yoon Kwon, Jisook Ahn, and Cheol-Goo Hur. 2012. “A Genome-Wide Comparison of NB-LRR Type of Resistance Gene Analogs (RGA) in the Plant Kingdom.” Molecules and Cells 33 (4): 385–92.

Kim, Tae-Houn, Felix Hauser, Tracy Ha, Shaowu Xue, Maik Böhmer, Noriyuki Nishimura, Shintaro Munemasa, et al. 2011. “Chemical Genetics Reveals Negative Regulation of Abscisic Acid Signaling by a Plant Immune Response Pathway.” Current Biology: CB 21 (11): 990–97.

Knepper, Caleb, Elizabeth A. Savory, and Brad Day. 2011. “Arabidopsis NDR1 Is an Integrin-Like Protein with a Role in Fluid Loss and Plasma Membrane-Cell Wall Adhesion.” Plant Physiology 156 (1): 286–300.

Koops, P., S. Pelser, M. Ignatz, C. Klose, K. Marrocco-Selden, and T. Kretsch. 2011. “EDL3 Is an F-Box Protein Involved in the Regulation of Abscisic Acid Signalling in Arabidopsis Thaliana.” Journal of Experimental Botany 62 (15): 5547–60.

Krasileva, Ksenia V., Douglas Dahlbeck, and Brian J. Staskawicz. 2010. “Activation of an Arabidopsis Resistance Protein Is Specified by the in Planta Association of Its Leucine-Rich Repeat Domain with the Cognate Oomycete Effector.” The Plant Cell 22 (7): 2444–58.

Krishnan, Arjun, Madana Ambavaram, Utlwang Batlang, and Andy Pereira. 2010. “7. A Resource for Systems Analysis of Transcriptional Modules Involved in Drought Response in Rice.” Systems Analysis of Stress Response in Plants, 138.

Kroj, T., E. Chanclud, C. Michel-Romiti, X. Grand, and J. B. Morel. 2016. “Integration of Decoy Domains Derived from Protein Targets of Pathogen Effectors into Plant Immune Receptors Is Widespread.” The New Phytologist 210 (2): 618–26.

Lapin, Dmitry, Viera Kovacova, Xinhua Sun, Joram A. Dongus, Deepak Bhandari, Patrick von Born, Jaqueline Bautor, et al. 2019. “A Coevolved EDS1-SAG101-NRG1 Module Mediates Cell Death Signaling by TIR-Domain Immune Receptors.” The Plant Cell 31 (10): 2430.

Le Roux, Clémentine, Gaëlle Huet, Alain Jauneau, Laurent Camborde, Dominique Trémousaygue, Alexandra Kraut, Binbin Zhou, et al. 2015. “A Receptor Pair with an Integrated Decoy Converts Pathogen Disabling of Transcription Factors to Immunity.” Cell 161 (5): 1074–88.

Leushkin, Evgeny V., Roman A. Sutormin, Elena R. Nabieva, Aleksey A. Penin, Alexey S. Kondrashov, and Maria D. Logacheva. 2013. “The Miniature Genome of a Carnivorous Plant Genlisea Aurea Contains a Low Number of Genes and Short Non-Coding Sequences.” BMC Genomics 14 (July): 476.

Lievens, Laurens, Jacob Pollier, Alain Goossens, Rudi Beyaert, and Jens Staal. 2017. “Abscisic Acid as Pathogen Effector and Immune Regulator.” Frontiers in Plant Science 8: 587.

Li, Heng, Bob Handsaker, Alec Wysoker, Tim Fennell, Jue Ruan, Nils Homer, Gabor Marth, Goncalo Abecasis, and Richard Durbin. 2009. “The Sequence Alignment/Map Format and SAMtools.” Bioinformatics 25 (16): 2078–79.

Li, Li, Christian J. Stoeckert, and David S. Roos. 2003. “OrthoMCL: Identification of Ortholog Groups for Eukaryotic Genomes.” Genome Research 13 (9): 2178–89.

Maekawa, Takaki, Barbara Kracher, Saskia Vernaldi, Emiel Ver Loren van Themaat, and Paul Schulze-Lefert. 2012. “Conservation of NLR-Triggered Immunity across Plant Lineages.” Proceedings of the National Academy of Sciences 109 (49): 20119–23.

Marcel, S., R. Sawers, E. Oakeley, H. Angliker, and U. Paszkowski. 2010. “Tissue-Adapted Invasion Strategies of the Rice Blast Fungus Magnaporthe Oryzae.” The Plant Cell 22 (9): 3177–87.

Matasci, N., L. H. Hung, Z. Yan, E. J. Carpenter, N. J. Wickett, S. Mirarab, N. Nguyen, et al. 2014. “Data Access for the 1,000 Plants (1KP) Project.” GigaScience 3: 1–17. eCollection 2014.

Meyers, Blake C., Alexander Kozik, Alyssa Griego, Hanhui Kuang, and Richard W. Michelmore. 2003. “Genome-Wide Analysis of NBS-LRR–Encoding Genes in Arabidopsis.” The Plant Cell. https://doi.org/10.1105/tpc.009308.

Michael, Todd P., Douglas Bryant, Ryan Gutierrez, Nikolai Borisjuk, Philomena Chu, Hanzhong Zhang, Jing Xia, et al. 2017. “Comprehensive Definition of Genome Features in Spirodela Polyrhiza by High-Depth Physical Mapping and Short-Read DNA Sequencing Strategies.” The Plant Journal: For Cell and Molecular Biology 89 (3): 617–35.

Moeder, W., H. Ung, S. Mosher, and K. Yoshioka. 2010. “SA-ABA Antagonism in Defense Responses.” Plant Signaling & Behavior 5 (10): 1231–33.

Narusaka, Mari, Katsunori Hatakeyama, Ken Shirasu, and Yoshihiro Narusaka. 2014. “Arabidopsis Dual Resistance Proteins, Both RPS4 and RRS1, Are Required for Resistance to Bacterial Wilt in Transgenic Brassica Crops.” Plant Signaling & Behavior 9 (7): e29130.

Nicol, John W., Gregg A. Helt, Steven G. Blanchard, Archana Raja, and Ann E. Loraine. 2009. “The Integrated Genome Browser: Free Software for Distribution and Exploration of Genome-Scale Datasets.” Bioinformatics 25 (20): 2730–31.

Olsen, Jeanine L., Pierre Rouzé, Bram Verhelst, Yao-Cheng Lin, Till Bayer, Jonas Collen, Emanuela Dattolo, et al. 2016. “The Genome of the Seagrass Zostera Marina Reveals Angiosperm Adaptation to the Sea.” Nature 530: 331.

One Thousand Plant Transcriptomes Initiative. 2019. “One Thousand Plant Transcriptomes and the Phylogenomics of Green Plants.” Nature, October. https://doi.org/10.1038/s41586-019-1693-2.

Ortiz, Diana, Karine de Guillen, Stella Cesari, Véronique Chalvon, Jérome Gracy, André Padilla, and Thomas Kroj. 2017. “Recognition of the Magnaporthe Oryzae Effector AVR-Pia by the Decoy Domain of the Rice NLR Immune Receptor RGA5.” The Plant Cell 29 (1): 156–68.

Pandey, Sona, Rui-Sheng Wang, Liza Wilson, Song Li, Zhixin Zhao, Timothy E. Gookin, Sarah M. Assmann, and Réka Albert. 2010. “Boolean Modeling of Transcriptome Data Reveals Novel Modes of Heterotrimeric G-Protein Action.” Molecular Systems Biology 6 (1). https://www.embopress.org/doi/abs/10.1038/msb.2010.28.

Qi, Tiancong, Kyungyong Seong, Daniela P. T. Thomazella, Joonyoung Ryan Kim, Julie Pham, Eunyoung Seo, Myeong-Je Cho, Alex Schultink, and Brian J. Staskawicz. 2018. “NRG1 Functions Downstream of EDS1 to Regulate TIR-NLR-Mediated Plant Immunity in Nicotiana Benthamiana.” Proceedings of the National Academy of Sciences of the United States of America 115 (46): E10979.

Radhakrishnan, Guru V., Jean Keller, Melanie K. Rich, Tatiana Vernié, Duchesse L. Mbadinga Mbaginda, Nicolas Vigneron, Ludovic Cottret, et al. 2019. “An Ancestral Signalling Pathway Is Conserved in Plant Lineages Forming Intracellular Symbioses.” bioRxiv. https://doi.org/10.1101/804591.

Ramegowda, Venkategowda, and Muthappa Senthil-Kumar. 2015. “The Interactive Effects of Simultaneous Biotic and Abiotic Stresses on Plants: Mechanistic Understanding from Drought and Pathogen Combination.” Journal of Plant Physiology 176: 47–54.

Ryals, J., K. Weymann, K. Lawton, L. Friedrich, D. Ellis, H. Y. Steiner, J. Johnson, et al. 1997. “The Arabidopsis NIM1 Protein Shows Homology to the Mammalian Transcription Factor Inhibitor I Kappa B.” The Plant Cell 9 (3): 425–39.

Sanderson, Michael J., Jeffrey L. Thorne, Niklas Wikström, and Kåre Bremer. 2004. “Molecular Evidence on Plant Divergence Times.” American Journal of Botany 91 (10): 1656–65.

Sarris, Panagiotis F., Volkan Cevik, Gulay Dagdas, Jonathan D. G. Jones, and Ksenia V. Krasileva. 2016. “Comparative Analysis of Plant Immune Receptor Architectures Uncovers Host Proteins Likely Targeted by Pathogens.” BMC Biology 14 (1): 8.

Saucet, Simon B., Yan Ma, Panagiotis F. Sarris, Oliver J. Furzer, Kee Hoon Sohn, and Jonathan D. G. Jones. 2015. “Two Linked Pairs of Arabidopsis TNL Resistance Genes Independently Confer Recognition of Bacterial Effector AvrRps4.” Nature Communications 6 (March): 6338.

Shah, Jyoti, Frank Tsui, and Daniel F. Klessig. 1997. “Characterization of a S Alicylic a Cid-I Nsensitive Mutant (sai1) of Arabidopsis Thaliana, Identified in a Selective Screen Utilizing the SA-Inducible Expression of the tms2 Gene.” Molecular Plant-Microbe Interactions: MPMI 10 (1): 69–78.

Simão, Felipe A., Robert M. Waterhouse, Panagiotis Ioannidis, Evgenia V. Kriventseva, and Evgeny M. Zdobnov. 2015. “BUSCO: Assessing Genome Assembly and Annotation Completeness with Single-Copy Orthologs.” Bioinformatics 31 (19): 3210–12.

Stamatakis, Alexandros. 2014. “RAxML Version 8: A Tool for Phylogenetic Analysis and Post-Analysis of Large Phylogenies.” Bioinformatics 30 (9): 1312–13.

Stam, Remco, Gustavo Silva-Arias, and Aurelien Tellier. 2019. “Subsets of NLR Genes Show Differential Signatures of Adaptation during Colonization of New Habitats.” The New Phytologist 224 (1): 367–79.

Steuernagel, Burkhard, Florian Jupe, Kamil Witek, Jonathan D. G. Jones, and Brande B. H. Wulff. 2015. “NLR-Parser: Rapid Annotation of Plant NLR Complements.” Bioinformatics 31 (10): 1665–67.

Steuernagel, Burkhard, Kamil Witek, Simon G. Krattinger, Ricardo H. Ramirez-Gonzalez, Henk-Jan Schoonbeek, Guotai Yu, Erin Baggs, et al. 2018. “Physical and Transcriptional Organisation of the Bread Wheat Intracellular Immune Receptor Repertoire.” https://repository.kaust.edu.sa/handle/10754/628448.

Stolzer, M., Lai, H., Xu, M., Sathaye, D., Vernot, B., and Durand, D. (2012). Inferring duplications, losses, transfers and incomplete lineage sorting with nonbinary species trees. Bioinformatics 28: i409–i415

Thanki, Anil S., Nicola Soranzo, Wilfried Haerty, and Robert P. Davey. 2018. “GeneSeqToFamily: A Galaxy Workflow to Find Gene Families Based on the Ensembl Compara GeneTrees Pipeline.” GigaScience 7 (3): 1–10.

Torres Zabala, Marta de, Mark H. Bennett, William H. Truman, and Murray R. Grant. 2009. “Antagonism between Salicylic and Abscisic Acid Reflects Early Host--Pathogen Conflict and Moulds Plant Defence Responses.” The Plant Journal: For Cell and Molecular Biology 59 (3): 375–86.

VanBuren, Robert, Doug Bryant, Patrick P. Edger, Haibao Tang, Diane Burgess, Dinakar Challabathula, Kristi Spittle, et al. 2015. “Single-Molecule Sequencing of the Desiccation-Tolerant Grass Oropetium Thomaeum.” Nature 527: 508.

VanBuren, Robert, Ching Man Wai, Jens Keilwagen, and Jeremy Pardo. 2018. “A Chromosome Scale Assembly of the Model Desiccation Tolerant Grass Oropetium Thomaeum.” bioRxiv, 378943.

Van de Weyer, Anna-Lena, Freddy Monteiro, Oliver J. Furzer, Marc T. Nishimura, Volkan Cevik, Kamil Witek, Jonathan D. G. Jones, Jeffery L. Dangl, Detlef Weigel, and Felix Bemm. 2019. “The Arabidopsis Thaliana Pan-NLRome.” bioRxiv. https://doi.org/10.1101/537001.

Van Ghelder, Cyril, Geneviève J. Parent, Philippe Rigault, Julien Prunier, Isabelle Giguère, Sé Caron, Juliana Stival Sena, et al. 2019. “The Large Repertoire of Conifer NLR Resistance Genes Includes Drought Responsive and Highly Diversified RNLs.” Scientific Reports 9 (1): 11614.

Velzen, Robin van, Rens Holmer, Fengjiao Bu, Luuk Rutten, Arjan van Zeijl, Wei Liu, Luca Santuari, et al. 2018. “Comparative Genomics of the Nonlegume Parasponia Reveals Insights into Evolution of Nitrogen-Fixing Rhizobium Symbioses.” Proceedings of the National Academy of Sciences 115 (20): E4700–4709.

Venugopal, Srivathsa C., Rae-Dong Jeong, Mihir K. Mandal, Shifeng Zhu, A. Chandra-Shekara, Ye Xia, Matthew Hersh, et al. 2009. “Enhanced Disease Susceptibility 1 and Salicylic Acid Act Redundantly to Regulate Resistance Gene-Mediated Signaling.” PLoS Genetics 5 (7): e1000545.

Wagner, Stephan, Johannes Stuttmann, Steffen Rietz, Raphael Guerois, Elena Brunstein, Jaqueline Bautor, Karsten Niefind, and Jane E. Parker. 2013. “Structural Basis for Signaling by Exclusive EDS1 Heteromeric Complexes with SAG101 or PAD4 in Plant Innate Immunity.” Cell Host & Microbe 14 (6): 619–30.

Wang, Di, Yajiao Pan, Xiuqin Zhao, Linghua Zhu, Binying Fu, and Zhikang Li. 2011. “Genome-Wide Temporal-Spatial Gene Expression Profiling of Drought Responsiveness in Rice.” BMC Genomics 12 (March): 149.

Wang, Guan-Feng, Jiabing Ji, Farid El Kasmi, Jeffery Dangl, Guri Johal, and Peter Balint-Kurti. 2015. “Molecular and Functional Analyses of a Maize Autoactive NB-LRR Protein Identify Precise Structural Requirements for Activity.” PLoS Pathogens 11: e1004674.

Wang, Wenguo, Rui Li, Qili Zhu, Xiaoyu Tang, and Qi Zhao. 2016. “Transcriptomic and Physiological Analysis of Common Duckweed Lemna Minor Responses to NH4+ Toxicity.” BMC Plant Biology 16 (1): 92.

Wang, W., G. Haberer, H. Gundlach, C. Gläßer, T. Nussbaumer, M. C. Luo, A. Lomsadze, et al. 2014. “The Spirodela Polyrhiza Genome Reveals Insights into Its Neotenous Reduction Fast Growth and Aquatic Lifestyle.” Nature Communications 5: 3311.

Wan, Li, Kow Essuman, Ryan G. Anderson, Yo Sasaki, Freddy Monteiro, Eui-Hwan Chung, Erin Osborne Nishimura, et al. 2019. “TIR Domains of Plant Immune Receptors Are NAD -Cleaving Enzymes That Promote Cell Death.” Science. https://doi.org/10.1126/science.aax1771.

Waterhouse, Andrew M., James B. Procter, David M. A. Martin, Michèle Clamp, and Geoffrey J. Barton. 2009. “Jalview Version 2—a Multiple Sequence Alignment Editor and Analysis Workbench.” Bioinformatics 25 (9): 1189–91.

Wen, Zhifeng, Liping Yao, Stacy D. Singer, Hanif Muhammad, Zhi Li, and Xiping Wang. 2017. “Constitutive Heterologous Overexpression of a TIR-NB-ARC-LRR Gene Encoding a Putative Disease Resistance Protein from Wild Chinese Vitis Pseudoreticulata in Arabidopsis and Tobacco Enhances Resistance to Phytopathogenic Fungi and Bacteria.” Plant Physiology and Biochemistry: PPB / Societe Francaise de Physiologie Vegetale 112: 346–61.

Wheeler, Travis J., and Sean R. Eddy. 2013. “Nhmmer: DNA Homology Search with Profile HMMs.” Bioinformatics 29 (19): 2487–89.

Wickett, N. J., S. Mirarab, N. Nguyen, T. Warnow, E. Carpenter, N. Matasci, S. Ayyampalayam, et al. 2014. “Phylotranscriptomic Analysis of the Origin and Early Diversification of Land Plants.” Proceedings of the National Academy of Sciences of the United States of America 111 (45): 4859.

Wu, Chih-Hang, Ahmed Abd-El-Haliem, Tolga O. Bozkurt, Khaoula Belhaj, Ryohei Terauchi, Jack H. Vossen, and Sophien Kamoun. 2017. “NLR Network Mediates Immunity to Diverse Plant Pathogens.” Proceedings of the National Academy of Sciences of the United States of America 114 (30): 8113–18.

Wu, Chih-Hang, Lida Derevnina, and Sophien Kamoun. 2018. “Receptor Networks Underpin Plant Immunity.” Science 360 (6395): 1300–1301.

Wu, Z., M. Li, O. X. Dong, S. Xia, W. Liang, Y. Bao, G. Wasteneys, and X. Li. 2019. “Differential Regulation of TNL-Mediated Immune Signaling by Redundant Helper CNLs.” The New Phytologist 222 (2): 938–53.

Xu, Guoyong, Meng Yuan, Chaoren Ai, Lijing Liu, Edward Zhuang, Sargis Karapetyan, Shiping Wang, and Xinnian Dong. 2017. “uORF-Mediated Translation Allows Engineered Plant Disease Resistance without Fitness Costs.” Nature 545 (7655): 491–94.

Yu, Yanhua, Jana Streubel, Sandrine Balzergue, Antony Champion, Jens Boch, Ralf Koebnik, Jiaxun Feng, Valérie Verdier, and Boris Szurek. 2011. “Colonization of Rice Leaf Blades by an African Strain of Xanthomonas Oryzae Pv. Oryzae Depends on a New TAL Effector That Induces the Rice Nodulin-3 Os11N3 Gene.” Molecular Plant-Microbe Interactions. https://doi.org/10.1094/mpmi-11-10-0254.

Zeng, Liping, Qiang Zhang, Renran Sun, Hongzhi Kong, Ning Zhang, and Hong Ma. 2014. “Resolution of Deep Angiosperm Phylogeny Using Conserved Nuclear Genes and Estimates of Early Divergence Times.” Nature Communications 5 (1): 4956.

Zhong, Bojian, Linhua Sun, and David Penny. 2015. “The Origin of Land Plants: A Phylogenomic Perspective.” Evolutionary Bioinformatics Online 11: EBO.S29089.

Zhu, Shifeng, Rae-Dong Jeong, Srivathsa C. Venugopal, Ludmila Lapchyk, Duroy Navarre, Aardra Kachroo, and Pradeep Kachroo. 2011. “SAG101 Forms a Ternary Complex with EDS1 and PAD4 and Is Required for Resistance Signaling against Turnip Crinkle Virus.” PLoS Pathogens 7 (11): e1002318.

