## Supplementary material for "Convergent loss of an EDS1/PAD4 signalling pathway in several plant lineages predicts new components of plant immunity and drought response": Supplemetal Figures

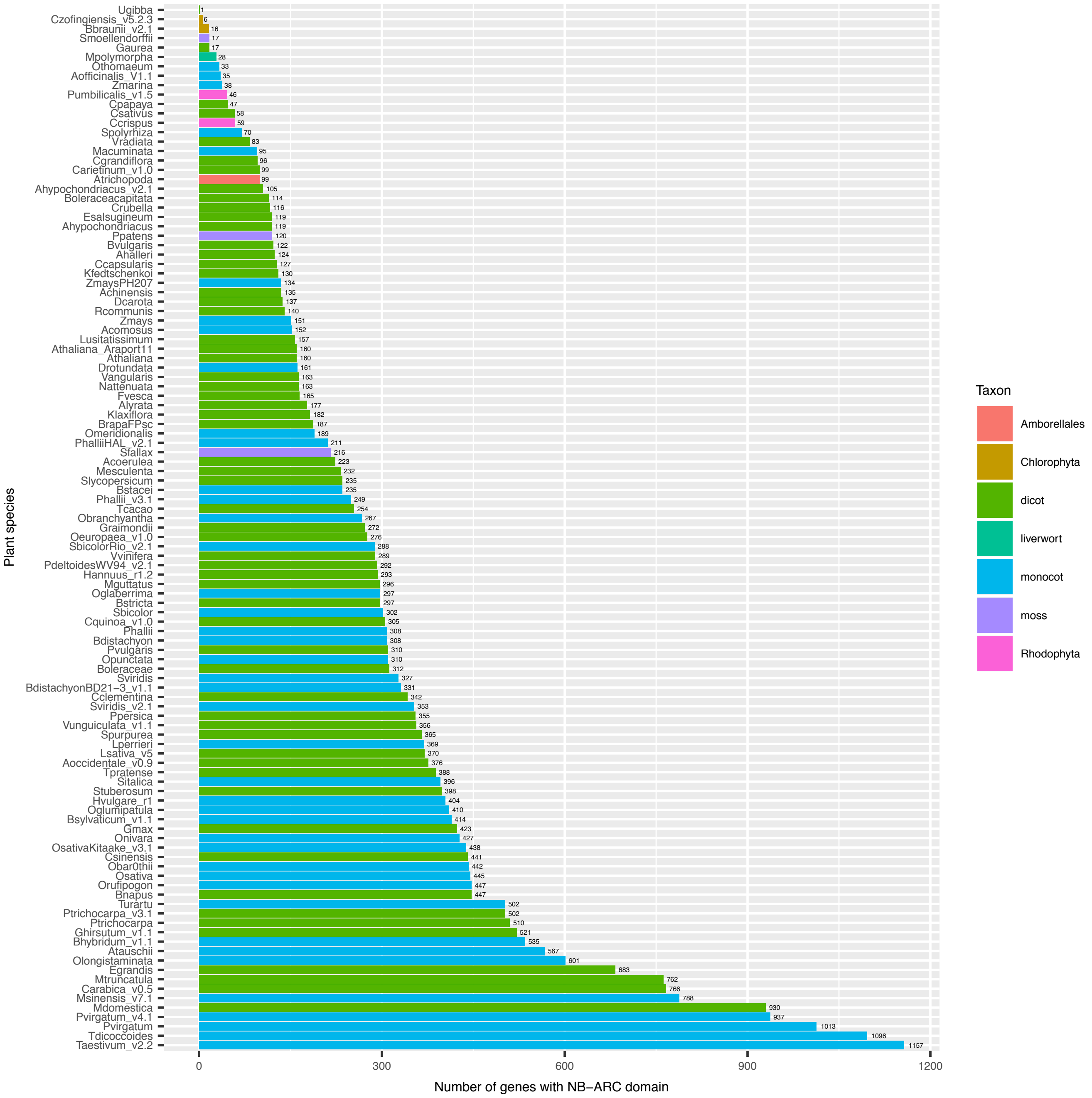


Supplementary fig 1 - Histogram of the number of genes with NB-ARC domains for 109 available plant genomes. Colour of bars indicates taxon and number at the end of bar is the number of NB-ARC domains identified. Labels of species correspond to given name of genome in database from which downloaded.

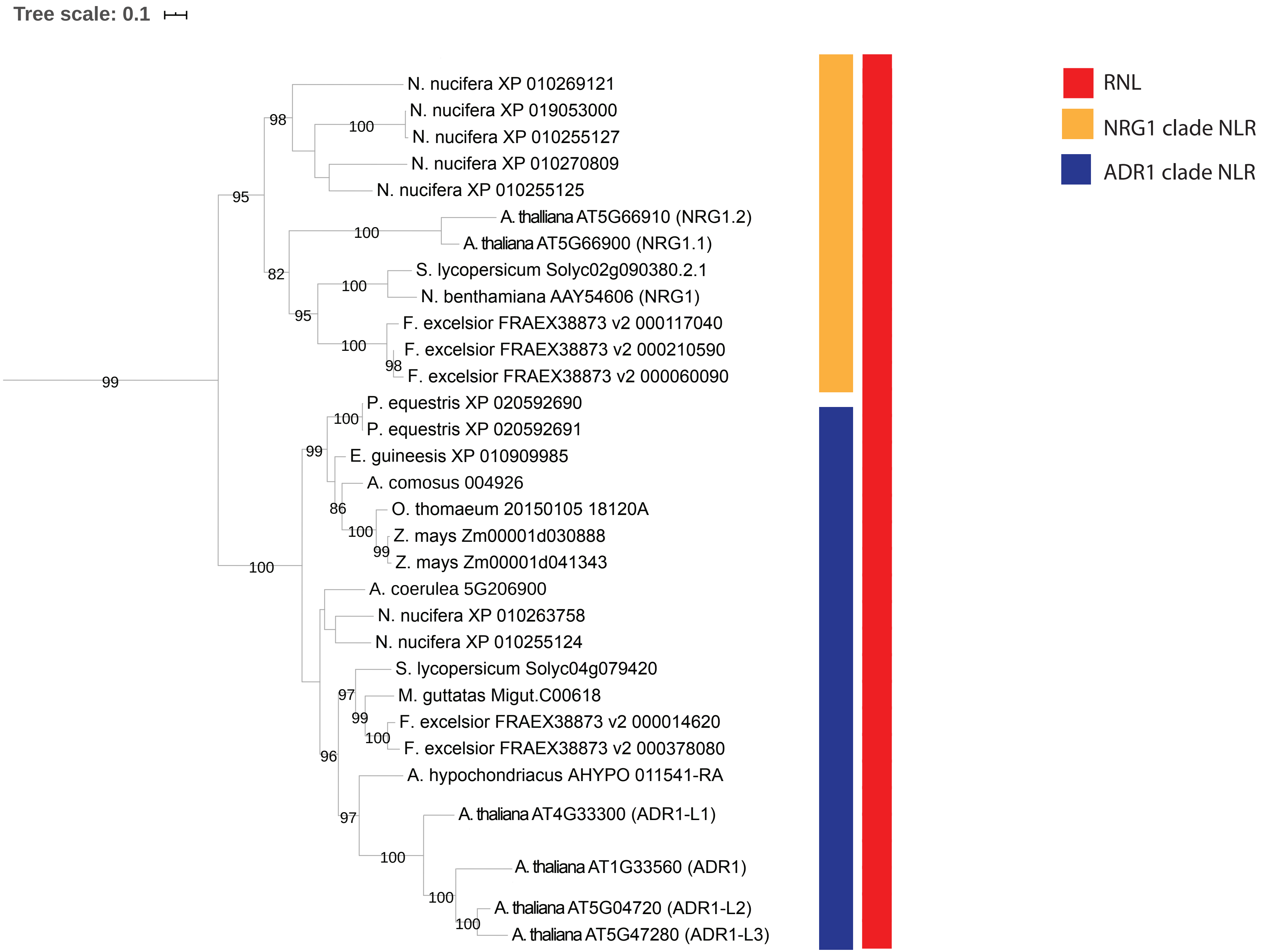


Supplementary fig 2 - Maximum likelihood phylogeny of the NB-ARC domain of RNL genes across the 18 representative species. Number on branch indicates bootstrap value. Leafs are given as species followed by gene identifier. Red bar indicates RNL type NLR whilst Orange and blue indicate clades NRG1 and ADR1 respectively.

Table 1

| Species | PBS1 | RIN4 | ICS2 | SGT1 | NPR1 | NPR2 | EDS1 | PAD4 | SAG101 | ADR1-L1 (NRG1) | SID1 | SID2/ICS1 | FMO1 |
|---|---|---|---|---|---|---|---|---|---|---|---|---|---|
| | AT5G13160 | AT3G25070 | AT1G18870 | AT2G43820 | AT1G64280 | AT4G26120 | AT3G48090/80 | AT3G52430 | AT5G14930 | AT4G33300 | AT4G39030 | AT1G74710 | AT1G19250 |
| Amborella trichopoda | | | | | | | | | | | | | |
| Eelgrass | | | | | | | | | | | | | |
| Duckweed | | | | | | | | | | | | | |
| Orchid | | | | | | | | | | | | | |
| Oil palm | | | | | | | | | | | | | |
| Pineapple | | | | | | | | | | | | | |
| Rice | | | | ?- fused gene model to neighbour Arab A2G43840??? | | | | | | | | | |
| Resurrection grass | | | | | | | Annotated as fused to neighbour protein | | | | | | |
| Maize | | | | | | | | | | | | | |
| A. coerulea | | | | | | | | | | | | | |
| Lotus | | | | | | | | | | | | | |
| A. thaliana | | | | | | | | | | | | | |
| Amaranthus | | | | | | | | | | | | | |
| Tomato | | | | | | | | | | | | | |
| Ash | | | | | | | | | | | | | |
| Monkey Flower | | | | | Top hit on [illegible] | | | | | | | | |
| Humped Bladderwort | | | | | | | | | | | | | |
| Corkscrew Flower | | | | | NPR1 like BOP2 | | | | | | | | |

| Species | ALD1 | MC1 | MC4 | BIK1 | PBL1 | PBL9 | PBL10 | PBL11 | MPK4 | VIP1 | DTX42 | DTX46 | YUC3 |
|---|---|---|---|---|---|---|---|---|---|---|---|---|---|
| | AT2G13810 | AT1G02170 | AT1G79340 | AT2G39660 | AT3G55450 | AT1G07570 | AT2G28930 | AT5G02290 | AT4G01370 | AT1G43700 | AT1G51340 | AT2G21340 | AT1G04610 |
| Amborella trichopoda | | | | | | | | | | | | | |
| Eelgrass | | | | | | | | | | | | | |
| Duckweed | | | | | | | | | | | | | |
| Orchid | | | | | | | | | | | | | |
| Oil palm | | | ?MC5 neighbour | | | | | | | | | | |
| Pineapple | | | | | | | | | | | | | |
| Rice | | | ?MC5 neighbour | | | | | | | | | | |
| Resurrection grass | | | | | | | | | | | | | |
| Maize | | | ?MC5 neighbour | | | | | | | | | | |
| A. coerulea | | | | | | | | | | | | | |
| Lotus | | | | | | | | | | | | | |
| A. thaliana | | | | | | | | | | | | | |
| Amaranthus | | | ?MC5 neighbour | | | | | | | | | | |
| Tomato | | | | | | | | | | | | | |
| Ash | | | | | | | | | | | | | |
| Monkey Flower | | | | | | | | | | | | | |
| Humped Bladderwort | | | | | | | | | | | | | |
| Corkscrew Flower | | | | | | | | | | | | | |

Supplementary fig 3 -Extended species plot of presence and absence of known immunity genes from reciprocal BLAST searches .
Red indicates the absence of an identified ortholog by reciprocal blast whilst green represence presence. Gene names given as in Arabidopsis thaliana.

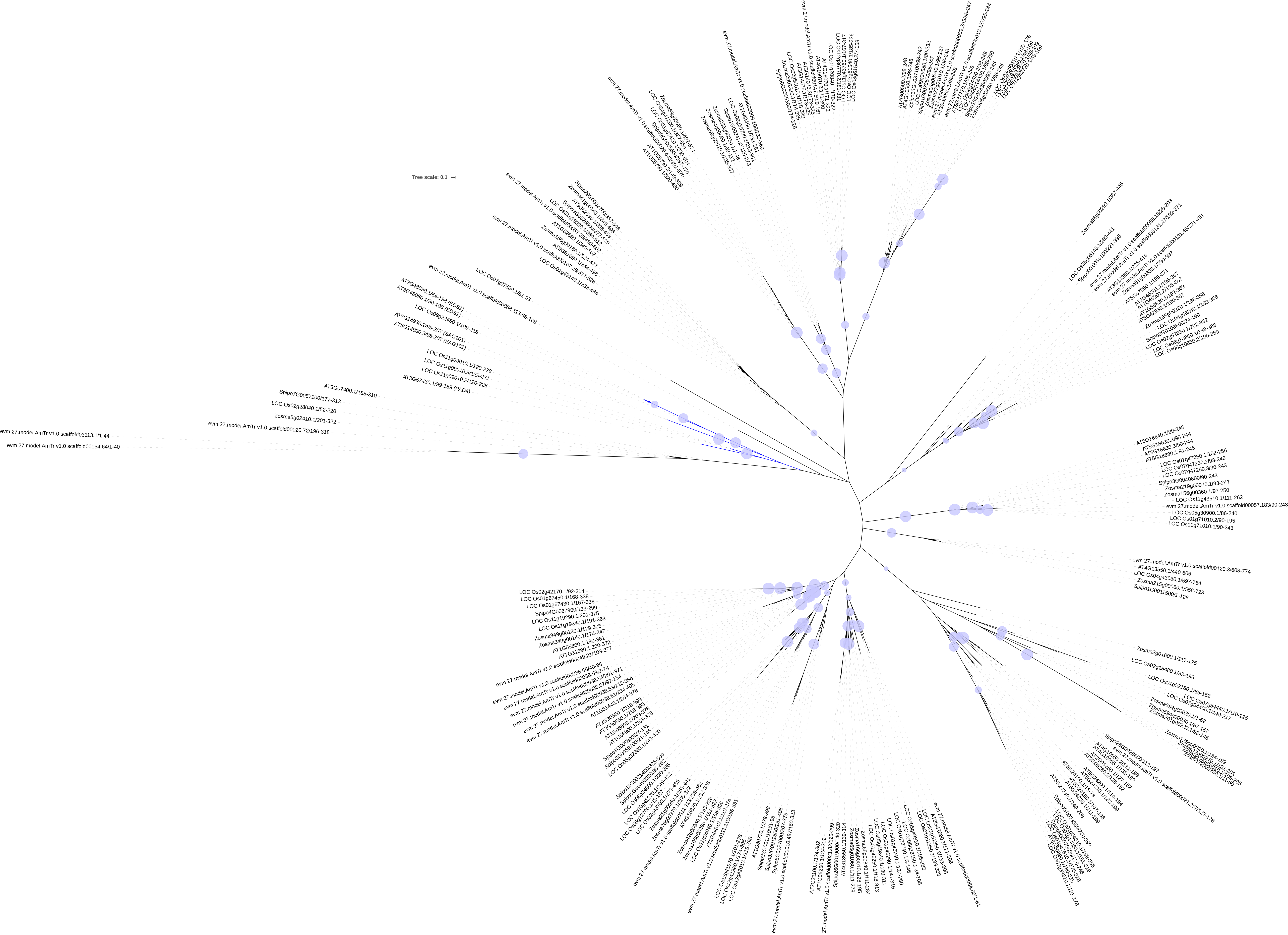


Supplementary fig 4 - Maximum-likelihood gene tree from proteins containing lipase-3 HMM domain.
Blue branches highlight the clade of EDS1/PAD4/SAG101 type lipase 3 motif proteins. Blue dots represent bootstrap values >80. Gene identifiers given as leaves.

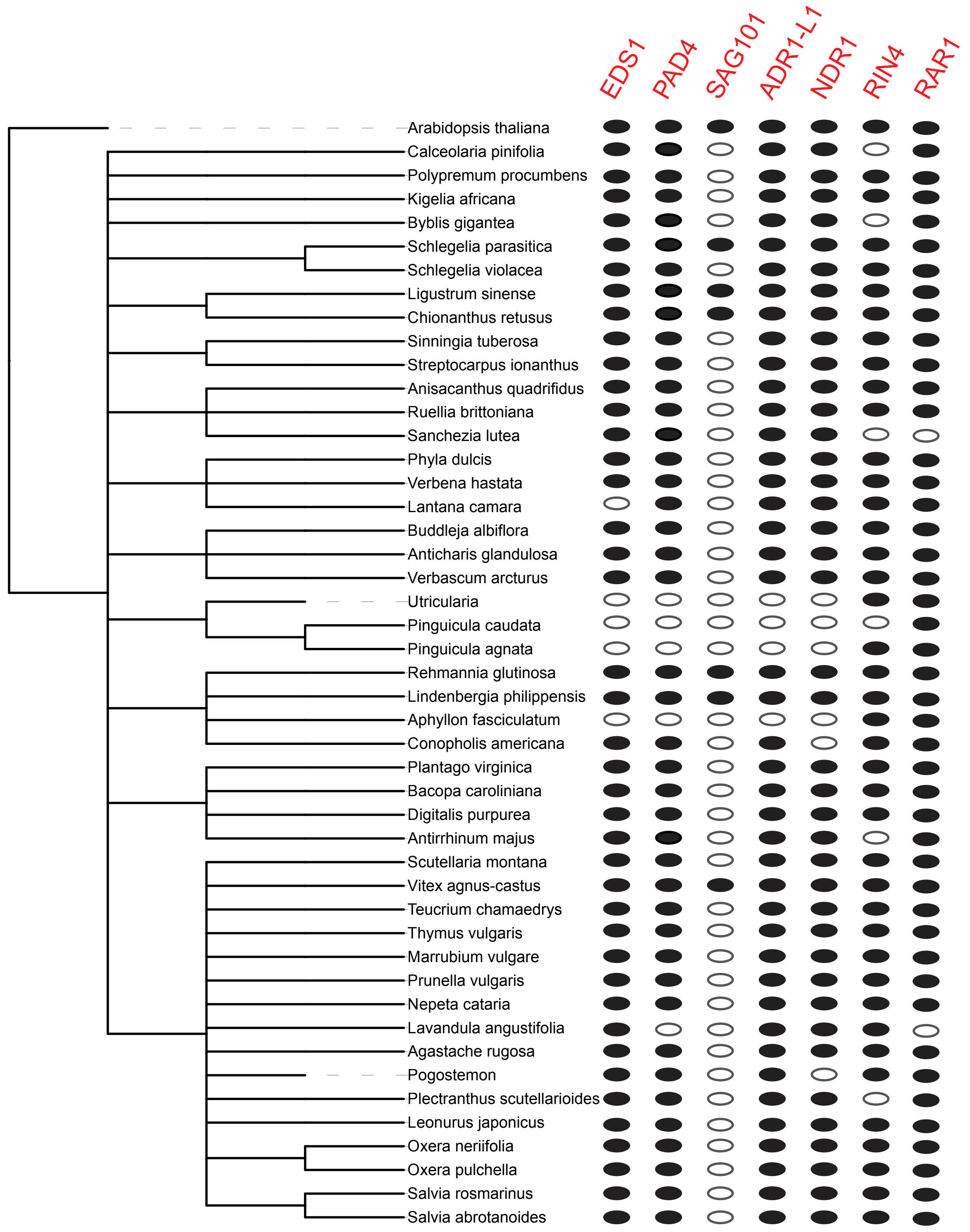


Supplementary fig 5 - Plot of presence and absence of 7 characterised ETI components across transcriptomes of dicots and magnoliids.
Gene names correspond to those from Arabidopsis thaliana. Phylogeny from NCBI taxonomy with species name at leaves. Black filled circle indicate identified reciprocal blast ortholog to arabidopsis query in transcriptome. Grey outlined circles indicate no reciprocal best blast hit in available transcriptome.

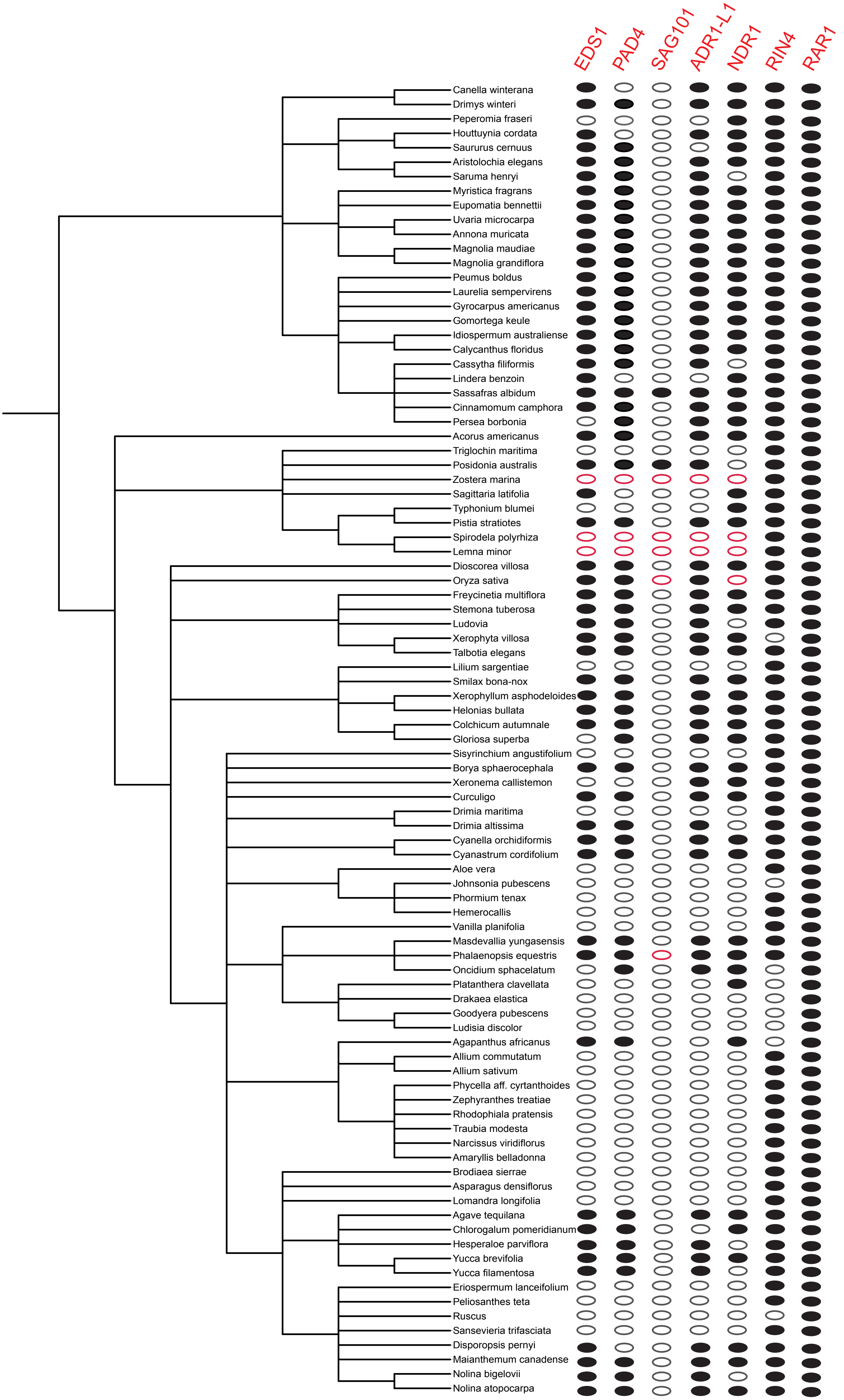


Supplementary fig 6 - Plot of presence and absence of 7 characterised ETI components across transcriptomes of monocots. Gene names correspond to those from Arabidopsis thaliana. Phylogeny from NCBI taxonomy with species name at leaves. Black filled circle indicate identified reciprocal blast ortholog to arabidopsis or rice query in transcriptome. Grey outlined circles indicate no reciprocal best blast hit in available transcriptome. Red outlined circles indicate absence in transcriptome and genome.

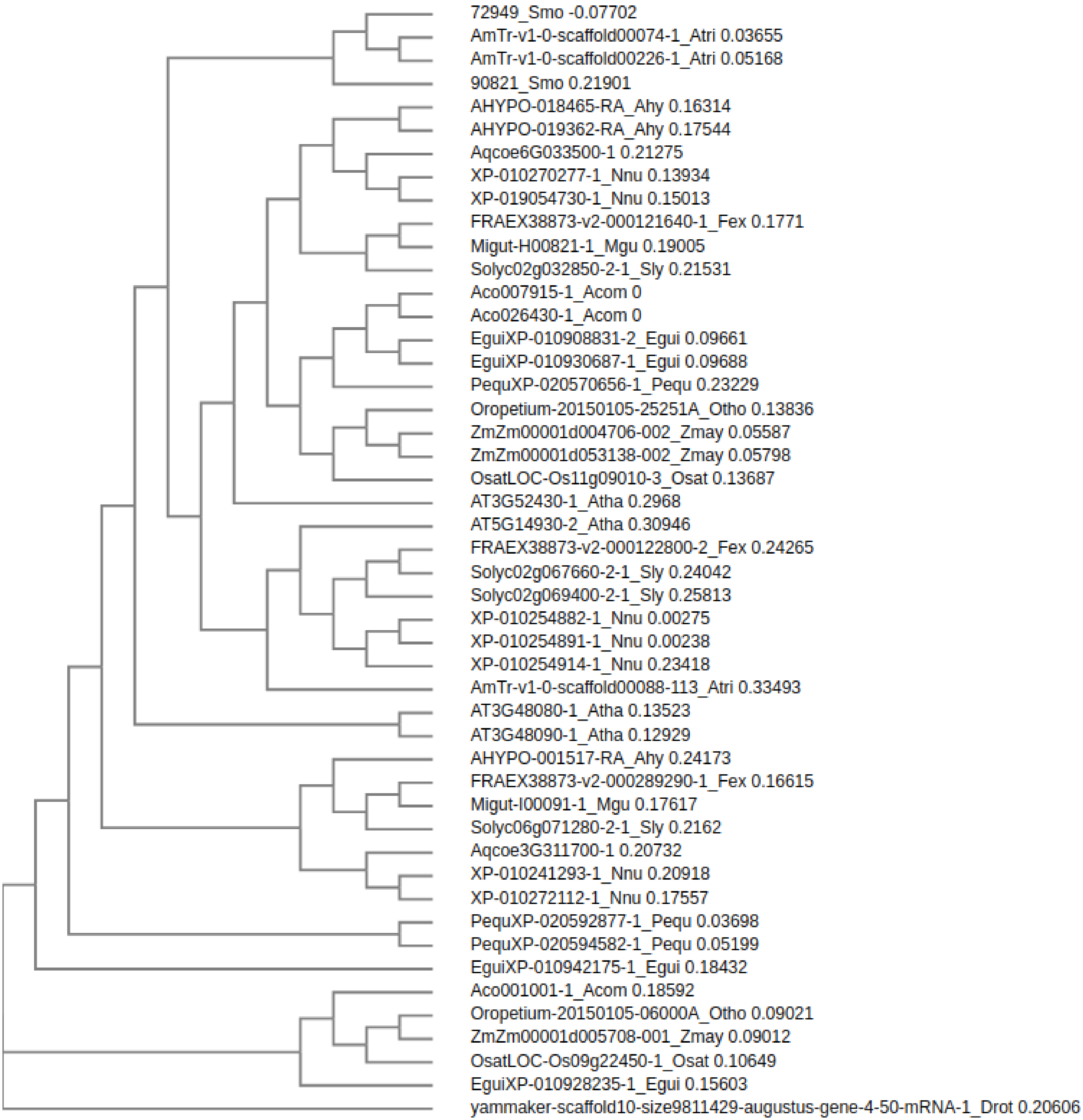


Supplementary fig 7 - Gene tree for EDS1, PAD4 and SAG101 gene family from orthologs identified from GeneSeqToFamily.

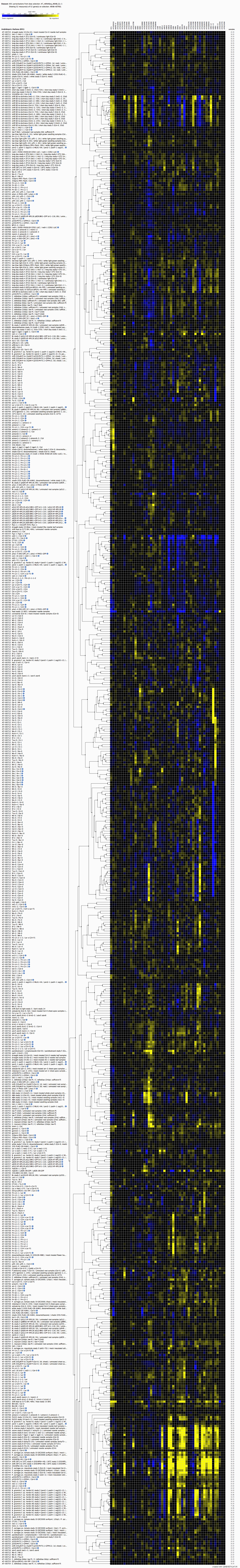

Supplementary fig 8 - Heatmap of differential expression of ASTREL genes across publicly available RNAseq experiments using Geneinvestigator.

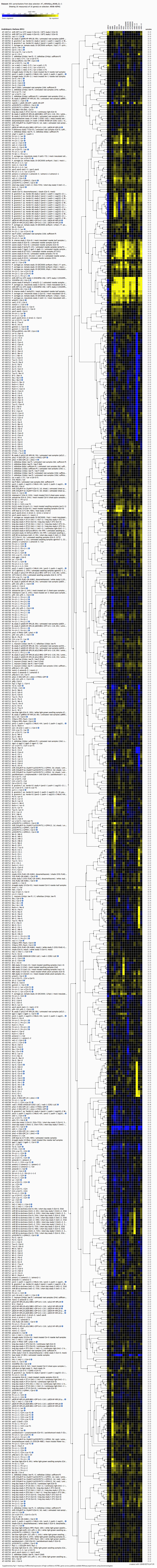

Supplementary fig 9 - Heatmap of differential expression of high confidence ASTREL genes across publicly available RNAseq experiments using Geneinvestigator.

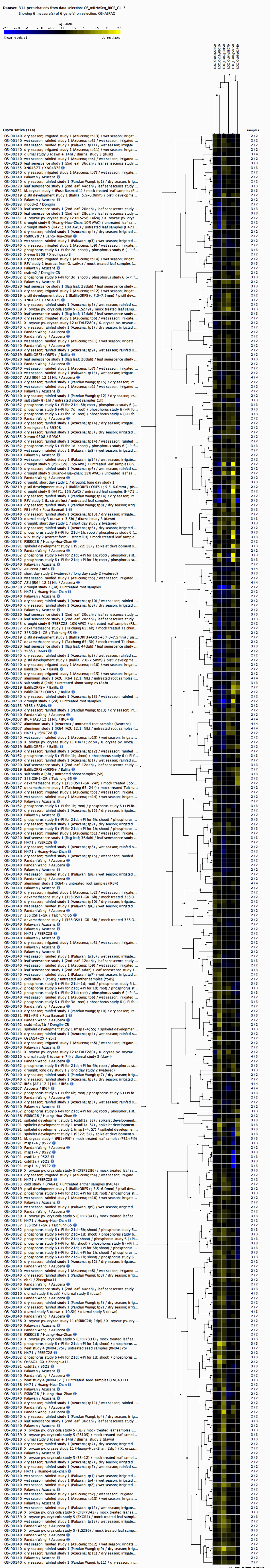


Supplementary fig 10 - Heatmap of differential expression of high confidence ASTREL genes across publicly available RNAseq experiments using Geneinvestigator.

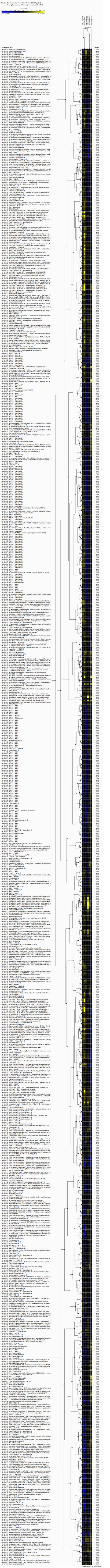


Supplementary fig 11 - Heatmap of differential expression of high confidence ASTREL genes across publicly available microarray experiments using Geneinvestigato[illegible]
