## Supplementary material for "Convergent loss of an EDS1/PAD4 signalling pathway in several plant lineages predicts new components of plant immunity and drought response": Supplemetal Tables

Table S1

| PF00931 | Species_abbreviated | Species | Taxon | Database |
|---|---|---|---|---|
| 223 | Acoerulea | Aquilegia coerulea | dicot | phytozome12.1.6 |
| 152 | Acomosus | Ananas comosus | monocot | phytozome12.1.6 |
| 124 | Ahalleri | Arabidopsis halleri | dicot | phytozome12.1.6 |
| 119 | Ahypochondriacus | Amaranthus hypochondriacus | dicot | phytozome12.1.6 |
| 105 | Ahypochondriacus_v2.1 | Amaranthus hypochondriacus | dicot | phytozome12.1.6 |
| 177 | Alyrata | Arabidopsis lyrata | dicot | phytozome12.1.6 |
| 376 | Aoccidentale_v0.9 | Anacardium occidentale | dicot | phytozome12.1.6 |
| 35 | Aofficinalis_V1.1 | Asparagus officinalis | monocot | phytozome12.1.6 |
| 160 | Athaliana | Arabidopsis thaliana columbia | dicot | phytozome12.1.6 |
| 160 | Athaliana_Araport11 | Arabidopsis thaliana columbia | dicot | phytozome12.1.6 |
| 99 | Atrichopoda | Amborella trichopoda | Amborellales | phytozome12.1.6 |
| 16 | Bbraunii_v2.1 | Botryococcus braunii | Chlorophyta | phytozome12.1.6 |
| 308 | Bdistachyon | Brachypodium distachyon | monocot | phytozome12.1.6 |
| 331 | BdistachyonBD21-3_v1.1 | Brachypodium distachyon Bd21-3 | monocot | phytozome12.1.6 |
| 535 | Bhybridum_v1.1 | Brachypodium hybridum | monocot | phytozome12.1.6 |
| 114 | Boleraceacapitata | Brassica oleracea capitata | dicot | phytozome12.1.6 |
| 187 | BrapaFPsc | Brassica rapa FPsc | dicot | phytozome12.1.6 |
| 235 | Bstacei | Brachypodium stacei | monocot | phytozome12.1.6 |
| 297 | Bstricta | Boechera stricta | dicot | phytozome12.1.6 |
| 414 | Bsylvaticum_v1.1 | Brachypodium sylvaticum | monocot | phytozome12.1.6 |
| 766 | Carabica_v0.5 | Coffea arabica | dicot | phytozome12.1.6 |
| 99 | Carietinum_v1.0 | Cicer arietinum | dicot | phytozome12.1.6 |
| 342 | Cclementina | Citrus clementina | dicot | phytozome12.1.6 |
| 96 | Cgrandiflora | Capsella grandiflora | dicot | phytozome12.1.6 |
| 47 | Cpapaya | Carica papaya | dicot | phytozome12.1.6 |
| 305 | Cquinoa_v1.0 | Chenopodium quinoa | dicot | phytozome12.1.6 |
| 116 | Crubella | Capsella rubella | dicot | phytozome12.1.6 |
| 58 | Csativus | Cucumis sativus | dicot | phytozome12.1.6 |
| 441 | Csinensis | Citrus sinensis | dicot | phytozome12.1.6 |
| 6 | Czofingiensis_v5.2.3 | Chromochloris zofingiensis | Chlorophyta | phytozome12.1.6 |
| 137 | Dcarota | Daucus carota | dicot | phytozome12.1.6 |
| 683 | Egrandis | Eucalyptus grandis | dicot | phytozome12.1.6 |
| 119 | Esalsugineum | Eutrema salsugineum | dicot | phytozome12.1.6 |
| 165 | Fvesca | Fragaria vesca | dicot | phytozome12.1.6 |
| 521 | Ghirsutum_v1.1 | Gossypium hirsutum | dicot | phytozome12.1.6 |
| 423 | Gmax | Glycine max | dicot | phytozome12.1.6 |
| 272 | Graimondii | Gossypium raimondii | dicot | phytozome12.1.6 |
| 293 | Hannuus_r1.2 | Helianthus annuus | dicot | phytozome12.1.6 |
| 404 | Hvulgare_r1 | Hordeum vulgare | monocot | phytozome12.1.6 |
| 130 | Kfedtschenkoi | Kalanchoe fedtschenkoi | dicot | phytozome12.1.6 |
| 182 | Klaxiflora | Kalanchoe laxiflora | dicot | phytozome12.1.6 |
| 370 | Lsativa_v5 | Lactuca sativa | dicot | phytozome12.1.6 |
| 157 | Lusitatissimum | Linum usitatissimum | dicot | phytozome12.1.6 |
| 95 | Macuminata | Musa acuminata | monocot | phytozome12.1.6 |
| 930 | Mdomestica | Malus domestica | dicot | phytozome12.1.6 |

| | | | | |
|---|---|---|---|---|
| 232 | Mesculenta | Manihot esculenta | dicot | phytozome12.1.6 |
| 296 | Mguttatus | Mimulus guttatus | dicot | phytozome12.1.6 |
| 28 | Mpolymorpha | Marchantia polymorpha | liverwort | phytozome12.1.6 |
| 788 | Msinensis_v7.1 | Miscanthus sinensis | monocot | phytozome12.1.6 |
| 762 | Mtruncatula | Medicago truncatula | dicot | phytozome12.1.6 |
| 276 | Oeuropaea_v1.0 | Olea europaea var. sylvestris | dicot | phytozome12.1.6 |
| 445 | Osativa | Oryza sativa | monocot | phytozome12.1.6 |
| 438 | OsativaKitaake_v3.1 | Oryza sativa Kitaake | monocot | phytozome12.1.6 |
| 33 | Othomaeum | Oropetium thomaeum | monocot | phytozome12.1.6 |
| 292 | PdeltoidesWV94_v2.1 | Populus deltoides WV94 | dicot | phytozome12.1.6 |
| 308 | Phallii | Panicum hallii | monocot | phytozome12.1.6 |
| 211 | PhalliiHAL_v2.1 | Panicum hallii var. hallii | monocot | phytozome12.1.6 |
| 249 | Phallii_v3.1 | Panicum hallii var. filipes | monocot | phytozome12.1.6 |
| 120 | Ppatens | Physcomitrella patens | moss | phytozome12.1.6 |
| 355 | Ppersica | Prunus persica | dicot | phytozome12.1.6 |
| 510 | Ptrichocarpa | Populus trichocarpa | dicot | phytozome12.1.6 |
| 502 | Ptrichocarpa_v3.1 | Populus trichocarpa | dicot | phytozome12.1.6 |
| 46 | Pumbilicalis_v1.5 | Porphyra umbilicalis | Rhodophyta | phytozome12.1.6 |
| 1013 | Pvirgatum | Panicum virgatum | monocot | phytozome12.1.6 |
| 937 | Pvirgatum_v4.1 | Panicum virgatum | monocot | phytozome12.1.6 |
| 310 | Pvulgaris | Phaseolus vulgaris | dicot | phytozome12.1.6 |
| 140 | Rcommunis | Ricinus communis | dicot | phytozome12.1.6 |
| 302 | Sbicolor | Sorghum bicolor | monocot | phytozome12.1.6 |
| 288 | SbicolorRio_v2.1 | Sorghum bicolor rio | monocot | phytozome12.1.6 |
| 216 | Sfallax | Sphagnum fallax | moss | phytozome12.1.6 |
| 396 | Sitalica | Setaria italica | monocot | phytozome12.1.6 |
| 235 | Slycopersicum | Solanum lycopersicum | dicot | phytozome12.1.6 |
| 17 | Smoellendorffii | Selaginella moellendorffii | moss | phytozome12.1.6 |
| 70 | Spolyrhiza | Spirodela polyrhiza | monocot | phytozome12.1.6 |
| 365 | Spurpurea | Salix purpurea | dicot | phytozome12.1.6 |
| 398 | Stuberosum | Solanum tuberosum | dicot | phytozome12.1.6 |
| 327 | Sviridis | Setaria viridis | monocot | phytozome12.1.6 |
| 353 | Sviridis_v2.1 | Setaria viridis | monocot | phytozome12.1.6 |
| 1157 | Taestivum_v2.2 | Triticum aestivum | monocot | phytozome12.1.6 |
| 254 | Tcacao | Theobroma cacao | dicot | phytozome12.1.6 |
| 388 | Tpratense | Trifolium pratense | dicot | phytozome12.1.6 |
| 356 | Vunguiculata_v1.1 | Vigna unguiculata | dicot | phytozome12.1.6 |
| 289 | Vvinifera | Vitis vinifera | dicot | phytozome12.1.6 |
| 38 | Zmarina | Zostera marina | monocot | phytozome12.1.6 |
| 151 | Zmays | Zea mays | monocot | phytozome12.1.6 |
| 134 | ZmaysPH207 | Zea mays PH207 | monocot | phytozome12.1.6 |
| 135 | Achinensis | Actinidia chinensis Red5 | dicot | plantensembl43 |
| 567 | Atauschii | Aegilops tauschii | monocot | plantensembl43 |
| 447 | Bnapus | Brassica napus | dicot | plantensembl43 |
| 312 | Boleraceae | Brassica oleracea | dicot | plantensembl43 |
| 122 | Bvulgaris | Beta vulgaris | dicot | plantensembl43 |
| 127 | Ccapsularis | Corchorus capsularis | dicot | plantensembl43 |

| | | | | |
|---|---|---|---|---|
| 59 | Ccrispus | Chondrus crispus | Rhodophyta | plantensembl43 |
| 161 | Drotundata | Dioscorea rotundata | monocot | plantensembl43 |
| 369 | Lperrieri | Leersia perrieri | monocot | plantensembl43 |
| 163 | Nattenuata | Nicotiana attenuata | dicot | plantensembl43 |
| 442 | Obar0thii | Oryza barthii | monocot | plantensembl43 |
| 267 | Obranchyantha | Oryza brachyantha | monocot | plantensembl43 |
| 297 | Oglaberrima | Oryza glaberrima | monocot | plantensembl43 |
| 410 | Oglumipatula | Oryza glumipatula | monocot | plantensembl43 |
| 601 | Olongistaminata | Oryza longistaminata | monocot | plantensembl43 |
| 189 | Omeridionalis | Oryza meridionalis | monocot | plantensembl43 |
| 427 | Onivara | Oryza nivara | monocot | plantensembl43 |
| 310 | Opunctata | Oryza punctata | monocot | plantensembl43 |
| 447 | Orufipogon | Oryza rufipogon | monocot | plantensembl43 |
| 1096 | Tdicoccoides | Triticum dicoccoides | monocot | plantensembl43 |
| 502 | Turartu | Triticum urartu | monocot | plantensembl43 |
| 163 | Vangularis | Vigna angularis | dicot | plantensembl43 |
| 83 | Vradiata | Vigna radiata | dicot | plantensembl43 |
| 1 | Ugibba | Utricularia gibba | dicot | CoGe |
| 17 | Gaurea | Genlisea aurea | dicot | CoGe |

Table S2

| Species | Genome size | No of Primary transcripts | No of NLRs with 6 motifs (HC NLRs) | Fraction of primary transcripts that are HC NLRs | Number of Low + High Confidence NLRs | Fraction of primary transcripts that are NLRs | Number NLRs with NB-ARC detected in genome by NLRannotator |
|---|---|---|---|---|---|---|---|
| Pineapple | 316 Mb | 27024 | 130 | 0.481 | 189 | 0.699 | 204 |
| *Amborella trich* | 706 Mb | 26846 | 54 | 0.201 | 109 | 0.406 | 77 |
| *Aquilegia coeru* | 306.5Mb | 30023 | 191 | 0.636 | 243 | 0.809 | 190 |
| *Amaranthus* | 377 Mb | 23054 | 80 | 0.347 | 130 | 0.564 | 162 |
| Chinese lotus | 803Mb | 38191 | 192 | 0.503 | 237 | 0.621 | 234 |
| Ash | 867.5 Mb | 38949 | 155 | 0.398 | 161 | 0.413 | 145 |
| Corkscrew plan | 63.3Mb | 17685 | 8 | 0.045 | 17 | 0.096 | 17 |
| Monkey flower | 312.7 Mb | 28140 | 280 | 0.995 | 326 | 1.158 | 261 |
| Tomato | ~900Mb | 34725 | 166 | 0.478 | 263 | 0.757 | 174 |
| Humped bladde | 99Mb | 31511 | 0 | 0.000 | 1 | 0.003 | 33 |
| Maize | 2,106 Mb | 39474 | 115 | 0.291 | 155 | 0.393 | 109 |
| Oil palm | 1.535 Gb | 41887 | 226 | 0.540 | 269 | 0.642 | 168 |
| Orchid | 980Mb | 29894 | 44 | 0.147 | 54 | 0.181 | 145 |
| Rice | 372Mb | 42099 | 381 | 0.905 | 516 | 1.226 | 419 |
| Ressurection gr | 244 Mb | 28354 | 7 | 0.025 | 68 | 0.240 | 142 |
| Eelgrass | 202.3 Mb | 20450 | 38 | 0.186 | 45 | 0.220 | 39 |
| Duckweed softmasked | | | | | | | 64 |
| Duckweed hardmasked | | | | | | | 46 |
| Duckweed_290_ | 142Mb | 19623 | 61 | 0.311 | 85 | 0.433 | 64 |
| *Arabidopsis* | 135Mb | 27416 | 156 | 0.569 | 167 | 0.609 | 141 |

Table S3

| Species | Complete BUSCO | Complete and single-copy BUSCO | Complete duplicated BUSCO | Fragmented BUSCO | Missing BUSCO | Total BUSCOs searched |
|---|---|---|---|---|---|---|
| Pineapple | 1264 | 1043 | 221 | 92 | 84 | 1440 |
| *Amborella trich* | 1196 | 1029 | 167 | 121 | 123 | 1440 |
| *Aquilegia coeru* | 1370 | 695 | 675 | 675 | 27 | 1440 |
| *Amaranthus* | 1209 | 1016 | 193 | 128 | 103 | 1440 |
| Chinese lotus | 1402 | 625 | 777 | 21 | 17 | 1440 |
| Ash | 1313 | 640 | 673 | 81 | 46 | 1440 |
| Corkscrew plan | 961 | 869 | 92 | 191 | 288 | 1440 |
| Monkey flower | 1347 | 1104 | 243 | 41 | 52 | 1440 |
| Tomato | 1368 | 1190 | 178 | 48 | 24 | 1440 |
| *Selagniella moe* | 860 | 664 | 196 | 182 | 398 | 1440 |
| Humped bladde | 950 | 818 | 132 | 136 | 354 | 1440 |
| Maize | 1359 | 1113 | 246 | 49 | 32 | 1440 |
| Oil palm | 1401 | 636 | 765 | 30 | 9 | 1440 |
| Orchid | 1271 | 658 | 613 | 74 | 95 | 1440 |
| Rice | 1373 | 905 | 468 | 39 | 28 | 1440 |
| Ressurection gr | 998 | 870 | 128 | 194 | 248 | 1440 |
| Eelgrass | 1212 | 1048 | 164 | 82 | 146 | 1440 |
| Yam | 1076 | 926 | 150 | 116 | 248 | 1440 |
| Duckweed | 1134 | 996 | 138 | 178 | 128 | 1440 |

| Arabidopsis GID for BUSCO validated as lost in Zmarina |
|---|
| morphogenesis GO:0010103 14857 P developmental processes IMP analysis of |
| morphogenesis GO:0010103 14857 P developmental processes RCA manually |
| stomatal complex development GO:2000122 35956 P developmental processes IGI |

Table S3

| Species | Query | No. | % protein coding genes | Query | No. | % protein coding genes | Query | No. | % protein coding genes | Total protein coding genes |
|---|---|---|---|---|---|---|---|---|---|---|
| *Amborella trich* | 'kinase' + 'LRR | 125 | 0.466 | kinase | 732 | 2.727 | Actin | 16 | 0.060 | 26846 |
| *Spirodella poly* | 'kinase' + 'LRR | 184 | 0.938 | kinase | 886 | 4.515 | Actin | 15 | 0.076 | 19623 |
| *Zoestera marin* | 'kinase' + 'LRR | 171 | 0.836 | kinase | 823 | 4.024 | Actin | 23 | 0.112 | 20450 |
| *Lemna minor* | 'kinase' + 'LRR | 125 | 0.559 | kinase | 757 | 3.383 | Actin | 17 | 0.076 | 22375 |
| Orchid | 'kinase' + 'LRR | 226 | 0.756 | kinase | 1268 | 4.242 | Actin | 28 | 0.094 | 29894 |
| Oil Palm | 'kinase' + 'LRR | 403 | 0.962 | kinase | 2087 | 4.982 | Actin | 56 | 0.134 | 41887 |
| *A coerulea* | 'kinase' + 'LRR | 313 | 1.043 | kinase | 1771 | 5.899 | Actin | 29 | 0.097 | 30023 |
| Chinese Lotus | 'kinase' + 'LRR | 398 | 1.042 | kinase | 1878 | 4.917 | Actin | 22 | 0.058 | 38191 |
| *Utricularia gib* | 'kinase' + 'LRR | 97 | 0.308 | kinase | 326 | 1.035 | Actin | 22 | 0.070 | 31511 |
| *Genlisea aurea* | 'kinase' + 'LRR | 1 | 0.006 | kinase | 585 | 3.308 | Actin | 19 | 0.107 | 17685 |
| Ash | 'kinase' + 'LRR | 448 | 1.150 | kinase | 2487 | 6.385 | Actin | 51 | 0.131 | 38949 |
| Monkey flower | 'kinase' + 'LRR | 262 | 0.931 | kinase | 1235 | 4.389 | Actin | 24 | 0.085 | 28140 |

Table S4

| Gene ID | Name | Description |
|---|---|---|
| AT5G13160 | PBS1 | avrPphB Susceptible 1 |
| AT3G25070 | RIN4 | RPM1-interacting protein 4 |
| AR3G55450 | PBL1 | PBS1-Like 1 |
| AT1G07570/ A | PBL9/10 | PBS1-Like 9 / PBS1-Like 10 |
| AT4G01370 | MPK4 | MAP Kinase 4 |
| AT3G48090/AT | EDS1 | Enhanced Disease Susceptiblity 1 |
| AT3G52430 | PAD4 | Phytoalexin deficient 4 |
| AT5G14930 | SAG101 | Senesence-associated gene 101 |
| AT4G33300 | ADR1-L1 | ADR1-Like 1 |
| AT2G43820 | SGT1 | Salicyclic acid glucosyltransferase 1 |
| AT1G74710/ A | ICS1/ICS2 | Isochorismate synthase 1 / sochorismate synthase 2 |
| AT1G19250 | FMO1 | Flavin-dependent monooxygenase 1 |
| AT2G13810 | ALD1 | AGD2-like defense response protein 1 |
| AT1G64280/ A | NPR1/NPR2 | Non-inducible immunity 1 / Non-inducible immunity 2 |
| AT1G02170 | MC1 | Metacaspase 1 |
| AT1G79340/ A | MC4/MC5 | Metacaspase 4/ Metacaspase 5 |
| AT1G43700 | VIP1 | VIRE2-interacting protein 1 |
| AT3G20600 | NDR1 | Non race-specific disease resistance 1 |
| AT5G51700 | RAR1 | Required for MLA12 Resistance 1 |

Table S5

| F00931 | Species_abb | Species | Taxon | EDS1 | AD4 | SAG101 | ADR1-L1 | NDR1 | RIN4 | RAR1 |
|---|---|---|---|---|---|---|---|---|---|---|
| 135 | Achinensis | Actinidia chir | dicot | P | P | P | P | P | P | P |
| 567 | Atauschii | Aegilops taus | monocot | P | P | A | P | A | P | P |
| 119 | Ahypochondr | Amaranthus | dicot | P | P | A | P | P | P | P |
| 99 | Atrichopoda | Amborella tr | Amborellales | P | A | P | P | P | P | P |
| 152 | Acomosus | Ananas como | monocot | P | P | A | P | A | P | P |
| 223 | Acoerulea | Aquilegia coe | dicot | P | A | P | P | P | P | P |
| 124 | Ahalleri | Arabidopsis h | dicot | P | P | P | P | P | P | P |
| 177 | Alyrata | Arabidopsis l | dicot | P | P | P | P | P | P | P |
| 160 | Athaliana | Arabidopsis t | dicot | P | P | P | P | P | P | P |
| 35 | Aofficinalis_ | Asparagus of | monocot | A | A | A | P | A | P | P |
| 122 | Bvulgaris | Beta vulgaris | dicot | P | P | A | P | P | P | P |
| 297 | Bstricta | Boechera str | dicot | P | P | P | P | P | P | P |
| 16 | Bbraunii_v2. | Botryococcus | Chlorophyta | A | X | A | A | A | A | A |
| 331 | Bdistachyon | Brachypodiur | monocot | P | P | A | P | A | P | P |
| 235 | Bstacei | Brachypodiur | monocot | P | P | A | P | A | P | P |
| 447 | Bnapus | Brassica nap | dicot | P | P | P | P | P | P | P |
| 114 | Boleraceacap | Brassica oler | dicot | P | P | P | P | P | P | A |
| 187 | BrapaFPsc | Brassica rapa | dicot | P | P | P | P | P | P | P |
| 96 | Cgrandiflora | Capsella grar | dicot | P | P | P | P | P | P | P |
| 116 | Crubella | Capsella rube | dicot | P | P | P | P | P | P | P |
| 47 | Cpapaya | Carica papay | dicot | P | P | P | P | P | P | P |
| 305 | Cquinoa_v1.0 | Chenopodiun | dicot | P | P | A | P | P | P | P |
| 59 | Ccrispus | Chondrus cris | Rhodophyta | A | A | A | A | A | A | A |
| 6 | Czofingiensis | Chromochlor | Chlorophyta | X | X | A | A | A | A | A |
| 342 | Cclementina | Citrus clemer | dicot | P | P | P | P | P | P | P |
| 441 | Csinensis | Citrus sinens | dicot | P | P | P | P | P | P | P |
| 127 | Ccapsularis | Corchorus ca | dicot | P | T | P | P | A | P | P |
| 58 | Csativus | Cucumis sati | dicot | P | P | P | P | P | P | P |
| 137 | Dcarota | Daucus carot | dicot | P | P | P | P | P | P | T |
| 161 | Drotundata | Dioscorea rot | monocot | P | P | A | P | P | A | P |
| 683 | Egrandis | Eucalyptus gr | dicot | P | T | P | P | P | P | P |
| 119 | Esalsugineur | Eutrema sals | dicot | P | P | P | P | P | P | P |
| 165 | Fvesca | Fragaria vesc | dicot | P | P | P | P | P | T | P |
| 423 | Gmax | Glycine max | dicot | P | P | P | P | P | P | P |
| 272 | Graimondii | Gossypium ra | dicot | P | P | P | P | P | P | P |
| 293 | Hannuus_r1. | Helianthus a | dicot | P | P | P | P | T | P | T |
| 404 | Hvulgare_r1 | Hordeum vul | monocot | P | P | A | P | A | P | P |
| 130 | Kfedtschenko | Kalanchoe fe | dicot | P | P | P | P | P | P | P |
| 182 | Klaxiflora | Kalanchoe la | dicot | P | P | P | P | P | P | P |
| 370 | Lsativa_v5 | Lactuca sativ | dicot | P | P | P | P | P | P | P |
| 369 | Lperrieri | Leersia perri | monocot | P | P | A | P | A | P | P |
| 157 | Lusitatissimu | Linum usitati | dicot | P | P | P | P | P | P | P |
| 930 | Mdomestica | Malus domes | dicot | P | P | P | P | P | T | P |
| 232 | Mesculenta | Manihot escu | dicot | P | P | P | P | P | P | P |
| 28 | Mpolymorph | Marchantia p | liverwort | A | A | A | A | P | A | P |
| 762 | Mtruncatula | Medicago tru | dicot | P | P | P | P | P | P | P |
| 296 | Mguttatus | Mimulus gut | dicot | P | P | A | P | P | P | P |
| 788 | Msinensis_v | Miscanthus s | monocot | P | P | A | P | A | P | P |
| 95 | Macuminata | Musa acumir | monocot | P | P | P | P | T | P | P |
| | | Nelumbo nuc | dicot | P | P | P | P | P | P | P |
| 163 | Nattenuata | Nicotiana att | dicot | P | P | P | P | P | P | P |
| 276 | Oeuropaea_v | Olea europae | dicot | P | P | P | P | P | P | P |
| 33 | Othomaeum | Oropetium th | monocot | P | P | A | P | A | P | P |
| 442 | Obarthii | Oryza barthii | monocot | P | P | A | P | A | P | P |
| 267 | Obranchyant | Oryza brachy | monocot | P | P | A | P | A | P | P |
| 297 | Oglaberrima | Oryza glaber | monocot | P | T | A | P | A | P | P |
| 410 | Oglumipatula | Oryza glumip | monocot | P | P | A | P | A | P | P |
| 601 | Olongistamin | Oryza longist | monocot | P | P | A | P | A | P | P |
| 189 | Omeridionali | Oryza meridi | monocot | T | P | A | A | A | P | P |
| 427 | Onivara | Oryza nivara | monocot | P | P | A | P | A | P | P |
| 310 | Opunctata | Oryza puncta | monocot | P | P | A | P | A | P | P |
| 447 | Orufipogon | Oryza rufipog | monocot | P | P | A | P | A | P | P |
| 445 | Osativa | Oryza sativa | monocot | P | P | A | P | A | P | P |
| 211 | PhalliiHAL_v2 | Panicum hall | monocot | P | P | A | P | A | P | P |
| 1013 | Pvirgatum | Panicum virg | monocot | P | P | A | P | A | P | P |
| 310 | Pvulgaris | Phaseolus vu | dicot | P | P | P | P | P | P | P |
| 120 | Ppatens | Physcomitrel | moss | A | A | A | A | A | P | P |
| 510 | Ptrichocarpa | Populus trich | dicot | P | P | P | P | P | P | P |
| 46 | Pumbilicalis_ | Porphyra um | Rhodophyta | A | A | A | A | A | A | A |
| 355 | Ppersica | Prunus persic | dicot | P | P | P | P | P | P | P |
| 140 | Rcommunis | Ricinus comr | dicot | P | P | P | P | T | P | P |
| 365 | Spurpurea | Salix purpure | dicot | P | P | P | P | P | P | P |
| 17 | Smoellendor | Selaginella n | moss | X | A | A | A | X | P | P |
| 396 | Sitalica | Setaria italic | monocot | P | P | A | P | A | P | P |
| 327 | Sviridis | Setaria viridi | monocot | P | P | A | P | A | P | P |
| 235 | Slycopersicur | Solanum lyco | dicot | P | P | P | P | P | P | P |
| 398 | Stuberosum | Solanum tub | dicot | P | P | P | P | P | P | P |
| 302 | Sbicolor | Sorghum bic | monocot | P | P | A | P | A | P | P |
| 216 | Sfallax | Sphagnum fa | moss | A | A | A | A | A | P | P |
| 70 | Spolyrhiza | Spirodela pol | monocot | A | A | A | A | A | P | P |
| 254 | Tcacao | Theobroma c | dicot | P | P | P | P | P | P | P |
| 388 | Tpratense | Trifolium pra | dicot | P | P | P | P | P | P | P |
| 1157 | Taestivum_v | Triticum aest | monocot | P | P | A | P | A | P | P |
| 1096 | Tdicoccoides | Triticum dicc | monocot | P | P | A | P | A | P | P |

| | |
|---|---|
| P | BLASTp |
| T | tBLASTn |
| A | Absemt |
| X | PARTIAL |

| | | | | | | | | | | |
|---|---|---|---|---|---|---|---|---|---|---|
| 502 | Turartu | Triticum urar | monocot | P | P | A | P | A | P | P |
| 83 | Vradiata | Vigna radiata | dicot | P | P | P | P | A | P | P |
| 289 | Vvinifera | Vitis vinifera | dicot | P | P | P | P | T | P | P |
| 151 | Zmays | Zea mays | monocot | P | P | A | P | A | P | P |
| 38 | Zmarina | Zostera mari | monocot | A | A | A | A | A | P | P |
| 237 | Fraxinus exce | Fraxinus exce | dicot | P | P | P | P | P | P | P |
| 17 | Genlisea aur | Genlisea aur | dicot | A | A | A | A | A | P | P |
| 326 | Mimulus gut | Mimulus gut | dicot | P | P | A | P | P | P | P |
| 1 | Utricularia gi | Utricularia gi | dicot | A | A | A | A | A | P | P |
| 54 | Phalaenopsis | Phalaenopsis | monocot | P | P | A | P | P | P | P |
| 269 | Elaeis guinee | Elaeis guinee | monocot | P | P | A | P | P | P | P |

Core Asterids

| Order | Genus | Family/Species | Code | AT3G48080<br>EDS1<br>AT3G48080 | AT3G52430<br>PAD4<br>AT3G52430 | AT5G14930<br>SAG101<br>AT5G14930 | AT4G33300<br>ADR1-L1<br>AT4G33300 | AT3G20600<br>NDR1<br>AT3G20600 | AT3G25070<br>RIN4<br>AT3G25070 | AT5G51700<br>RAR1<br>AT5G51700 |
|---|---|---|---|---|---|---|---|---|---|---|
| Lamiales | Oleaceae | Chionanthus retusus | KTAR | | | | | | | |
| Lamiales | Oleaceae | Ligustrum sinense | MZLD | | | | | | | |
| Lamiales | Tetrachondra | Polypremum procumbens | COBX | | | ■ | | | | |
| Lamiales | Gesneriacea | Saintpaulia ionantha | RWKR | | | ■ | | | | |
| Lamiales | Gesneriacea | Sinningia tuberosa | DTNC | | | ■ | | | | |
| Lamiales | Calceolariac | Calceolaria pinifolia | DCCI | | | ■ | | | ■ | |
| Lamiales | Plantaginace | Plantago virginica | PTBJ | | | ■ | | | | |
| Lamiales | Plantaginace | Digitalis purpurea | GNRI | | | ■ | | | | |
| Lamiales | Plantaginace | Antirrhinum majus | EBOL | | | ■ | | | ■ | |
| Lamiales | Plantaginace | Bacopa caroliniana | CLRW | | | ■ | | | | |
| Lamiales | Scrophularia | Anticharis glandulosa | EJBY | | | ■ | | | | |
| Lamiales | Scrophularia | Buddleja sp. | GRFT | | | ■ | | | | |
| Lamiales | Scrophularia | Celsia arcturus | SIBR | | | ■ | | | | |
| Lamiales | Byblidaceae | Byblis gigantea | GDZS | | | ■ | | | ■ | |
| Lamiales | Acanthaceae | Anisacanthus quadrifidus | PCGJ | | | ■ | | | | |
| Lamiales | Acanthaceae | Ruellia brittoniana | AYIY | | | ■ | | | | |
| Lamiales | Acanthaceae | Sanchezia sp. | NBMW | | | ■ | | | ■ | ■ |
| Lamiales | Bignoniacea | Kigelia africana | QKEI | | | ■ | | | | |
| Lamiales | Lentibulariac | Utricularia sp. | HRUR | ■ | ■ | ■ | ■ | ■ | | |
| Lamiales | Lentibulariac | Pinguicula caudata | JCMU | ■ | ■ | ■ | ■ | ■ | ■ | |
| Lamiales | Lentibulariac | Pinguicula agnata | MXFG | ■ | ■ | ■ | ■ | ■ | | |
| Lamiales | Schlegeliace | Schlegelia parasitica | GAKQ | | | ■ | | | | |
| Lamiales | Schlegeliace | Schlegelia violacea | EDXZ | | | ■ | | | | |
| Lamiales | Schlegeliace | Schlegelia parasitica | CWLL | | | | | | | |
| Lamiales | Verbenaceae | Lantana camara | | ■ | | ■ | | | | |
| Lamiales | Verbenaceae | Verbena hastata | GCFE | | | ■ | | | | |
| Lamiales | Verbenaceae | Phyla dulcis | MQIV | | | ■ | | | | |
| Lamiales | Lamiaceae | Scutellaria montana | ATYL | | | ■ | | | | |
| Lamiales | Lamiaceae | Solenostemon scutellarioides | BAHE | | | ■ | | | ■ | |
| Lamiales | Lamiaceae | Vitex agnus-castus | DMLT | | | | | | | |
| Lamiales | Lamiaceae | Marrubium vulgare | EAAA | | | ■ | | | | |
| Lamiales | Lamiaceae | Salvia spp. | EQDA | | | ■ | | | | |
| Lamiales | Lamiaceae | Rosmarinus officinalis | FDMM | | | ■ | | | | |
| Lamiales | Lamiaceae | Nepeta cataria | FUMQ | | | ■ | | | | |
| Lamiales | Lamiaceae | Lavandula angustifolia | FYUH | | ■ | ■ | | | ■ | ■ |
| Lamiales | Lamiaceae | Pogostemon sp. | GETL | | | ■ | | ■ | | |
| Lamiales | Lamiaceae | Oxera neriifolia | GNPX | | | ■ | | | | |
| Lamiales | Lamiaceae | Thymus vulgaris | IYDF | | | ■ | | | | |
| Lamiales | Lamiaceae | Teucrium chamaedrys | LRRR | | | ■ | | | | |
| Lamiales | Lamiaceae | Prunella vulgaris | PHCE | | | ■ | | | | |
| Lamiales | Lamiaceae | Leonurus japonicus | SNNC | | | ■ | | | | |
| Lamiales | Lamiaceae | Agastache rugosa | PUCW | | | ■ | | | | |
| Lamiales | Lamiaceae | Oxera pulchella | RTNA | | | ■ | | | | |
| Lamiales | Orobanchace | Orobanche fasciculata | PHOQ | ■ | ■ | ■ | ■ | ■ | | |
| Lamiales | Orobanchace | Lindenbergia philippensis | ZVFS | | | | | | | |
| Lamiales | Orobanchace | Lindenbergia philippensis | WUZV | | | | | | | |
| Lamiales | Orobanchace | Conopholis americana | FAMO | | | ■ | | ■ | | |
| Lamiales | Rehmnniace | Rehmannia glutinosa | OWAS | | | | | | | |

Present

Absent in transcriptome

| Order | Genus | Family/Species | Code | | AT3G48080 EDS1 | AT3G52430 PAD4 | AT5G14930 SAG101 | AT4G33300 ADR1-L1 | AT3G20600 NDR1 | AT3G25070 RIN4 | AT5G51700 RAR1 |
|---|---|---|---|---|---|---|---|---|---|---|---|
| Acorales | Acoraceae | Acorus americanus | MTII | Wetland | | | | | | | |
| Alismatales | Alismatacea | Sagittaria latifolia | DWZT | Shallow wetland | | | | | | | |
| Alismatales | Araceae | Spirodela polyrhiza | | | | | | | | | |
| Alismatales | Araceae | Lemna minor | | | | | | | | | |
| Alismatales | Araceae | Pistia stratiotes | MFIN | | | | | | | | |
| Alismatales | Juncaginacea | Triglochin maritima | COCP | Marshes | | | | | | | |
| Alismatales | Zosteraceae | Zostera marina | | | | | | | | | |
| Alismatales | Posidoniacea | Posidonia australis | BYQM | Seagrass | | | ????????? | | | | |
| Alismatales | Araceae | Typhonium blumei | YMES | Grassland/Between rocks - Rhizomes | | | | | | | |
| Dioscoreales | Dioscoreacea | Dioscorea villosa | OCWZ | | | | | | | | |
| Pandanales | Cyclanthacea | Ludovia sp. | VVVV | | | | | | | | |
| Pandanales | Pandanaceae | Freycinetia multiflora | DGXS | | | | | | | | |
| Pandanales | Stemonaceae | Stemona tuberosa | VBHQ | | | | | | | | |
| Pandanales | Velloziaceae | Talbotia elegans | SILJ | | | | | | | | |
| Pandanales | Velloziaceae | Xerophyta villosa | QOXT | | | | | | | | |
| Asparagales | Agavaceae | Agave tequilana | KXSK | | | | | | | | |
| Asparagales | Agavaceae | Chlorogalum pomeridianum | PLBZ | | | | | | | | |
| Asparagales | Agavaceae | Hesperaloe parviflora | CMCY | | | | | | | | |
| Asparagales | Agavaceae | Yucca brevifolia | YBML | | | | | | | | |
| Asparagales | Agavaceae | Yucca filamentosa | ICNN | | | | | | | | |
| Asparagales | Asparagacea | Asparagus densiflorus | FGRF | | | | | NRG1 | | | |
| Asparagales | Asparagacea | Disporopsis pernyi | | | | | | | | | |
| Asparagales | Asparagacea | Maianthemum canadense | | | | | | | | | |
| Asparagales | Asparagacea | Maianthemum sp. | RCUX | | | | | | | | |
| Asparagales | Hemerocallic | Hemerocallis spp. | BLAJ | | | | | | | | |
| Asparagales | Rusaceae | Nolina atopocarpa | | | | | | | | | |
| Asparagales | Orchidaceae | Oncidium sphacelatum | CNTZ | | | | | | | | |
| Asparagales | Amaryllidace | Phycella aff. cyrtanthoides | DMIN | | | | | | | | |
| Asparagales | Amaryllidace | Zephyranthes treatiae | DPFW | | | | | | | | |
| Asparagales | Boryaceae | Borya sphaerocephala | EMJJ | | | | | | | | |
| Asparagales | Hemerocallic | Phormium tenax | FCEL | | | | | | | | |
| Asparagales | Alliaceae | Allium sativum | GJPF | | | | | | | | |
| Asparagales | Ruscaceae | Nolina atopocarpa | HOKG | | | | | | | | |
| Asparagales | Amaryllidace | Narcissus viridiflorus | IQYY | | | | | | | | |
| | Themidaceae | Brodiaea sierrae | IXEM | | | | | | | | |
| | Amaryllidace | Rhodophiala pratensis | JDYT | | | | | | | | |
| | Hemerocallic | Hemerocallis sp. | JHUL | | | | | | | | |
| | Asphodelace | Aloe vera | JVBR | | | | | | | | |
| | Alliaceae | Allium commutatum | KBXS | | | | | | | | |
| | Hyacinthacea | Urginea maritima | KOFB | | | | | | | | |
| | Tecophilaeac | Cyanella orchidiformis | KYNE | | | | | | | | |
| | Amaryllidace | Amaryllis belladonna | LDME | | | | | | | | |
| | Orchidaceae | Haemaria discolor | LELS | | | | | | | | |
| | Orchidaceae | Oncidium sphacelatum | | | | | | | | | |
| | Orchidaceae | Masdevallia yuangensis | JSAG | | | | | | | | |
| | Ruscaceae | Ruscus sp. | LSJW | | | | | | | | |
| | Iridaceae | Sisyrinchium angustifolium | LTZF | | | | | | | | |
| | Orchidaceae | Platanthera clavellata | MTHW | | | | | | | | |
| | Laxmanniace | Lomandra longifolia | MUMD | | | | | | | | |
| | Ruscaceae | Sansevieria trifasciata | MVRF | | | | | | | | |
| | Ruscaceae | Eriospermum lancifolia | ONBE | | | | | | | | |
| | Agapanthace | Agapanthus africanus | PRFO | | | 2 EDS1 | | | | | |
| | Tecophilaeac | Cyanastrum cordifolium | RDYY | | | | | | | | |
| | Ruscaceae | Nolina bigelovii | RQZP | | | | | | | | |
| | Xeronematac | Xeronema callistemon | SART | | | | | | | | |
| | Hyacinthacea | Drimia altissima | SVTS | | | | | | | | |
| | Ruscaceae | Peliosanthes minor | TCYS | | | | | | | | |
| | Orchidaceae | Vanilla planifolia | THDM | NA | | | | | | | |
| | Asparagacea | Disporopsis pernyi | UZXL | | | | | | | | |
| | Orchidaceae | Goodyera pubescens | VTUS | young leaves | | | | | | | |
| | Xanthorrhoe | Johnsonia pubescens | WTDE | flowers | | | | | | | |
| | Amaryllidace | Narcissus viridiflorus | XEUV | young leaf | | | | | | | |
| | Asparagacea | Maianthemum canadense | XFG | | | | | | | | |
| | Orchidaceae | Drakaea elastica | XZME | leaves, flowers, and buds | | | | | | | |
| | Hypoxidacea | Curculigo sp. | YJUG | | | | | | | | |
| | Amaryllidace | Traubia modesta | ZPFK | Sepals | | | | | | | |
| Liliales | Melanthiace | Xerophyllum asphodeloides | AFLV | | | | | | | | |
| Liliales | Colchicaceae | Gloriosa superba | GDKK | | | | | | | | |
| | Smilacaceae | Smilax bona-nox | MWYQ | | | | | | | | |
| | Melanthiace | Helonias bullata | OOSO | | | | | | | | |
| | Colchicaceae | Colchicum autumnale | OVRB | | | | | | | | |
| | Colchicaceae | Colchicum autumnale | NHIX | | | | | | | | |
| | Colchicaceae | Colchicum autumnale | SFCT | | | | | | | | |
| | Liliaceae | Lilium sargentiae | THEW | woodland/grassland | | | | | | | |
| Laurales | Lauraceae | Sassafras albidum | ABSS | | | | | | | | |
| | Lauraceae | Cinnamomum camphora | BCGB | | | | | | | | |
| | Hernandiace | Gyrocarpus americanus | BSVG | | | | | | | | |

Present
Absent in transcriptome
Absent in genome

| | | | | |
|---|---|---|---|---|
| | Saururaceae | Houttuynia cordata | CSSK | |
| | Canellaceae | Canella winterana | DDEV | |
| Magnoliales | Eupomatiace | Eupomatia bennettii | DHPO | |
| Laurales | Calycanthace | Calycanthus floridus | FALI | |
| | Canellaceae | Canella winterana | IFCJ | |
| | Lauraceae | Lindera benzoin | KAYP | Shrub mainland US |
| | Monimiacea | Peumus boldus | KRJP | |
| Laurales | Gomortegac | Gomortega keule | MAQO | |
| | Piperaceae | Piper auritum | MUNP | |
| | Myristicacea | Myristica fragrans | OBPL | |
| | Saururaceae | Saururus cernuus | OPDF | |
| | Aristolochiac | Aristolochia elegans | PAWA | |
| Magnoliales | Annonaceae | Uvaria microcarpa | PSJT | |
| | Aristolochiac | Saruma henryi | QDVW | |
| | Lauraceae | Cassytha filiformis | VYLQ | |
| Laurales | Atherosperm | Laurelia sempervirens | WAIL | |
| | Magnoliacea | Magnolia grandiflora | WBOD | |
| | Lauraceae | Persea borbonia | WIGA | |
| | Winteraceae | Drimys winteri | WKSU | |
| | Calycanthac | Idiospermum australiense | WPHN | |
| | Magnoliacea | Michelia maudiae | XQWC | |
| Piperales | Piperaceae | Peperomia fraseri | XSZI | Tropical soil |
| | Annonaceae | Annona muricata | YZRI | |

AT3G20590 present
AT3G20610 present

Table S8

| Orthogroup | Arabidopsis GID |
|---|---|
| Dicot_10831- monocots_10960 | AT4G02075 |
| Dicot_10831-monocots_10960 | AT1G02610 |
| Dicot_8212-monocots_6587 | AT1G27920 |
| Galaxy group 3847 | AT5G56060 |
| Galaxy group 3847 | AT5G56075 |
| Galaxy group 3847 | AT5G25030 |
| Galaxy group 3847 | AT1G55790 |
| Galaxy group 3847 | AT4G26485 |
| Galaxy group 3847 | AT1G55800 |
| Galaxy group 29536 | AT1G79760 |
| Galaxy group 1912 | AT4G33930 |
| Galaxy group 1912 | AT2G15770 |
| Galaxy group 1912 | AT4G34300 |
| Galaxy group 1912 | AT2G15780 |
| Dicot_3622-monocots_9489 | AT2G19930 |
| Dicot_3622-monocots_9489 | AT2G19920 |
| Dicot_3622-monocots_9489 | AT2G19910 |
| Galaxy group-654 | AT2G30890 |
| Dicot_8689-monocots_8858 | AT3G50400 |
| Dicot_8689-monocots_8858 | AT2G23540 |
| Galaxy group 26335 | AT2G37870 |
| Galaxy group 26335 | AT3G53980 |
| Galaxy group 26335 | AT5G05960 |
| Galaxy group 15377 | AT3G08490 |
| Dicot_10092-monocots_5982 | AT1G53650 |
| Dicot_10092-monocots_5982 | AT3G14450 |
| Dicot_6524-monocots_3737 | AT3G14470 |
| Dicot_8371-monocots_7573 | AT3G20015 |
| Dicot_3663A-monocots_1124 | AT2G47120 |
| Dicot_3663A-monocots_1124 | AT2G47130 |
| Dicot_3663A--monocots_1124 | AT2G47140 |
| Dicot_3663B-monocots-4965 | AT3G29250 |
| Dicot_3663B-monocots_4965 | AT3G29260 |
| Galaxy group 16353 | AT3G48080 |
| Galaxy group 16353 | AT3G48090 |
| Galaxy group 16353 | AT5G14930 |
| Galaxy group 16353 | AT3G52430 |
| Dicot_10185-monocots_9077 | AT3G63060 |
| Galaxy group 8237 | AT4G02170 |
| Dicot_9707-monocots_6672 | AT5G10080 |
| Galaxy group 5398 - Dicot_7418· | AT5G11580 |
| Dicot_1692-monocots_3327 | AT5G44380 |
| Dicot_1692-monocots_3327 | AT5G44390 |
| Dicot_1692-monocots_3327 | AT5G44400 |
| Galaxy group 4516 | AT5G66890 |
| Galaxy group 4516 | AT3G26470 |
| Galaxy group 4516 | AT5G66900 |
| Galaxy group 4516 | AT5G66910 |
| Galaxy group 4516 | AT5G04720 |
| Galaxy group 4516 | AT5G47280 |
| Galaxy group 4516 | AT1G33560 |
| Galaxy group 4516 | AT4G33300 |

ASTREL gene known to be part of the plant immune pathway

ASTREL gene not previously implicated in the plant immune pathway

Table S9

| Orthogroup | Rice ASTRAL GID |
|---|---|
| Dicot_10831- monocots_10960 | LOC_Os01g19800 |
| Dicot_8212-monocots_6587 | LOC_Os08g41890 |
| Galaxy group 3847 | LOC-Os09g30210 |
| Galaxy group 3847 | LOC-Os09g30200 |
| Galaxy group 29536 | LOC_Os01g54090 |
| Galaxy 19121 | LOC_Os06g11310 |
| Dicot_3622-monocots_9489 | LOC_Os01g10130 |
| Dicot_3622-monocots_9489 | LOC_Os01g10140 |
| Galaxy-654 | LOC_Os01g47635 |
| Galaxy-654 | LOC_Os05g49040 |
| Dicot_8689-monocots_8858 | LOC_Os07g47210 |
| Galaxy group 15377 | LOC-Os03g11710 |
| Dicot_10092-monocots_5982 | LOC_Os01g11120 |
| Dicot_6524-monocots_3737 | LOC_Os05g41310 |
| Dicot_6524-monocots_3737 | LOC_Os10g36270 |
| Dicot_6524-monocots_3737 | LOC_Os05g41290 |
| Dicot_8371-monocots_7573 | LOC_Os04g58070 |
| Dicot_3663B-monocots_4965 | LOC_Os03g61740 |
| Galaxy group 16353 | LOC-Os11g09010 |
| Galaxy group 16353 | LOC-Os09g22450 |
| Dicot_10185-monocots_9077 | LOC_Os01g58850 |
| Galaxy group 8237 | LOC-Os03g19070 |
| Galaxy group 8237 | LOC-Os01g13050 |
| Galaxy group 8237 | LOC-Os11g04580 |
| Galaxy group 8237 | LOC-Os01g64230 |
| Galaxy group 8237 | LOC-Os03g24880 |
| Galaxy group 26335 | LOC-Os04g33930 |
| Galaxy group 26335 | LOC-Os04g33920 |
| Galaxy group 26335 | LOC-Os05g06780 |
| Galaxy group 26335 | LOC-Os01g62980 |
| Dicot_9707-monocots_6672 | LOC_Os02g51540 |
| Dicot_7418-monocots_8406 | LOC_Os01g52630 |
| Dicot_1692-monocots_3327 | LOC_Os06g35590 |
| Dicot_1692-monocots_3327 | LOC_Os06g35630 |
| Dicot_1692-monocots_3327 | LOC_Os06g35650 |
| Dicot_1692-monocots_3327 | LOC_Os06g35700 |
| Dicot_1692-monocots_3327 | LOC_Os11g30310 |
| Dicot_1692-monocots_3327 | LOC_Os06g35560 |
| Dicot_1692-monocots_3327 | LOC_Os06g35660 |

ASTREL gene known to be part of the plant immune pathway

ASTREL gene not implicated in the plant immune pathway

| | |
|---|---|
| Galaxy group 4516 | LOC-Os02g10150 |

Table S8

| Arabidopsis ASTRAL Genes | Other Names | Aquatic P/A | Asparagus P/A | Oryza sativa |
|---|---|---|---|---|
| dicot_10831- monocots_10960-AT4G02075 | | | | |
| dicot_10831-monocots_10960-AT1G02610 | | | | |
| dicot_8212-monocots_6587-AT1G27920 | | | | |
| Galaxy group 3847- AT5G56060 | | | | |
| Galaxy group 3847- AT5G56075 | | | | |
| Galaxy group 3847- AT5G25030 | | | | |
| Galaxy group 3847 - AT1G55790 | | | | |
| Galaxy group 3847 -AT4G26485 | | | | |
| Galaxy group 3847 AT1G55800 | | | | |
| Galaxy-29536-AT1G79760 | | | | |
| Galaxy 19121-AT4G33930 | | | | |
| Galaxy 19121-AT2G15770 | | | | |
| Galaxy 19121-AT4G34300 | | | | |
| Galaxy 19121-AT2G15780 | | | | |
| dicot_3622-monocots_9489-AT2G19930 | | | | |
| dicot_3622-monocots_9489-AT2G19920 | | | | |
| dicot_3622-monocots_9489-AT2G19910 | | | | |
| Galaxy group 654-AT2G30890 | | | | |
| dicot_8689-monocots_8858-AT3G50400 | | | | |
| dicot_8689-monocots_8858-AT2G23540 | | | | |
| Galaxy group 26335-AT2G37870 | | | | |
| Galaxy group 26335- AT3G53980 | | | | |
| Galaxy group 26335 - AT5G05960 | | | | |
| Galaxy group 15377 - AT3G08490 | | | | |
| dicot_10092-monocots_5982-AT1G53650 | | | | |
| dicot_10092-monocots_5982-AT3G14450 | | | | |
| dicot_6524-monocots_3737-AT3G14470 | | | | |
| dicot_8371-monocots_7573-AT3G20015 | ASPG2 | | | |
| dicot_3663A-monocots_1124-AT2G47120 | | | | |
| dicot_3663A-monocots_1124-AT2G47130 | | | | |
| dicot_3663A--monocots_1124AT2G47140 | | | | |
| dicot_3663B-monocots-4965-AT3G29250 | SDR4 | | | |
| dicot_3663B-monocots_4965-AT3G29260 | SDR5 | | | |
| Galaxy group 16353 - AT3G48080 | EDS1 | | | |
| Galaxy group 16353 - AT3G48090 | EDS1 | | | |
| Galaxy group 16353 - AT5G14930 | SAG101 | | | |
| Galaxy group 16353 - AT3G52430 | PAD4 | | | |
| dicot_10185-monocots_9077-AT3G63060 | EDL3 | | | |
| Galaxy group 8237- AT4G02170 | | | | Rice lineage expansion |
| dicot_9707-monocots_6672-AT5G10080 | | | | |
| dicot_7418-monocots_8406-AT5G11580 | | | | |
| dicot_1692-monocots_3327-AT5G44380 | | | | |
| dicot_1692-monocots_3327-AT5G44390 | | | | |
| dicot_1692-monocots_3327-AT5G44400 | | | | |
| Galaxy group 4516 - AT5G66890 | | | | |
| Galaxy group 4516 - AT3G26470 | | | | |
| Galaxy group 4516 - AT5G66900 | NRG1 | | | |
| Galaxy group 4516 - AT5G66910 | NRG1.2 | | | |
| Galaxy group 4516 - AT5G04720 | | | | |
| Galaxy group 4516 -AT5G47280 | | | | |
| Galaxy group 4516 - AT1G33560 | ADR1 | | | |
| Galaxy group 4516 -AT4G33300 | ADR1-L1 | | | |

- Absent
- Present
- Lineage specific expansion

| Orthogroup | Arabidopsis ( | Rice GID |
|---|---|---|
| Dicot_8371-mo | AT3G20015 | LOC_Os04g58070 |
| Dicot_3663B-n | AT3G29260 | LOC_Os03g61740 |
| Dicot_3663B-n | AT3G29250 | |
| Galaxy group 1 | AT3G48080 | LOC-Os11g09010 |
| Galaxy group 1 | AT3G48090 | LOC-Os09g22450 |
| Galaxy group 1 | AT5G14930 | |
| Galaxy group 1 | AT3G52430 | |
| Dicot_10185-n | AT3G63060 | LOC_Os01g58850 |
| Galaxy group | AT5G66890 | LOC-Os02g10150 |
| Galaxy group | AT3G26470 | |
| Galaxy group | AT5G66900 | |
| Galaxy group | AT5G66910 | |
| Galaxy group | AT5G04720 | |
| Galaxy group | AT5G47280 | |
| Galaxy group | AT1G33560 | |
| Galaxy group | AT4G33300 | |

Table S11

| Gene ID | Domain | Publication references – Drought upregulation |
|---|---|---|
| AT2G37870 | Lipid transfer | [illegible] of Abscisic Acid Signaling by a Plant Immune Response Pathway." Current Biology : CB, vol. 21, no. 11, 2011, pp. 990-997, doi:10.1016/j.cub.2011.04.045 |
| | | Allu, A. D., et al. "Salt Stress and Senescence: Identification of Cross-Talk Regulatory Components." Journal of Experimental Botany, vol. 65, no. 14, 2014, pp. 3993-4008, doi:10.1093/jxb/eru173 |
| | | Seki, Motoaki, et al. "Transcriptome Analysis in Abiotic Stress Conditions in Higher Plants." Plant Responses to Abiotic Stress. Edited by Heribert Hirt, and Kazuo Shinozaki. Springer Berlin Heidelberg, Berlin, Heidelberg, 2004, https://doi.org/10.1007/978-3-540-39402-0_11, doi:10.1007/978-3-540-39402-0_11. |
| | | Zandalinas Sara. Plant Strategies to Deal with a Combination of Drought and High Temperatures & nbsp;, Universitat Jaume, 2016. |
| AT2G15780 | Cupredoxin family | |
| | | Fàbregas, Norma, et al. "Overexpression of the Vascular Brassinosteroid Receptor BRL3 Confers Drought Resistance without Penalizing Plant Growth." Nature Communications, vol. 9, no. 1, 2018, pp. 4680. PubMed, https://www.ncbi.nlm.nih.gov/pubmed/30409967 https://www.ncbi.nlm.nih.gov/pmc/PMC6224425/, doi:10.1038/s41467-018-06861-3 |
| | | Zander, Mark. Arabidopsis Thaliana Class II TGA Transcription Factors Provide a Molecular Link between Salicyclic Acid and Ethylene Defence Signalling, Georg-August-Universität Göttingen, 2011 |
| | Microbial | Endo, Satoshi, Kuninori Iwamoto, and Hiroo Fukuda. "Overexpression and Cosuppression of Xylem-Related Genes in an Early Xylem Differentiation Stage-Specific Manner by the AtTED4 Promoter." Plant Biotechnology Journal, vol. 16, no. 2, 2018, pp. 451-458, https://doi.org/10.1111/pbi.12784, doi:10.1111/pbi.12784. |
| | | Ko, Jae-Heung, Eric P. Beers, and Kyung-Hwan Han. "Global Comparative Transcriptome Analysis Identifies Gene Network Regulating Secondary Xylem Development in Arabidopsis Thaliana." Molecular Genetics and Genomics, vol. 276, no. 6, 2006, pp. 517-531, https://doi.org/10.1007/s00438-006-0157-1, doi:10.1007/s00438-006-0157-1. |

| | | |
|---|---|---|
| MAP65-8 | Microbial tubule associated protein | Yamaguchi, Masatoshi, et al. "VASCULAR-RELATED NAC-DOMAIN 7 Directly Regulates the Expression of a Broad Range of Genes for Xylem Vessel Formation." The Plant Journal, vol. 66, no. 4, 2011, pp. 579-590, https://doi.org/10.1111/j.1365-313X.2011.04514.x, doi:10.1111/j.1365-313X.2011.04514.x. |
| | | Guo, Longbiao, et al. "Evaluating the Microtubule Cytoskeleton and its Interacting Proteins in Monocots by Mining the Rice Genome." Annals of Botany, vol. 103, no. 3, 2009, pp. 387-402. PubMed, https://www.ncbi.nlm.nih.gov/pubmed/19106179 https://www.ncbi.nlm.nih.gov/pmc/PMC2707338/, doi:10.1093/aob/mcn248. |
| | | Keech, O., et al. "Leaf Senescence is Accompanied by an Early Disruption of the Microtubule Network in Arabidopsis." Plant Physiology, vol. 154, no. 4, 2010, pp. 1710-1720, doi:10.1104/pp.110.163402 |
| AT4G02170 | Cotton fiber | Bergmann, Dominique C., Wolfgang Lukowitz, and Chris R. Somerville. "Stomatal Development and Pattern Controlled by a MAPKK Kinase." Science, vol. 304, no. 5676, 2004, pp. 1494-1497, http://science.sciencemag.org/content/304/5676/1494.abstract, doi:10.1126/science.1096014. |
| | | Sakuraba, Yasuhito, et al. "The Arabidopsis Transcription Factor NAC016 Promotes Drought Stress Responses by Repressing AREB1 Transcription through a Trifurcate Feed-Forward Regulatory Loop Involving NAP." The Plant Cell, vol. 27, no. 6, 2015, pp. 1771–1787. JSTOR, www.jstor.org/stable/plantcell.27.6.1771. |
| AT3G53980 | Cotton fibre protein | Wenzel, C. L., Q. Hester, and J. Mattsson. "Identification of Genes Expressed in Vascular Tissues using NPA-Induced Vascular Overgrowth in Arabidopsis." Plant & Cell Physiology, vol. 49, no. 3, 2008, pp. 457-468, doi:10.1093/pcp/pcn023 |
| | | Zandalinas Sara. Plant Strategies to Deal with a Combination of Drought and High Temperatures & nbsp;, Universitat Jaume, 2016. |
| AT5G04760 | AtDIV2 | Song, L., et al. "A Transcription Factor Hierarchy Defines an Environmental Stress Response Network." *Science (New York, N.Y.)*, vol. 354, no. 6312, 2016, pp. aag1550. doi: 10.1126/science.aag1550, doi:aag1550 |
| | *AtMYBL-O* | Jeong, Chan Y., et al. "Overexpression of Abiotic Stress-Induced AtMYBL-O Results in Negative Modulation of Abscisic Acid Signaling through the Downregulation of Abscisic Acid-Responsive Genes in Arabidopsis Thaliana." *Plant Growth Regulation*, vol. 84, no. 1, 2018, pp. 25-36, https://doi.org/10.1007/s10725-017-0318-8, doi:10.1007/s10725-017-0318-8. |

| | | |
|---|---|---|
| | | Jeong, Chan Y., et al. "AtMybL-O Modulates Abscisic Acid Biosynthesis to Optimize Plant Growth and ABA Signaling in Response to Drought Stress." *Applied Biological Chemistry*, vol. 61, no. 4, 2018b, pp. 473-477, https://doi.org/10.1007/s13765-018-0376-2, doi:10.1007/s13765-018-0376-2. |
| AT5G44400 | | Bauer, Hubert, et al. The Stomatal Response to Reduced Relative Humidity Requires Guard Cell-Autonomous ABA Synthesis. vol. 23, , 2013, http://www.sciencedirect.com/science/article/pii/S0960982212013310, doi://doi.org/10.1016/j.cub.2012.11.022. |
| | | Catala, Rafael, et al. "The Arabidopsis E3 SUMO Ligase SIZ1 Regulates Plant Growth and Drought Responses." The Plant Cell, vol. 19, no. 9, 2007, pp. 2952-2966, http://www.plantcell.org/content/19/9/2952.abstract, doi:10.1105/tpc.106.049981. |
| | | Vie, Ane K., et al. "NEVERSHED and INFLORESCENCE DEFICIENT IN ABSCISSION are Differentially Required for Cell Expansion and Cell Separation during Floral Organ Abscission in Arabidopsis Thaliana." Journal of Experimental Botany, vol. 64, no. 17, 2013, pp. 5345-5357, https://doi.org/10.1093/jxb/ert232, doi:10.1093/jxb/ert232. |
| | | Zhu, Zhangsheng, et al. "Overexpression of AtEDT1/HDG11 in Chinese Kale (Brassica Oleracea Var. Alboglabra) Enhances Drought and Osmotic Stress Tolerance." Frontiers in Plant Science, vol. 7, 2016, pp. 1285, https://www.frontiersin.org/article/10.3389/fpls.2016.01285. |

Table S10

| Gene ID | Name | Expression +Patho | eds1 DE | pad4 DE | npr1 DE | wrk33 DE | Literature | References |
|---|---|---|---|---|---|---|---|---|
| AT1G27920 | MAP65-8 | DOWN | | | | | Down regulated upon dark induced senesence | [illegible] Gutierrez L, et al. Leaf senescence is accompanied by an early disruption of the microtubule network in Arabidopsis. Plant |
| AT1G33560 | ADR1 | UP | | | | | | |
| AT1G55790 | HIPP41 | UP | X | X | X | X Opp | Ungrouped among other HIPPs as has lost conserved first residue | Tehseen, Muhammad, et al. "Metallochaperone-Like Genes in Arabidopsis Thaliana." Metallomics, vol. 2, no. 8, 2010, pp. 556-564, http://dx.doi.org/10.1039 |
| AT1G79760 | DTA4 | DOWN | | | | | Downstrea m of senesence and abscission master regulator AGL15 | Tang, W., and S. E. Perry. "Binding Site Selection for the Plant MADS Domain Protein AGL15: An in Vitro and in Vivo Study." The Journal of Biological Chemistry, vol. 278, no. 30, 2003, pp. |
| AT2G15780 | - | UP | X | X | X | | ABA & Patho overlapping genes | "Chemical Genetics Reveals Negative Regulation of Abscisic Acid Signaling by a Plant Immune Response Pathway." Current Biology : CB, vol. 21, no. 11, 2011, pp. 990-997, |
| AT2G30890 | - | DOWN | X | | | | [illegible] ferric reductase, predicted accept electron, though missing first motif maybe subfunction alised | [illegible] Fusako Takeuchi, and Nobuyuki Nakanishi. Cytochrome b561 Protein Family: Expanding Roles and Versatile Transmembrane Electron Transfer Abilities as Predicted by a New Classification System and Protein Sequence Motif Analyses. vol. 1753, , |

| | | | | | | | | |
|---|---|---|---|---|---|---|---|---|
| AT2G37870 | - | UP | X | | | | ABA sensence induced genes | [illegible] Stress and Senescence: Identification of Cross-Talk Regulatory Components." Journal of Experimental Botany, vol. 65, no. 14, 2014, pp. |
| AT3G14450 | CID9 | DOWN | X | | | | Downstream SR45 a supressor of plant immunity and involved in post transcriptio | Zhang, Xiao-Ning, et al. "Transcriptome Analyses Reveal SR45 to be a Neutral Splicing Regulator and a Suppressor of Innate Immunity in Arabidopsis Thaliana." BMC |
| AT3G14470 | RPPL1 | UP | X | X | X | X Opp | Hub NLR like | [illegible] al. "Genome-Wide Functional Analyses of Plant Coiled-Coil NLR-Type Pathogen Receptors Reveal Essential Roles of their N-Terminal Domain in Oligomerization, Networking, and Immunity." PLoS Biology, vol. 16, no. 12, 2018, pp. e2005821. PubMed, |
| AT3G20015 | ASPG2 | DOWN | | | | | | Hachez, Charles, et al. "Differentiation of Arabidopsis Guard Cells: Analysis of the Networks Incorporating the Basic Helix-Loop-Helix Transcription Factor, FAMA." *Plant Physiology*, vol. 155, no. 3, 2011, pp. 1458-1472, http://www.plantphysiol.org/content/155/3/1458.abstract, doi:10.1104/pp.110.167718. |
| AT3G26470 | RPW8 | UP | | | | | EDS1 DEPENDENT | Wang, Wenming, et al. "Specific Targeting of the Arabidopsis Resistance Protein RPW8.2 to the Interfacial Membrane Encasing the Fungal Haustorium Renders Broad-Spectrum Resistance to Powdery Mildew." The Plant Cell, vol. 21, no. 9, 2009, pp. 2898-2913. PubMed, https://www.ncbi.nlm.nih.gov/pubmed/19749153 |

| | | | | | | | | |
|---|---|---|---|---|---|---|---|---|
| AT3G48090 | EDS1 | UP | | | | | | |
| AT3G52430 | PAD4 | UP | | | | | | |
| AT3G63060 | EDL3 | UP | | | | | Positive regulator ABA signallng by inducing proteosomal degredation of ABA repressors | Bohnert. "Gene Networks in Arabidopsis Thaliana for Metabolic and Environmental Functions." *Molecular BioSystems*, vol. 4, no. 3, 2008, pp. 199-204, http://dx.doi.org/10.1039/B715811B, doi:10.1039/B715811B. |
| AT4G33300 | ADR1-L1 | UP | | | | | | |
| AT5G04720 | ADR1-L2 | UP | | | | | | |
| AT5G04760 | AtMYBL-C | UP | | | | X Opp | Negative regulator of ABA | "Overexpression of Abiotic Stress-Induced AtMYBL-O Results in Negative Modulation of Abscisic Acid Signaling through the Downregulation of Abscisic Acid-Responsive Genes in Arabidopsis Thaliana." *Plant Growth Regulation*, vol. 84, |
| AT5G10080 | - | DOWN | | | | | Stunts growth when OE | Pogorelko, Gennady V., et al. A New Technique for Activation Tagging in Arabidopsis. vol. 414, , 2008, http://www.sciencedirect.com/science/article/pii/S0378111908000784, |
| AT5G11580 | - | DOWN | X | | | X Opp | | |
| AT5G14930 | SAG101 | UP | | | | | | |
| AT5G56060 | - | UP | | | | | | |
| AT5G66890 | NRG1C | UP | X | X | X | X Opp | Prior analysis suggest truncated non-functional | Wu, Zhongshou, et al. "Differential Regulation of TNL-Mediated Immune Signaling by Redundant Helper CNLs." *New Phytologist*, vol. 0, no. 0, |
| AT5G66900 | NRG1 | UP | | | | | | |
| AT5G66910 | NRG1-L1 | UP | | | | | | |

Aquatic specific
